# A chemically reactive and Raman-active non-canonical amino acid reveals photocycle complexity in a blue-light receptor

**DOI:** 10.64898/2026.05.26.727966

**Authors:** Aditi Chatterjee, Phuong Ngoc Pham, Atripan Mukherjee, Petra Čubaková, Spyridon Kaziannis, Jakub Dostal, Miroslav Kloz, Aditya S. Chaudhari, Giusy Finocchiaro, Tomáš Špringer, Jiří Homola, Ondřej Honc, Martin Kizovský, Martin Šafařík, Jan Štursa, Lukáš Werner, Jaroslav Šebestík, Gustavo Fuertes

## Abstract

Photosensory protein function spans multiple time and length scales, demanding integrative approaches. We introduce 4-diacetylenyl-phenylalanine (DAF), a dual-purpose non-canonical amino acid (ncAA) that enables both chemical control and spectroscopic readout of photoreceptor dynamics. Genetically encoded in *E. coli*, DAF combines a reactive diyne for bioorthogonal ligations (thiols, azides, tetrazines) with a strong, solvatochromic Raman signal in the cell-silent region. Applied to the light-oxygen-voltage (LOV) transcription factor EL222, DAF enables multifaceted interrogation of its photocycle. We engineer a covalently cross-linked variant that suppresses light-driven conformational changes and DNA binding, and generate a donor–acceptor construct for Förster resonance energy transfer (FRET) tracking of photoinduced structural dynamics. Time-resolved stimulated Raman spectroscopy following flavin mononucleotide (FMN) excitation reveals additional processes -from vibrational energy transfer to local unfolding-spanning femtoseconds to milliseconds. DAF thus constitutes a versatile tool to resolve protein dynamics with high spatiotemporal resolution.

## Introduction

Highly evolved proteins and their associated small-molecule cofactors convert light absorption into metabolic changes that include alterations in gene expression, cell differentiation, molecular trafficking and cell motility.^1^ The photocontrolled transcription factor EL222 from the marine bacterium *Erythrobacter litoralis* binds DNA and promotes gene expression when blue-light is absorbed by the embedded flavin mononucleotide (FMN) chromophore.^2^ The overall photocycle of EL222, shared among the members of the light-oxygen-voltage (LOV) family,^3^ is well known, comprising at a minimum three intermediate species: singlet (1FMN*), triplet (3FMN*) and adduct (A390) states.^3–7^ The first two refer to electronic states of the FMN cofactor, while the latter indicates covalent bonding between the FMN and the protein chain via a conserved cysteine residue. Recent evidence leveraging on non-canonical amino acids (ncAA) and improved experimental set-ups, suggests that such a chromophore-centric view is an over-simplification and additional intermediates are likely present.^8–12^

The ability to incorporate ncAA into proteins offers the possibility to understand and probe protein structure and activity, and also to design and evolve proteins with new functions.^13,14^ Among the many methods for introducing ncAA into proteins, genetic code expansion (GCE) allows researchers to install one, or a few copies, of a given ncAA in a site-specific fashion concomitant with protein synthesis.^15^ Unnatural side-chains are being steadily added to the portfolio of ncAA yet the large majority are developed with a singlepurpose in mind, for instance as conjugation handles to attach a myriad of other groups,^16^ or as phototriggers to start out-of-equilibrium protein dynamics.^17,18^ A notable exception are ncAA bearing azido groups, like 4-azido-phenylalanine, which are chemically reactive against alkynes, photochemically reactive, and give rise to strong absorption bands in the “transparent window” infrared spectral region.^19–21^ A major benefit of using ncAA as reporters^21–27^ and modulators^28–30^ of photoreceptor dynamics is that they enable protein engineering with atomic precision.

Diynes, and particularly conjugated diynes (-C≡C-C≡C-)_n_ having an alternating ene-yne backbone, are attractive due to their unique structure and properties.^31^ They have a rod-like molecular shape with high rigidness, and are readily attacked by nucleophiles such as amines and thiols. In addition, they vibrate in the cell-silent region (1800 cm^-1^ to 2700 cm^-1^), a feature that makes them popular as metabolic tracers for vibrational-based imaging.^32,33^ However, diynes are typically appended to sugars, lipids, nucleotides, drugs, and other small molecules.^34,35^ In a few cases, diynes have been attached to proteins post-translationally.^36,37^ Genetically encoded Raman-active probes are scarce and based on single alkynes: homopropargylglycine (HPG),^38,39^ propargyl-lysine (PRK),^36^ and 4-ethynyl-phenylalanine (ETF).^40,41^ The inherently weak Raman signal severely limits their practical utility.

To address the above limitations, here we report the synthesis and genetic incorporation in *E. coli* of 4-diacetylenyl-phenylalanine (DAF), an ncAA featuring the smallest possible conjugated diyne (diacetylene) attached to the aromatic ring (H-C≡C-C≡C-Ph) of a phenylalanine amino acid at the *para* position. On the one hand, the diyne moiety reacts selectively with thiols, azides, and tetrazines, both in solution and in a cellular environment. We exploit such a feature to prepare (i) an intramolecularly cross-linked EL222 variant locked in the inactive dark-state conformation, and (ii) a labeled EL222 protein bearing donor and acceptor dyes suitable for time-resolved Förster resonance energy transfer (FRET) studies of EL222 structural dynamics. On the other hand, the Raman intensity of DAF surpasses that of any other alkyne-containing ncAA, while keeping an excellent response to solvent polarity and H-bonding. EL222 decorated with DAF at several residue positions confirms the excellent sensitivity of its alkyne vibration to light-dependent changes in local protein microenvironment and reveals new dynamical events in the sub-millisecond range.

## Results

### Synthesis and characterization of the novel non-canonical amino acid DAF

The diyne-containing ncAA DAF (**Fig. 1A**) was synthesized in three steps from the commercially available *tert*-butoxycarbonyl (Boc)-protected 4-iodo-L-phenylalanine (see detailed synthesis procedures in **Supplementary Note S1**). First, *para*-iodophenylalanine was converted to *para*-trimethylsilylbutadiynyl derivative (2). The trimethylsilyl group was subsequently removed. Finally, cleavage of the Boc protecting group yielded the final product DAF as a trifluoroacetate salt in 80% yield (5).

**Fig. 1.**
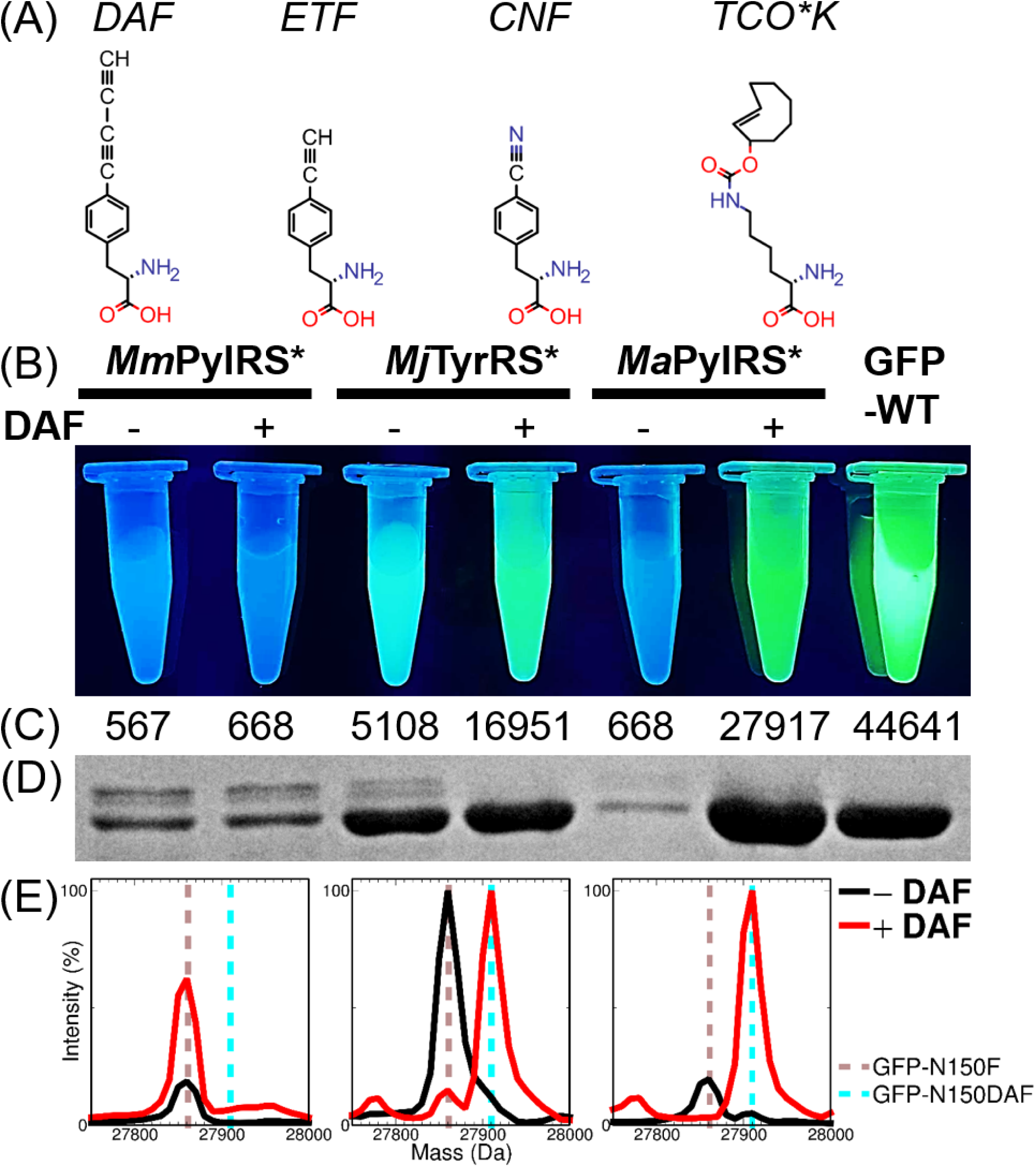
Genetic incorporation of DAF in *E. coli*. (A) Structure of the four main non-canonical amino acids discussed in this paper. DAF: 4-diacetylenyl-phenylalanine. ETF: 4-ethynyl-phenylalanine. CNF: 4-cyano-phenylalanine. TCO*K: trans-cyclooctene-lysine. (B) Images upon UV excitation of *E. coli* suspensions in microcentrifuge tubes co-expressing GFP-N150TAG and an orthogonal aminoacyl-tRNA synthetase/tRNA pair: *Mm*PylRS* (*left*) or *Mj*TyrRS* (*middle*) or *Ma*PylRS* (*right*, see mutations of all three synthetases in **Supplementary Note S4**) in the absence (-) and presence (+) of DAF as indicated. (C) Fluorescence emission intensities of the cultures shown in (B). (D) Corresponding Coomassie-Blue stained SDS-PAGE after purification of GFP-N150DAF by a C-terminal affinity handle. (E) Normalized mass spectra of the six purified proteins.

Next, we conducted a comprehensive exploration of the suitability of DAF both as a bioorthogonal reactive handle and as a Raman probe. Of biological relevance, diynes, as any other alkyne, show reactivity towards azides via copper-catalyzed alkyne-azide cycloaddition (CuAAC),^42^ and against thiols via thiol-yne coupling.^43^ In addition, we included two bio-orthogonal reactions typically employed in protein chemistry: strain-promoted alkyne-azide cycloaddition (SPAAC) and tetrazine ligation via inverse electron demand Diels Alder (IEDDA).^16^ DAF reactivity was assessed by a combination of mass spectrometry and fluorescence spectroscopy (**Supplementary Note S2, Table S1, Fig. S2, Fig. S3**). Taken together, the results summarized in **Fig. 2A**, suggest that DAF reacts with azides in the presence of copper(I) but not when the catalyst is absent. Interestingly, we found evidence of reactivity between the diyne of DAF and tetrazines, particularly H-tetrazines and to a lesser extent methyl-tetrazines, which to the best of our knowledge, has not been previously described in the literature. The related ncAA ETF, which contains a single aryl alkyne group, displays a similar reactivity pattern, although with somewhat reduced efficiency.

**Fig 2.**
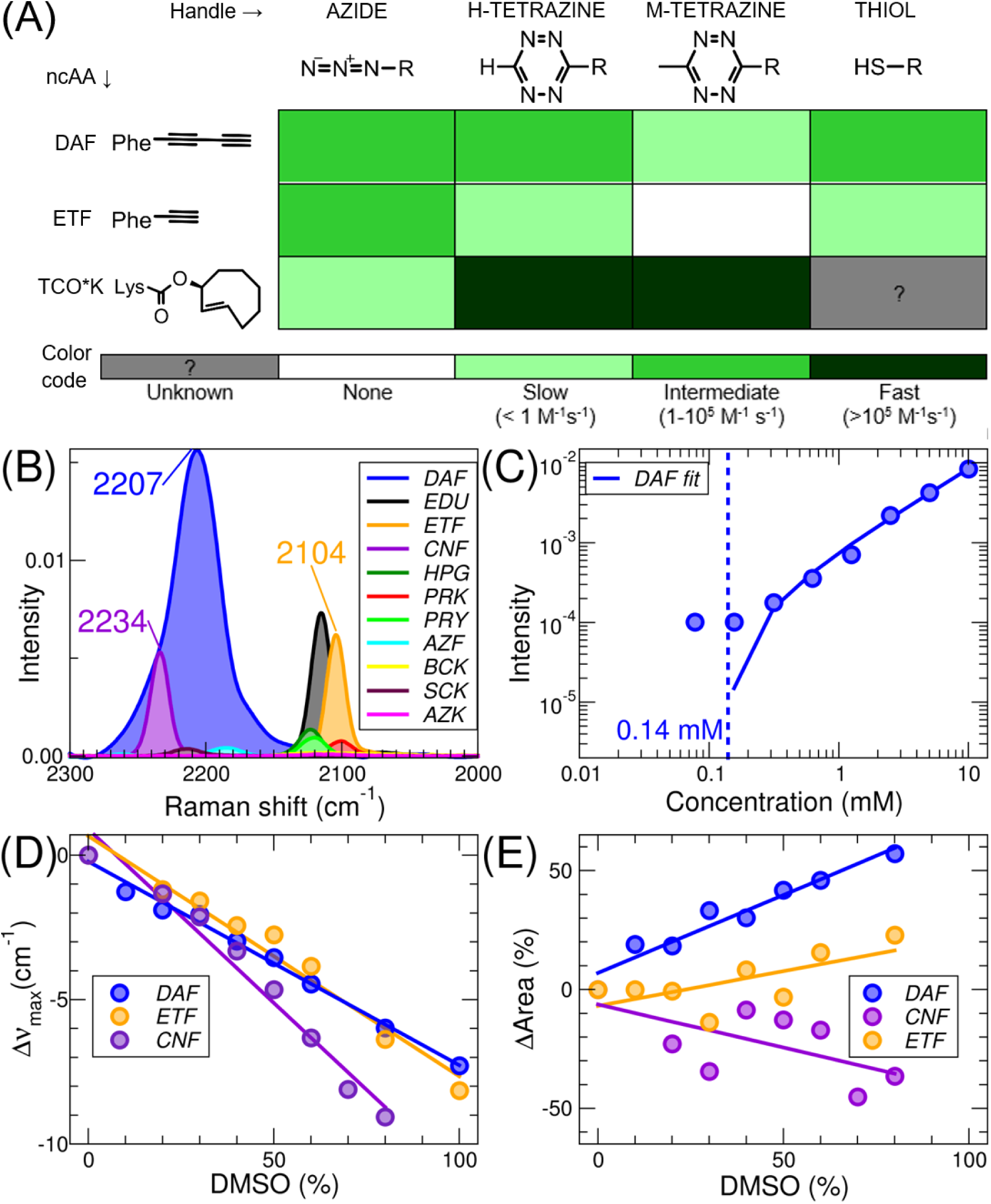
Reactivity and Raman activity of DAF. (A) Matrix of the reactivity between DAF (and other ncAA) and the indicated chemical handles. Each table cell is color coded (as indicated in the bottom bar) to convey the estimated second-order rate constant for the reaction of each pair of compounds. White boxes indicate that the reaction was not detected, and gray boxes indicate that the reactivity was not measured. (B) Bond-normalized femtosecond-stimulated Raman spectra (FSRS, λ_Raman pump_ = 800 nm, pulse energy = ∼4 μJ, same conditions in all panels) of DAF and related ncAA in different solvents (See supplementary **Table S3**). The peak maxima (in inverse centimeters) of DAF, ETF, and CNF are indicated. (C). Raman intensity of DAF peak at 2210 cm^-1^ in methanol solvent as a function of concentration. The dashed line indicates the limit of detection (∼0.14 mM). (D) Change in the position of the vibrational bands of the 3 indicated ncAA in aqueous solution as a function of DMSO content. (E) Change in the area of the vibrational bands of the 3 indicated ncAA as a function of DMSO.

The femtosecond-stimulated Raman spectra (FSRS) of DAF display an intense peak corresponding to the C≡C bond stretches (∼2207 cm^-1^) in the cellular silent region (**Fig. 2B**, see additional results in **Supplementary Note S3, Table S2, Fig. S5**). The Raman intensity of DAF is much higher than any other ncAA bearing a group vibrating in the “transparent window” region (**Fig. 2B**, see additional results in Supplementary Note S3, **Table S3**). The limit of detection of DAF in our FSRS setup was found to be ∼140 μM (**Fig. 2C**). Importantly, both the peak position (**Fig. 2D**) and peak area (**Fig. 2E**) of DAF depend on solvent composition. A similar behavior is observed with CNF, a commonly used vibrational reporter (**Fig. 2D** and **2E**). In contrast, the solvatochromic effect of ETF affects mainly the peak position rather than the area (**Fig. 2D** and **2E**).

In summary, the ncAA DAF combines two unique properties: multiple chemical reactivity (thiols, alkynes, and tetrazines), and Raman-activity in the cell-silent region (including frequency and intensity changes). We next looked for robust procedures to genetically encode DAF into proteins.

### Genetic encoding of DAF

Since DAF is essentially a phenylalanine modified at the *para* position, we hypothesized that previously engineered aminoacyl-tRNA synthetases (aaRS) could recognize it as a substrate. Specifically, we focused on two polyspecific aaRS: a tyrosyl-tRNA synthetase from *Methanocaldococcus jannaschii* (*Mj*TyRS*)^44^ and a pyrrolysyl-tRNA synthetase from *Methanosarcina mazei* (*Mm*PylRS*)^45^, (see specific mutations in **Supplementary Note S4**). We conducted stop codon suppression experiments using GFP as the target protein containing an amber codon at the permissive position 150 (see protein quality controls in **Supplementary Note S5**). In agreement with our expectations, we observed clear fluorescence upon DAF addition (1 mM) to *E. coli* cultures co-expressing GFP-N150TAG and *Mj*TyrRS*/tRNATyr_CUA_ (**Fig. 1B and 1C**). Moreover, the SDS-PAGE (**Fig. 1D**) and mass spectra (**Fig. 1E**) of the purified proteins confirmed the correct incorporation of DAF at the desired position. The absence of chemical modifications suggest that the intracellular concentration of endogenous thiols is not sufficient to efficiently react with DAF. However, DAF readily reacts with thiols at higher concentrations (> 10 mM, *vide infra*). Given the mediocre suppression efficiency (the protein yield of DAF-containing GFP variant was ∼40% relative to wild-type GFP) and fidelity (significant mis-incorporation of phenylalanine) of the *Mj*TyrRS*/*Mj*TyrtRNA pair as the orthogonal translation system (**Table S6**), we decided to evolve a DAF-specific aaRS (**Supplementary Note S6, Table S5**).^46^

Starting from a library of pyrrolysyl-tRNA synthetase from *Methanomethylophilus alvus* (*Ma*PylRS), a promising *Ma*PylRS mutant termed DafRS emerged after just one round of positive, negative, and fluorescence-based selections (**Fig. S7**). We leveraged again on GFP-N150TAG as a reporter and fluorescence (**Fig. 1C**)/SDS-PAGE (**Fig. 1D**)/mass spectrometry (**Fig. 1E**) assays to verify that the incorporation of DAF into the target protein had high efficiency (nearly 100 %) and fidelity (94 %, **Table S6**), indicative that the evolved *Ma*DafRS was specifically active for DAF but inactive for the canonical amino acids. Thus, we utilized the *Ma*DafRS/*Ma*PyltRNA_CUA_ pair for the site-specific encoding of DAF.

### Chemical reactivity of MBP-DAF

To understand and probe the versatility of DAF as a genetically encoded chemical handle for bio-orthogonal ligations (**Fig. 3A**), we performed labeling experiments with fluorescent dyes coupled to different reactive groups, first with purified proteins in solution and then in live *E. coli* cells.

**Fig. 3.**
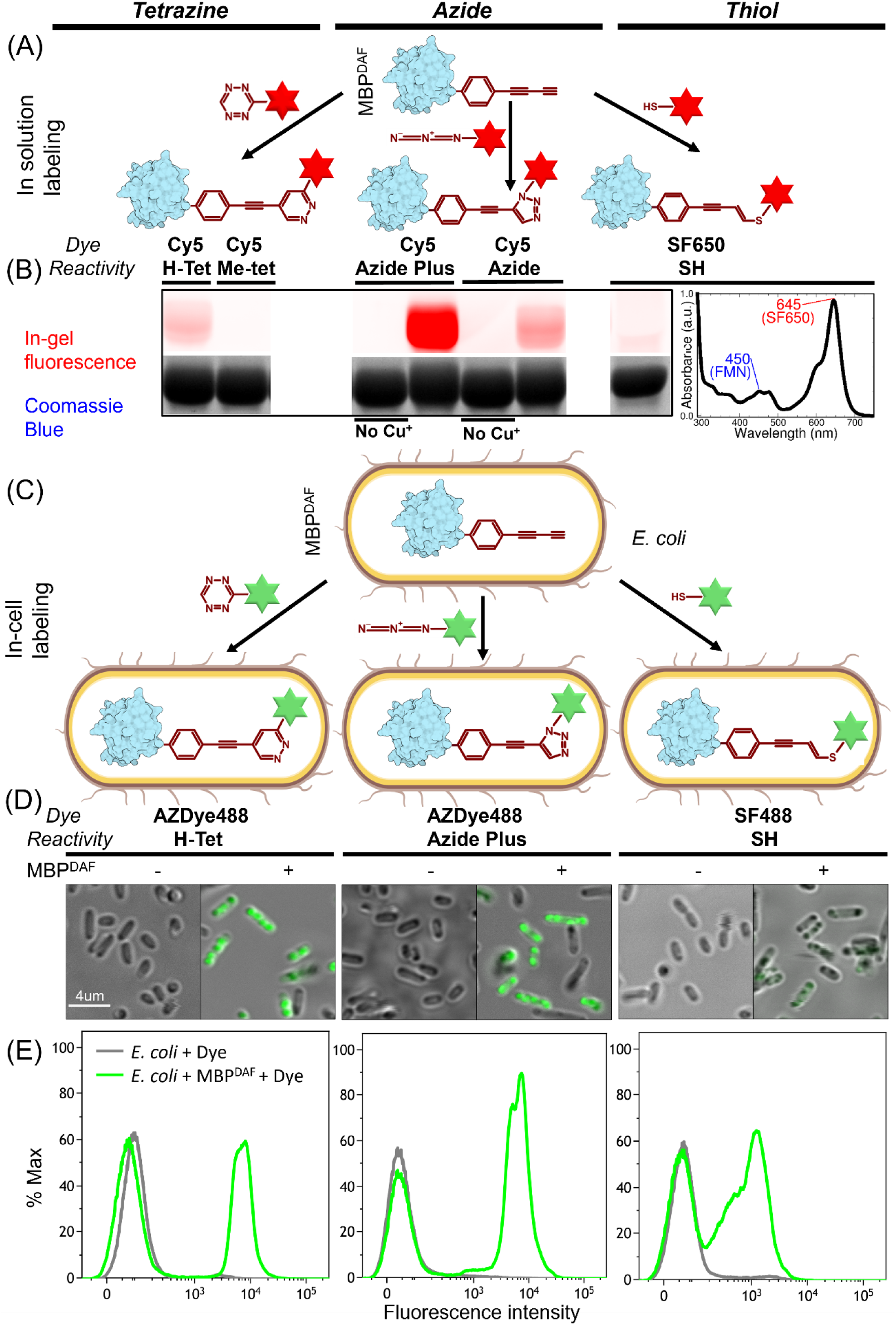
Labeling of DAF-containing proteins with fluorescent dyes in solution and *in cellulo*. (A) Scheme showing the reaction between purified MBP bearing DAF at residue 38 (MBP^DAF^, originally expressed in *E. coli*), and a red-fluorescing dye (Cy5 or SF650) carrying azide, tetrazine (H-tetrazine or Methyl-tetrazine), or thiol (SH) moieties. (B) Fluorescence image (λ_excitation_ = 650 nm, λ_emission_ ∼ 750 nm) and corresponding Coomassie Blue-stained SDS-PAGE of 30 µM MBP^DAF^ incubated with ten-fold excess of red-fluorescing dyes bearing distinct reactive handles for 22 hours. Azide reactions were done both in the absence and in the presence of copper (Cu^+^) as a catalyst to check for SPAAC and CUAAC reaction mechanisms, respectively. In the case of thiol-yne reaction, the absorbance spectrum of 1 mM purified EL222^DAF^ after incubation with four-fold excess of SF650-SH for 22 hours and removal of excess dye is shown on the right. The peak maxima in nanometer units of FMN and SF650 are indicated. (C) Scheme showing the reaction between maltose binding protein (MBP) bearing DAF at position 38 (MBP^DAF^) recombinantly expressed in *E. coli* BL21(DE3) and exogenously added green-fluorescing dyes (AZDye488 or SF488) containing azide, tetrazine or sulfhydryl groups. (D) Confocal fluorescence images (λ_excitation_ = 488 nm, λ_emission_ ∼575 nm) from of *E.coli* cells expressing MBP^DAF^ incubated with green fluorescent dyes (250 µM H-tetrazine, 100 µM AzidePlus and 50 µM thiol) for 30 minutes. The scale bar is indicated on the left panel. (E) Flow cytometry histograms showing the fluorescence intensity (λ_excitation_ = 488 nm, λ_emission_ ∼ 530 nm) distribution of cells within the gated *E.coli* population after the three labelling reactions depicted in (C). As a control to check the bio-orthogonality of the reactions, fluorescent dyes were also mixed with *E. coli* cells in the absence of DAF. The three tested reactions are shown in distinct sides of the panels: tetrazine-alkyne ligations (*left*), azide-alkyne cycloadditions (*middle*) and thiol-yne (*right*).

We introduced DAF at the residue position 38 of maltose binding protein (MBP). Purified MBP-V38DAF was incubated with red fluorescent dyes (Cy5 or the spectrally similar SF650) containing different thiols, azides (canonical azides and AzidePlus), and tetrazines (H-tetrazines and methyl-tetrazines) (**Fig. 3B**). The reactions were analyzed by SDS-PAGE followed by in-gel fluorescence and eventually stained by Coomassie Blue (**Fig. 3B**). Thiol labeling could not be determined unambiguously (even under non-reducing conditions). In these cases, DAF was introduced into the residue position 151 of EL222 and mixed with SF650-SH. After the incubation period, we removed the excess dye by gel filtration chromatography and recorded the UV/Visible spectrum (**Fig. 3B right**). The degree of thiol labeling was estimated to be 40 %. Additionally, the formation of adducts between DAF and the thiol-containing reducing agent DTT was assessed by mass spectrometry (**Supplementary Note S7**, **Fig. S8A**). Unexpectedly, slow reaction rates were observed between DAF and canonical azides. In contrast, the more reactive AzidePlus-containing dyes, which include a copper-chelating system in their structure, gave clear bands. Regarding tetrazines, H-tetrazines yielded abundant conjugates with MBP-V38DAF, while methyl-tetrazines produced virtually no bands. In parallel, we determined the reactivity pattern of ETF (**Supplementary Note S8**). Like DAF, ETF showed clear reactivity against AzidePlus and H-tetrazine dyes (**Fig. S9B**). However, no band was observed upon incubation with methyl-tetrazines, and the yield of thiol conjugates was significantly lower as judged by UV/Visible spectroscopy (**Fig. S9B right**) and mass spectrometry (**Fig. S8B**).

Then we tested DAF reactivity directly in living bacterial cells by fluorescent imaging and flow cytometry. *E. coli* cells expressing MBP-V38TAG were incubated first with DAF and later with green fluorescent dyes (AZDye488, **Fig. 3C**). Fluorescent images of *E. coli* cells suggested the accumulation of the dyes inside the bacteria only in the presence of DAF (**Fig. 3D**, see additional experiments in **Supplementary Note S11** and **Fig. S13**). The negative controls (MBP-WT) are explained in **Supplementary Note S9** (**Fig. S10**).To improve statistical robustness, the percentage of fluorescence-positive cells within the *E. coli* population was quantified by flow cytometry after washing out excess dye (**Fig. 3E**, see additional details in **Supplementary Note S10**). The higher percentage of fluorescence positive cells expressing MBP-V38DAF relative to the negative controls (**Table S7**) confirmed the feasibility of site-specific *in-cell* labeling with DAF. Both CuAAC between the double alkyne of DAF and AzidePlus probes, and tetrazine ligations between the alkyne of DAF and H-Tetrazines provided the best contrast due to the lowest fluorescent background (low non-specific binding and reactivity of dyes towards cellular components). In the case of thiol-yne reactions the difference between negative and positive controls were smaller.

A summary of the reactivity properties of DAF is included in **Supplementary Note S12** (**Table S8**). To illustrate the utility of DAF as a chemical handle while providing valuable information into EL222 photocycle, we concentrated on two reactions *in vitro*: thiol-yne and tetrazine ligation.

### Cross-linked EL222-DAF

Proximity-induced chemistry facilitates chemical transformations between a reactive group and a specific natural residue of proteins through the complex-induced proximity effect.^47,48^ Such a proximity-enabled thiol-yne click chemistry reaction could be leveraged to prepare an intramolecularly cross-linked EL222 variant unable to undergo large conformational transitions and thus essentially inactive. The lit-state structure(s) of EL222 is currently unknown at high resolution. However, according to our current understanding of EL222 photocycle, LOV-HTH interactions found in the dark should be replaced with LOV-LOV and HTH-HTH interactions. Thus, based on the known dark-state structure of EL222,^2,8^ we looked for residue pairs that could hold together the LOV and HTH domains. **Fig. 4A** shows a model of dark-state EL222 highlighting residues L35 (LOV) and V217 (HTH). CNF35 was previously shown to act as a reporter of LOV-HTH interactions.^9^ Thus, we designed, expressed and purified an EL222 double mutant: EL222-L35DAF-V217C. MS confirmed the presence of a cross-link involving residues 35 and 217 (**Supplementary Note S13, Fig. S14**). To check the light-induced conformational changes of this mutant, referred to as XL-EL222 for simplicity, we measured the protein response in the structure-sensitive Amide I’ band by infrared (IR) spectroscopy.^49^ Wild-type (WT) EL222 undergoes blue-light-induced unfolding of α-helices (**Fig. 4B**), most likely involving A’α at the N-terminus and/or the linking Jα element, as previously shown.^9^ Both XL-EL222 mutant and WT-EL222 show changes in the carbonyl groups of FMN associated to adduct formation between the FMN and Cys78. However, little spectral changes around ∼1650 cm^-1^ are observed for XL-EL222, suggesting that its protein conformation is locked in the dark state configuration (**Fig. 4B**). To characterize the binding efficiency of XL-EL222 to the consensus double-stranded DNA (dsDNA), we used a front-illuminated surface plasmon resonance (fiSPR) biosensor, enabling simultaneous monitoring and *in situ* illumination of interactions at the sensor surface, as described in our previous study.^50^ **Fig. 4C** reports the sensor response to the binding of XL-EL222 and WT-EL222 to dsDNA immobilized on the sensor surface, under *in situ* illumination at a wavelength of 450 nm. We observed a sensor response to XL-EL222 that is more than 25 times lower than that to WT-EL222, indicating a rather poor binding efficiency of XL-EL222 to dsDNA (as also shown in **Supplementary Note S14 (Fig. S15** and **Table S10)**. Taken together, the IR and SPR results suggest that cross-linking the LOV and HTH domains largely blocks the conformational change of XL-EL222 and hinders the exposure of DNA recognition elements (**Fig. 4D**).

**Fig 4.**
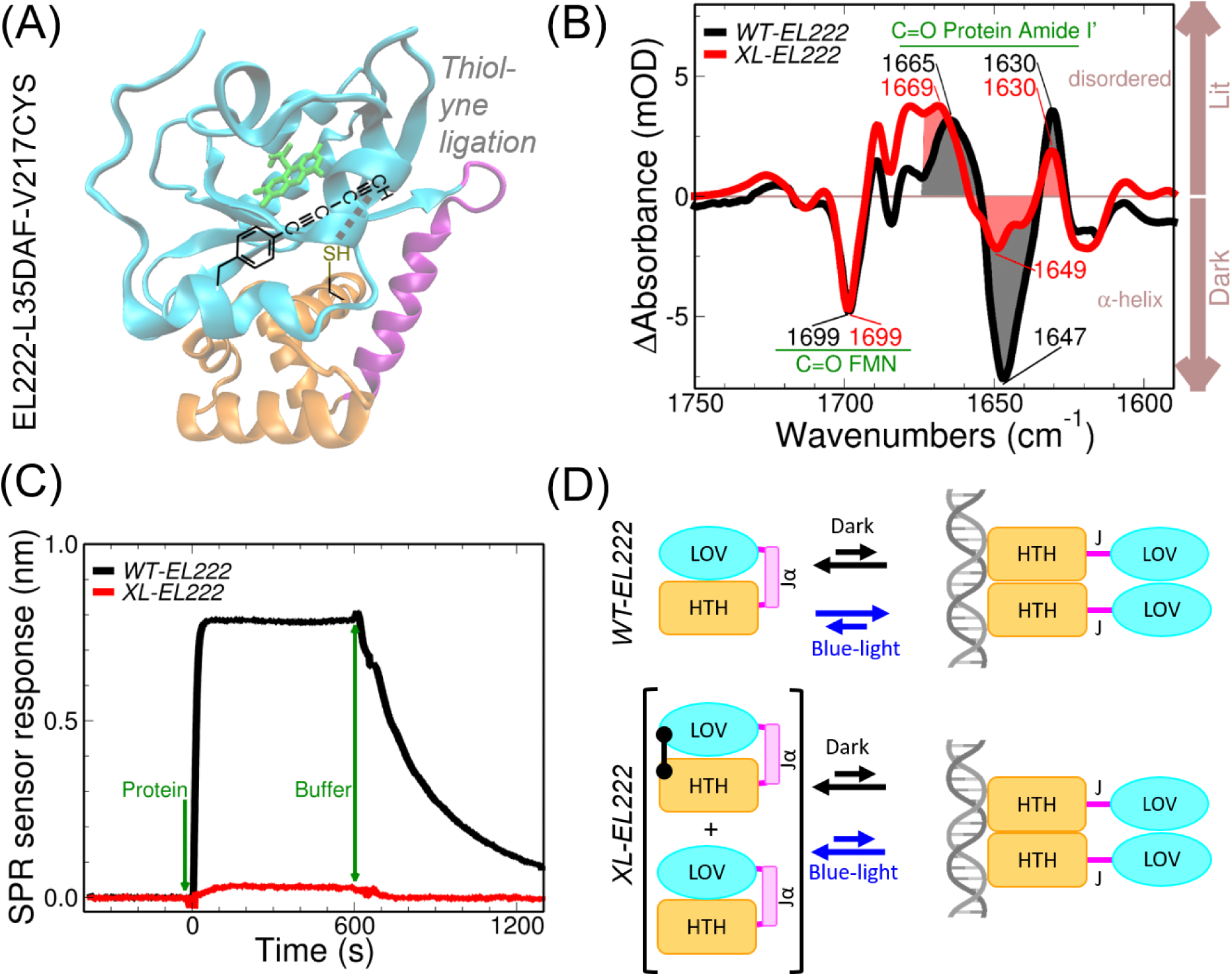
Proximity-enhanced thiol-yne reaction permits intramolecular crosslink formation in EL222. (A) Molecular model of EL222 mutant (based on PDB ID 8A5R), referred to as cross-linked EL2222 (XL-EL222) containing DAF at position 35 (A’α extension of the LOV domain) and a cysteine at position 217 (HTH domain). The dotted lines denote the most probable link between the carbon atoms of the diyne moiety of DAF35 and the sulfur atom of Cys217 sulfhydryl. (B) Light-minus-dark infrared difference spectra of 1 mM both of wild-type EL222 (WT-EL222) and XL-EL222 in the structure-sensitive Amide I’ region (D_2_O-buffered solution). The main bands assigned to C=O stretching modes from either the FMN chromophore or the protein backbone are indicated in reciprocal centimeters units. (C) Surface plasmon resonance (SPR) sensor response to the binding of 6 µM WT-EL222 or XL-EL222 to the attached dsDNA under continuous *in situ* illumination in H_2_O-based buffer solution. (D) Cartoons showing the light-dependent equilibria of WT-EL222 and XL-EL222. The crosslink between LOV and HTH domains is indicated with a black staple. The exact efficiency of cross-linking is unknown and thus a mixture of cross-linked and non-cross-linked species are most likely present in XL-EL222 samples. In (A) and (D) the different domains of EL222 are colored as follows: light-oxygen-voltage domain (LOV, cyan), linker (Jα or J, magenta), and helix-turn-helix (HTH, orange). Jα/J indicate the inferred α-helix/disordered states of the linker between the light-sensitive and DNA-binding domains, respectively.

### FRET of EL222-DAF

Next, we utilized time-resolved Förster resonance energy transfer (FRET), a technique that is sensitive to the distance and orientation between donor-acceptor pairs, to monitor the conformational changes of EL222 in the presence of blue-light (**Supplementary Note S15**). The chemical reactivity of DAF was employed to site-specifically attach a tetrazine-bearing acceptor dye (AZDye594) in residue position 225 (DNA-binding domain) via tetrazine-diyne ligation (**Fig. 5A**). We repurposed the intrinsic FMN chromophore as the fluorescent FRET donor. EL222-I225DAF was first expressed in *E. coli* cells. Subsequently, EL222 containing the ncAA DAF residue was reacted with AZDye594-tetrazine to obtain a donor/acceptor-labeled protein. Next, we measured the fluorescence decays of the fluorescence donor (FMN) in the absence and presence of the fluorescence acceptor (AZDye594) under different illumination conditions. Qualitative differences can already be appreciated (**Fig. 5B**). For further quantification, we extracted lifetime distributions via the maximum entropy method (**Supplementary Note S15**, **Table S11**, **Fig. S16**). Intriguingly, all lifetime distributions feature two main peaks: a longer lifetime peak centered at ∼ 3.5 ns, which is assigned to the donor-only population, and another shorter lifetime peak centered at ∼1 ns, which is assigned to the donor-acceptor population. The calculated lifetime-based FRET efficiencies (*E_FRET_*) and fraction of donor-acceptor lifetime 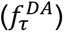 as a function of the time after photoactivation, shown in **Fig. 5C**, suggest a similar FMN-to-AZDye594 distance in all cases (estimated as ∼3.7 nm, **Supplementary Note S15**) but a different ratio of the donor-acceptor population. A plausible molecular interpretation is shown schematically in **Fig. 5D**. The absence of illumination favors a high-FRET “closed” conformation of EL222. Blue-light triggers the formation of a covalent bond between the FMN and the EL222 polypeptide chain (via Cys78). Such a FMN-cysteinyl adduct is essentially non-fluorescent and thus cannot longer act as FRET donor. This causes a drop in the donor-acceptor population following photoactivation. The shorter the time after illumination, the larger the drop. Eventually, the adduct breaks (∼20-60 s depending on the experimental conditions and reporting group)^2,9,51^ and EL222 returns to the dark-state conformation. Thus, the fluorescent decays collected at long post-illumination times resemble those in the absence of illumination. Overall, our DAF-enabled FRET-based assay reports on conformational equilibria along EL222 photocycle.

**Fig. 5.**
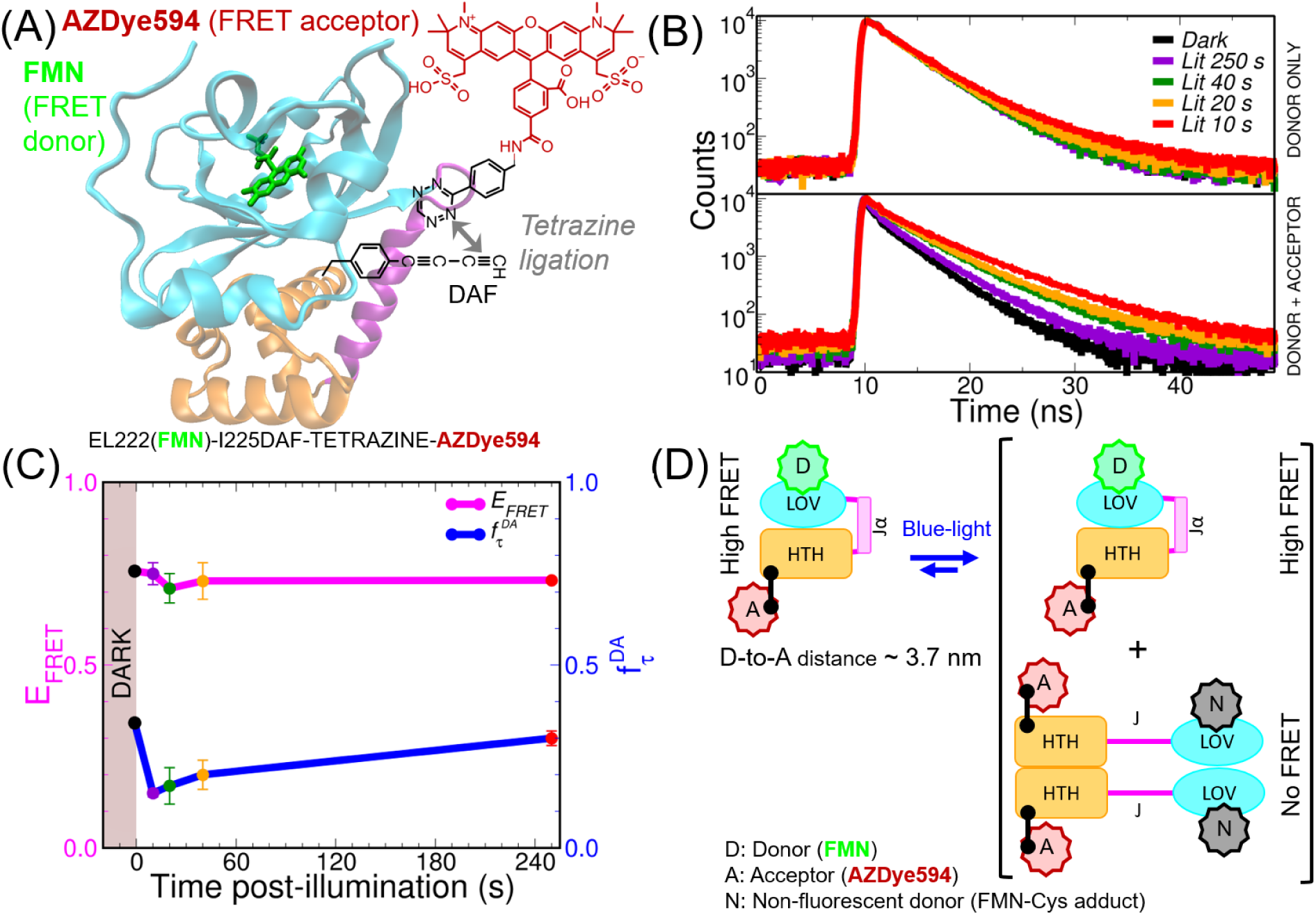
Tetrazine ligation with DAF enables the site-specific installation of a donor-acceptor pair for distance measurements by FRET. (A) Molecular model of EL222 encoding DAF at position 225, which is subsequently reacted with an H-tetrazine-containing acceptor dye (AZDye594) via tetrazine-diyne ligation. The intrinsic FMN cofactor acts as the FRET donor. (B). Representative fluorescence decay curves of donor-only EL222 natively containing FMN and donor/acceptor EL222-I225DAF-Tetrazine-AZDye594 samples (λ_excitation_= 450 nm, λ_emission_= 525 nm), both at a nominal concentration of 15.6 μM. Five different experimental conditions were studied: dark (no pre-illumination) and “lit” (four different times post-illumination with an LED emitting 35 mW at 455 nm). (C) Lifetime-based FRET efficiencies (*E_FRET_*) and fraction of donor-acceptor lifetime 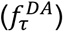 as a function of the elapsed time after photoactivation of the sample with the aforementioned LED. The points and bars represent the mean and standard deviations of two independent experiments. The samples measured in the dark are arbitrarily set at time -1 s. (D) Cartoon representing a molecular interpretation of the observed effects. Blue-light shifts the conformational equilibrium of EL222 towards the “active” (DNA-binding-competent dimer conformation), which is invisible to FRET (due the presence of a poorly fluorescent FMN-cysteinyl adduct state) and slowly returns to the dark-state conformation on a time scale of seconds. The distance between FMN and AZDye594 keeps constant (∼ 3.7 nm) across the five experimental conditions.

### Raman spectroscopy of EL222-DAF

To exploit the *a priori* convenient Raman properties of DAF as a site-specific reporter of protein microenvironment, we measured femtosecond-stimulated Raman spectra (FSRS) of EL222-DAF, first at the steady-state level and then in a time-resolved manner.

Initially, we screened for residue positions that show light-dependent Raman spectral changes. We selected three positions: L35, M151, and I225, which are located in the N-terminal extension of the LOV domain (so called A’α helix), in the linker between the LOV and HTH domains, and in the DNA-binding domain, respectively. As a reference, the dark *vs.* lit infrared spectra of the three CNF mutants (L35CNF, M151CNF, I225CNF) in the C≡N stretching band upon blue-light illumination are shown in **Fig. S17 (Supplementary Note S16**). The corresponding DAF mutants (L35DAF, M151DAF, and I225DAF) are displayed in **Fig. 6B** and **Fig. S18 (Supplementary Note S17)**. Two conclusions can be extracted. The distinct responses suggest that CNF and DAF sense different aspects of the local chemical environment. Additional FSRS experiments can be found in **Supplementary Note S17**, **Fig. S18**).

**Fig. 6.**
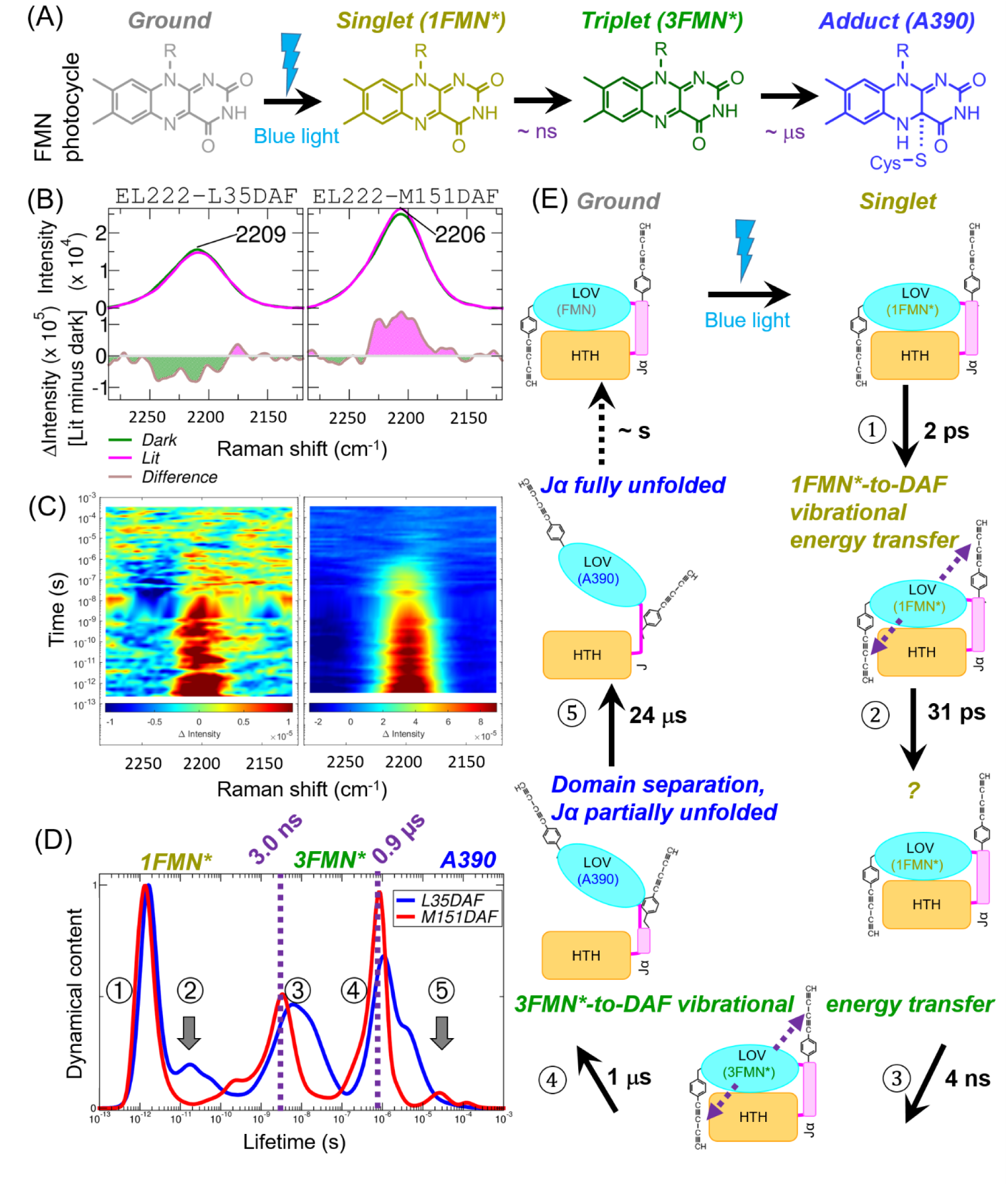
Time-resolved stimulated Raman spectroscopy of DAF-containing EL222 variants. (A) Consensus photocycle of light-oxygen-voltage (LOV) photoreceptors as revealed by multiple optical spectroscopy methods. (B) Steady-state femtosecond-stimulated Raman spectra (FSRS, λ_Raman pump_ = 800 nm) of two EL222^DAF^ mutants in the dark and under continuous blue light illumination (λ = 455 nm, 15 mW). Absolute spectra are shown in the top panels and difference (light-minus-dark) spectra are shown in the bottom panels. (C) 2D contour plots of differential Raman signal as a function of pump-probe time delays and wavenumbers of EL222-L35DAF (*left*) and EL222-M151DAF (*right*) mutants. (D) Normalized average dynamical content in the 2100-2300 cm^-1^ spectral region arising from lifetime distribution analysis of EL222-L35DAF and EL222-M151DAF. The three consensus FMN intermediates and the two corresponding transitions are indicated. The circled numbers indicate the five dynamical events revealed by time-resolved FSRS. (E) Extended photocycle inferred from site-specific time-resolved FSRS of EL222^DAF^. The cartoons show a plausible molecular interpretation of the five observed dynamical events. Events 3 and 4 correspond to the well-known singlet-to triplet and triplet-to-adduct transitions, respectively. The dotted violet arrows indicate the direction of energy transfer from excited FMN (singlet or triplet) to DAF. DAF35 is located in the A’α helix of the LOV domain and senses the close-open conformational equilibrium of EL222. DAF151 is the located in the linker region between the LOV and HTH domains and senses the folded-unfolded state of the Jα element. Event 2 has not been assigned.

The photocycle of EL222, like other LOV-containing proteins, is characterized by singlet, triplet, and adduct intermediates, which appear sequentially after blue-light excitation (**Fig. 6A**). To determine whether DAF can unveil more site-specific dynamical events, we monitored the photoactivation dynamics of EL222 with time-resolved FSRS from femtoseconds to milliseconds focusing on the “transparent window” region between 2100-2300 cm^-1^ (**Fig. 6C**).

Important dynamical information can be extracted by looking at the lifetimes. A convenient way to visualize the non-equilibrium dynamics is to plot the average dynamical content as a function of the lifetime (**Fig. 6D**). Lifetime distribution analysis of EL222-L35DAF and EL222-M151DAF (**Fig. 6D**) suggested the presence of four dynamical events in each sample with distinct spectral features (**Supplementary Note S18, Fig. S19**). As a reference, we also plotted the dynamical content arising from transient absorption (TA) in the 300-1000 nm region (**Supplementary Note S19, Fig. S20**), which reports exclusively on the FMN chromophore and immediate surroundings. Analysis of the TA datasets arising from EL222-L35DAF and EL222-M151DAF reveal the two well-known transitions: a 3 ns singlet-to-triplet transition, and a 0.9 μs triplet-to-adduct transition, with virtually identical timescales in both mutants. Interestingly, the same two events arise upon analysis of time-resolved Raman spectra in the 150-2000 cm^-1^ region^5^ and time-resolved infrared spectra in the 1500-1800 cm^-1^ region.^6^ However, FSRS in the cell-silent diyne region shows evidence of additional site-specific dynamics, which can be tentatively interpreted in molecular terms (**Fig. 6E**). Shortly after FMN excitation by the pulsed laser, we have a picosecond event in both positions (DAF35 and DAF151). This raises the possibility that the first component originates on vibrational energy transfer from excited FMN (singlet) acting as the donor and the diyne of DAF acting as the acceptor. The second dynamical event of L35DAF with a lifetime of 31 ps is more difficult to interpret. Sub-100 ps events have been previously observed, principally in free FMN, and assigned to ribityl-phosphate folding dynamics, solvent relaxation and H-bond rewiring.^5,52^ In principle, DAF35 could be sensing some of these molecular processes. Both mutants report on an event occurring at 4 ns, which could be related to vibrational energy transfer from the rising triplet state of FMN. The two mutants differ mainly at the latest delays. L35DAF shows dynamics largely overlapping with FMN-Cys78 adduct formation (1.1 μs). M151DAF has two dynamics, one concomitant (∼0.6 μs) and the other slower (∼24 μs) than triplet decay. Given the location of DAF151 in the α-helical linker region (Jα), which is expected to unfold thereby enabling the separation of LOV and HTH domains, such two events may be viewed as a two-step unfolding. Indeed, the complete unfolding of the bridging Jα helix was previously inferred to proceed multi-step-wise in other LOV domains.^53,54^

Importantly, the strong Raman signal of DAF can also be detected by spontaneous Raman microscopy in live *E. coli* cells (**Supplementary Note S20, Fig. S21**). However, these experiments require high intracellular concentrations (> 0.5 mM), which can be achieved in the case of MBP but not in the case of EL222.

We can conclude safely that the alkyne moiety of DAF reports on local dynamical events, reflecting its location in the protein, which can be tracked by time-resolved FSRS. These events are partially unique and partially redundant with other commonly used probes, like the FMN chromophore.

A summary of all the useful properties of DAF can be found in **Supplementary Note S12** (**Table S9**).

## Discussion

The practical utility of novel ncAA heavily depends on the existence of reliable platforms for their genetic encoding, an aspect sometimes overlooked in the field. Our DAF-specific amino acyl-tRNA synthetase from *Methanomethylophilus alvus* (*Ma*DafRS) has significant advantages over other encoding systems in terms of its broad orthogonality (both prokaryotes and eukaryotes may be utilized as expression hosts), superior fidelity and enhanced suppression yields. The latter two properties result in near-native levels of biosynthesized proteins with minimal heterogeneity at the recoded site. Thus, it should be possible to pair *Ma*DafRS with another aaRS and ncAA to achieve dual encoding and labeling (DEAL).^55^

DAF represents a new tool that enables the chemical modification of proteins and can also act as a Raman-active tag. The main advantage of DAF as a chemical handle is its multi-reactivity properties against azides, H-tetrazines, and thiols. From the point of view of the reactivity against tetrazines, the diyne of DAF behaves more similarly to a strained alkyne than to a linear alkyne. However, from the point of view of the reactivity against azides, DAF behaves closer to a linear mono-alkyne given the strict requirement for Cu(I) as a catalyst. That is to say, DAF primarily follows a CUAAC-type reaction mechanism, while a SPAAC-type reactivity is virtually absent. For instance, SCK shows fast kinetics against H-tetrazines and slow kinetics against azides,^56–58^ while PRK reacts rapidly towards azides and poorly towards tetrazines.^59,60^ Out of the four criteria that define an “ideal” bioorthogonal reaction: 1) speed, 2) selectivity, 3) stability, and 4) size,^61^ DAF does not truly fulfill the first one. Therefore, DAF cannot compete against other faster reacting ncAA, like those containing tetrazine,^62^ *trans*-cyclooctene,^63^ and bicyclononyne^64^ moieties. Nevertheless, the intermediate reaction kinetics of DAF may be compensated in different ways, e.g. by a proximity-enabled strategy or second-generation reactive dyes (AzidePlus), both illustrated in the present work. Alternatively, a careful optimization of the reaction conditions, as detailed in other studies,^65,66^ may be attempted in the future.

The “holy grail” of photosensory biology is the nature of the ultimate driving force of protein conformational changes. FMN-cysteinyl adduct formation has long been considered the “point of no-return” starting all subsequent protein rearrangements in LOV photoreceptors. This paradigm has been revisited to account for the fact that cysteine-less LOV mutants can still be activated by light and perform physiological roles, like regulation of the circadian clock or gene expression.^67,68^. These observations have been rationalized as photoinduced flavin reduction to the neutral semiquinone form being sufficient to trigger changes in LOV protein structure and function. The results arising from our proximity-enabled thiol-yne (cysteine-DAF) click chemistry reaction suggest that the helix-to-coil transition (unfolding of Jα and/or A’α) is the key event driving the functional photoactivation of EL222. By covalently linking the two individual domains of EL222, we effectively decoupled FMN photochemistry from protein structural/functional changes: FMN undergoes a normal photocycle (singlet → triplet → adduct) but these changes in the chromophore do not cause EL222 self-association nor interactions with DNA. In other words, allosteric communication in cross-linked EL222 is largely halted. The protein keeps the secondary structure and monomeric status of the dark-state conformation, which is essentially unable to bind DNA, as demonstrated by fiSPR biosensing experiments, and thus inactive from a functional point of view. This adds to a series of investigations into Jɑ unfolding.^69,70^

The light-dependent structural dynamics of EL22 was directly addressed by measuring LOV-HTH distances. In turn, the time-resolved FRET experiments were made possible by site-specifically labeling EL222 with a fluorescent acceptor dye via a tetrazine-diyne reaction. Dark-state and lit-state EL222 give rise to high-FRET and no-FRET species, respectively. The fraction of donor-acceptor reports on the dark recovery rate. These results underscore the dynamic behavior of EL222, which changes significantly after illumination and this shifts the conformational equilibria (domain separation, secondary structure), in line with previous findings.^8^ However, no distance information can be obtained from EL222 lit state using the intrinsic FMN moiety as the FRET donor. Direct FRET characterization of the signaling state of EL222 will necessitate a three-chromophore system in which FMN is utilized as the phototrigger, and a spectrally separated donor-acceptor dye pair is utilized as a spectroscopic ruler.

Site-specific time-resolved FSRS assisted by DAF provides a wealth of information about EL222 relaxation dynamics following FMN photoexcitation. Surprisingly, the diyne of DAF seems to act as an “antenna” moiety sensing the two excited states of FMN (singlet and triplet) despite the absence of a direct physical linkage (neither covalent bonding nor hydrogen bonding) and the relatively long distance between the two (more than 0.5 nm). This phenomenon, which awaits further scrutiny, raises the possibility of utilizing DAF as an “acceptor” in vibrational energy transfer studies of proteins.^71,72^ The critical role of the linker between the light-sensing and DNA-binding domains is also indirectly assessed by FSRS in the alkyne region. It is generally assumed that N5 protonation, adduct formation and loss of α-helicity happen simultaneously.^4,6^ However, such a view is challenged by our observation of linker dynamics occurring after adduct formation. We hypothesize that, apart from FMN-Cys covalent bond formation, dynamic fluctuations in the linker region play a crucial role in EL222 signal transduction and potentially in other LOV photosensors. Supporting our findings of site-specific protein dynamics being asynchronous with FMN dynamics, fluctuations in the N-terminal A’α helix precede adduct breakage in the dark recovery of EL222.^9^

Overall, our new Raman-active ncAA DAF enables site-specific Raman spectroscopy of proteins. FSRS of EL222-DAF mutants reveal that the LOV photocycle is more complex than previously found by transient stimulated-Raman in the “fingerprint” region from 200 to 2000 cm^-1,5^ transient infrared absorption in the 1500-1800 cm^-1^ region,^6^ and transient absorption in the 300-1000 nm.^5^ All three optical spectroscopies converge in the consensus LOV photocycle showing two major transitions: a nanosecond singlet-to-triplet transition, and a microsecond triplet-to-adduct transition.^4,7^ Inclusion of DAF reporters provides much richer information adding up to 3 dynamical events (vibrational energy transfer from singlet FMN, vibrational energy transfer from triplet FMN, and full Jα unfolding) in the photoactivation pathway of EL222. Nevertheless, future joined experimental-computational efforts are needed to fully exploit the Raman information content of DAF.

All these fabulous properties of DAF merit its inclusion in the modern molecular engineering toolbox, especially regarding protein conjugation and dynamics.

## Acknowledgments.

This study was supported by the Czech Science Foundation, contracts 24-11819S (G.F.*) and 25-15726S (J.S.*). The Institute of Biotechnology of the Czech Academy of Sciences acknowledges the institutional grant RVO 86652036. We acknowledge CF Biophysics, CF SMS of CIISB, Instruct-CZ Centre BIOCEV, supported by MEYS CR (LM2023042) and ERDF-Project “Innovation of Czech Infrastructure for Integrative Structural Biology” (No. CZ.02.01.01/00/23_015/0008175). We acknowledge funding by the Charles University Grant Agency (GAUK 407622). We also acknowledge Czech-BioImaging project (MEYS CR, project number LM2023050 Czech-BioImaging). The authors acknowledge the Imaging Methods Core Facility at BIOCEV for their support with obtaining flow cytometry and microscope image data presented in this paper. We acknowledge travel funding from GCEforAll center (Oregon State University, USA) for attending the course “Selecting your synthetase”

## Author contribution

G.F.^1^,*, A.C., P.N.P. designed the experiments; J.S.^6^,*, M.S., J.S.^1^, L.W. synthesized and characterized DAF; P.N.P. performed evolution of *Ma*DafRS; A.C. performed the small molecule chemical reactivity experiments; A.C., P.N.P.,O.H. performed in solution and *in-cell* protein labelling experiments; P.N.P. performed fluorescence microscopy experiments; A.C., A.M., P.C., J.D., S.K., M.K.^2^ performed FSRS and TA experiments; A.C., A.S.C., G.F.^3^, T.S., J.H. performed experiments with the cross-linked mutant; A.C. performed FRET experiments; A.C., M.K.^5^ performed in-cell Raman microspectroscopy; G.F.^1,^*, J.S.^6,^*, A.C., P.N.P. analyzed the data and wrote the paper with contributions from all authors.

## Abbreviations

ACN: Acetonitrile
AZF: 4-Azido-phenylalanine
AZK: N⁶-(2-azidoethoxy)carbonyl-L-lysine
BCK: Bicyclononyne-lysine
BME: β-Mercaptoethanol
BSA: Bovine serum albumin
CaF₂: rCalcium fluoride
CDCl₃: rDeuterated chloroform
CNF: r4-Cyano-phenylalanine
CuI: rCopper(I) iodide
DAF: r4-Diacetylenyl-phenylalanine
DCM: rDichloromethane
DEA: rDiethylamine
DMSO: rDimethyl sulfoxide
DMSO-*d6*: rDeuterated dimethyl sulfoxide
DPY: rDiphenylbutadiyne
DPA: rDiphenylacetylene
DTT: rDithiothreitol
EA: rEthanolamine hydrochloride
EDC: r1-ethyl-3-(3-dimethylaminopropyl)carbodiimide hydrochloride
EDTA: rEthylenediaminetetraacetic acid
EDU: r5-Ethynyl-2′-deoxyuridine
ETF: r4-Ethynyl-phenylalanine
EtOAc: rEthyl acetate
FMN: rFlavin mononucleotide
GFP: rGreen fluorescent protein
HCl: rHydrochloric acid
HPG: rHomopropargyl-glycine
IPTG: rIsopropyl β-D-1-thiogalactopyranoside
KCl: rPotassium chloride
KHSO₄: rPotassium hydrogen sulfate
LB: rLuria broth
MBP: rMaltose-binding protein
MES: r2-(N-morpholino)ethanesulfonic acid
MeLi: rMethyllithium
MgSO₄: rMagnesium sulfate
NaCl: rSodium chloride
NaOH: rSodium hydroxide
Na₂SO₄: rSodium sulfate
NH₄Cl: rAmmonium chloride
NHS: rN-hydroxysuccinimide
PAE: rPhenylacetylene
PdCl₂(PPh₃)₂: rBis(triphenylphosphine)palladium(II) dichloride
PMSF: rPhenylmethylsulfonyl fluoride
PPh₃: rTriphenylphosphine
PRY: rPropargyl-tyrosine
PRK: rPropargyl-lysine
SCO-PEG3-NH₂: rStrained cyclooctyne-PEG₃-amine
SCOK: rStrained cyclooctyne-lysine
SDS: rSodium dodecyl sulfate
TB: rTerrific broth
TBAF: rTetrabutylammonium fluoride trihydrate
TCSPC: rTime-correlated single photon counting
TCEP: rTris(2-carboxyethyl)phosphine
TEA: rTriethylamine *(verify vs. triethanolamine if needed)*
TFA: rTrifluoroacetic acid
Tet.2-Et: r(*2S*)-2-amino-3-[4-(6-ethyl-1,2,4,5-tetrazin-3-yl)phenyl]propanoic acid
THF: rTetrahydrofuran
THPTA: rTris(3-hydroxypropyltriazolylmethyl)amine
TCO*K: rtrans-Cyclooct-2-en-1-yloxycarbonyl-lysine
TMS-C4-TMS: r1,4-Bis(trimethylsilyl)butane

## Methods

### Materials

**Table 0.**
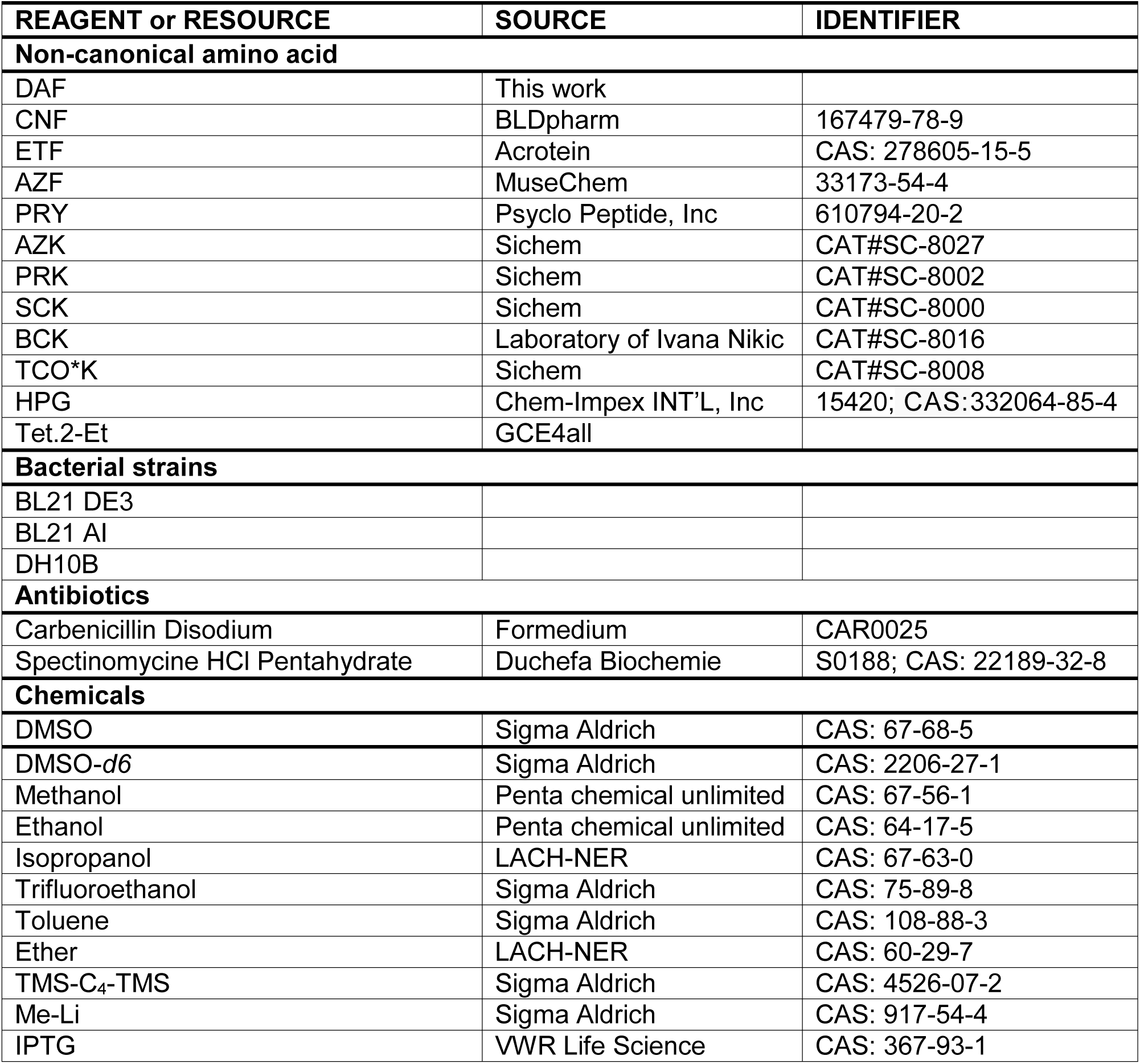
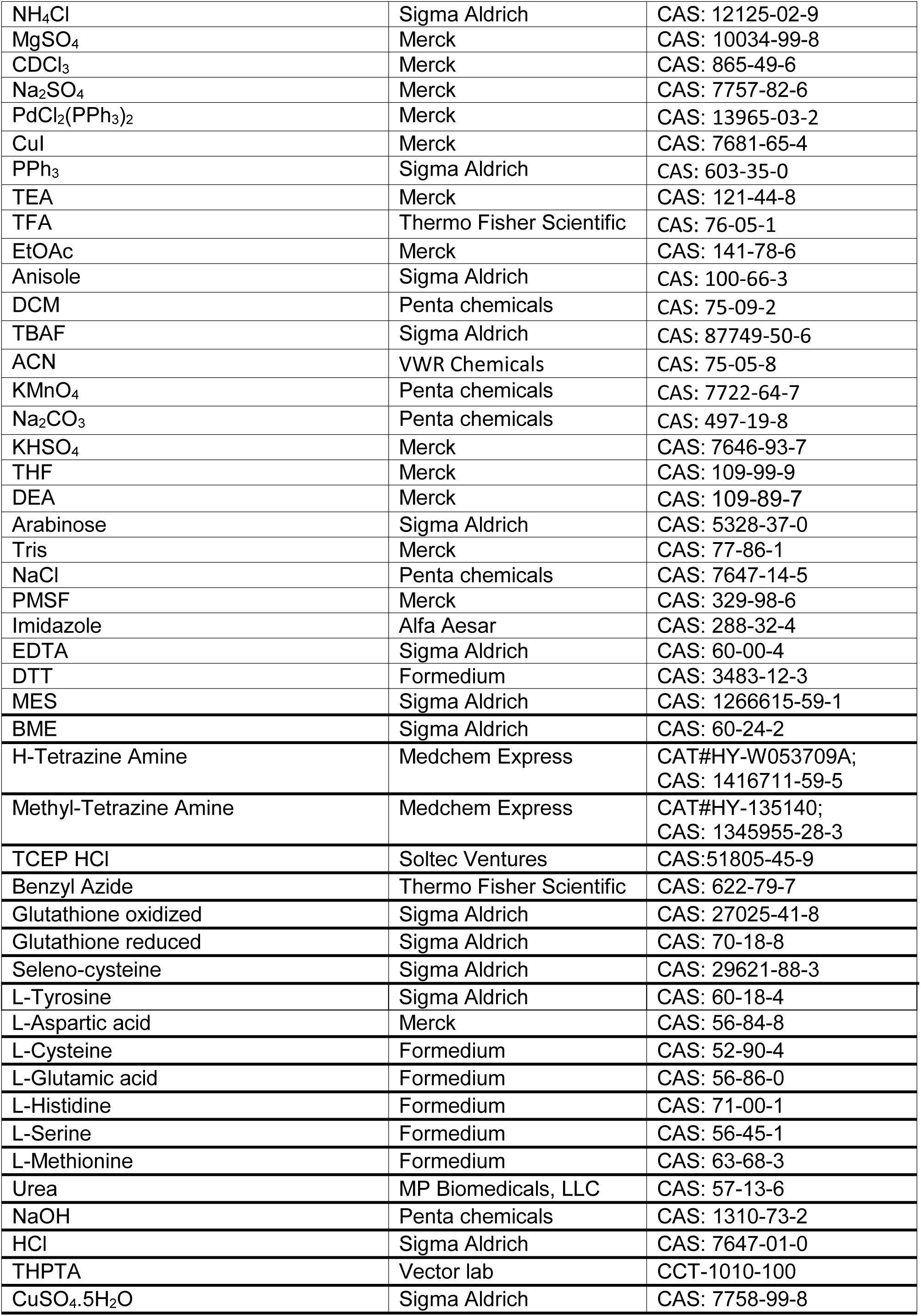
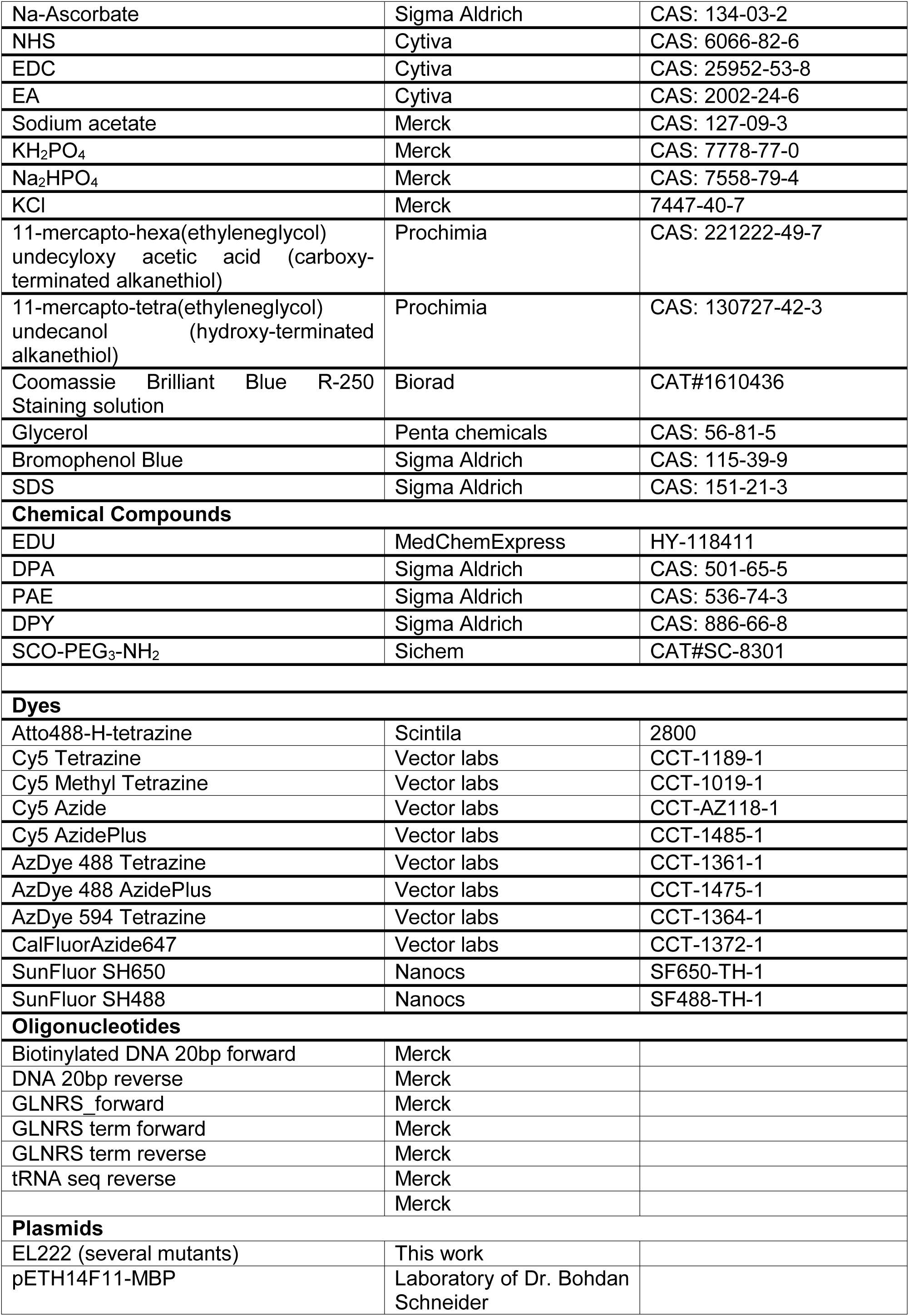
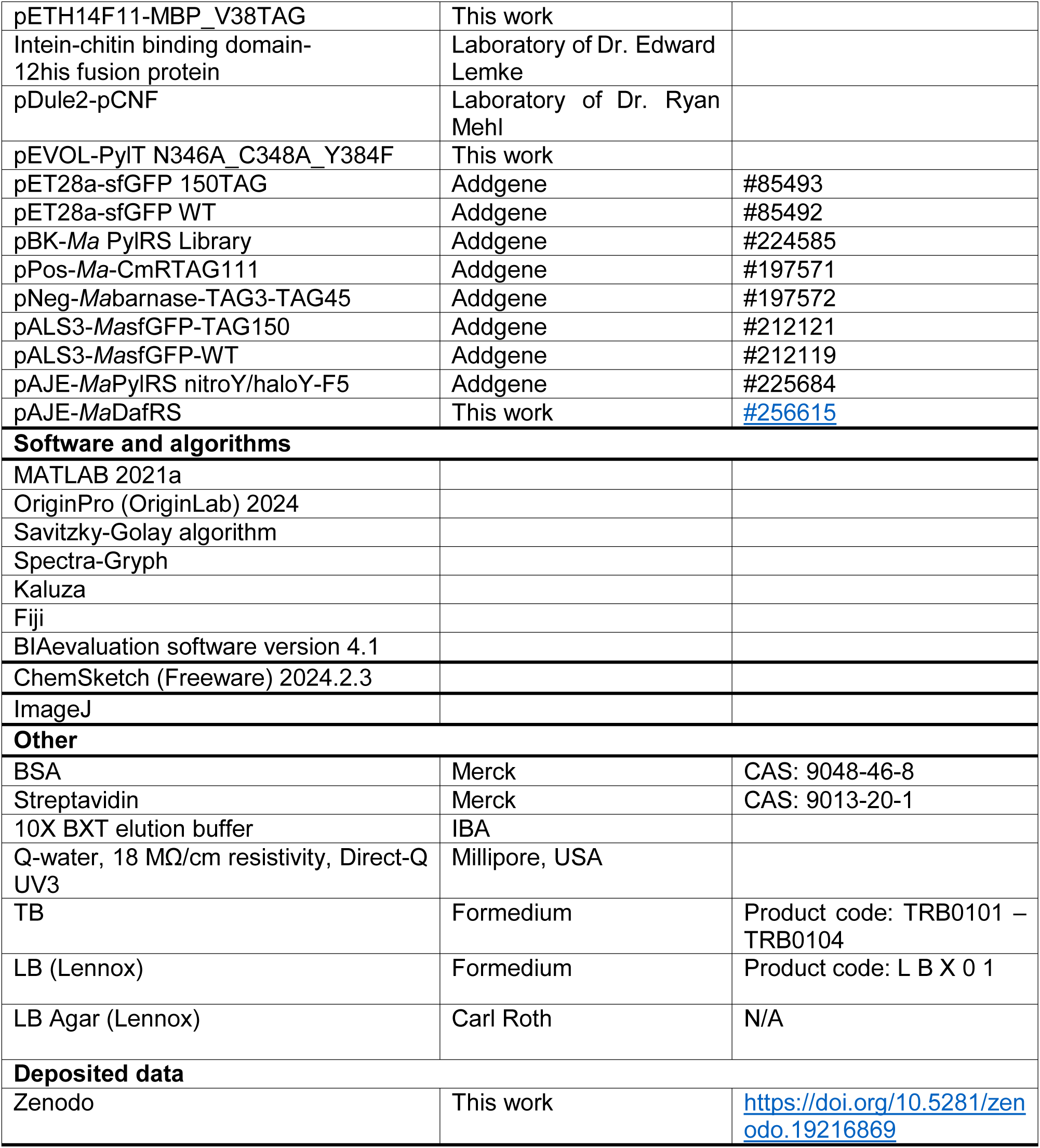
Key resources table.

## 1. DAF synthesis

All reagents were purchased from local chemical suppliers (for details see Table above) and used without further purification, except for drying of ether, and tetrahydrofuran (THF). Detailed synthesis procedures are in **Supplementary Note S1.** The products were dried in a vacuum drying box at room temperature for 16 h. Progress of syntheses was monitored by TLC on silica gel-coated aluminum plates. The compounds were visualized by exposure to UV light at 254 nm or by KMnO_4_/Na_2_CO_3_ spraying.^73^ Alternatively, the HPLC was also used for monitoring the reactions. Analytical HPLC was carried out with a Poroshell 120 SB-C18 2.7 μm, 3.0 × 50 mm column, a flow rate 1 mL/min, and diode array and evaporative light scattering detectors using gradient 5–5–100–100% of ACN in 0.05% aqueous TFA within 0-1-11-13 min at 40 °C. The ^1^H-NMR and ^13^C-NMR, spectra were measured using Bruker Avance II™ spectrometers with these frequency setups (^1^H at 400 MHz and ^13^C at 100 MHz frequency) in DMSO-*d6* or CDCl_3_. Proton and carbon-13 signals were structurally assigned by means of the series of 1D-(^1^H, ^13^C) and 2D-NMR spectra (^1^H,^1^H-COSY, ^1^H,^13^C-HMQC, ^1^H,^13^C-HMBC). During the syntheses, molecular weights of compounds were determined using electrospray ionization mass spectroscopy (ESI).

## 2. Protein preparation and characterization

### 2.1. Expression and purification of GFP N150DAF Mutant

BL21AI cells were electroporated with pET28a-sfGFP WT for expression of wild-type sfGFP. For incorporation of DAF, BL21AI cells were co-electroporated with pET28a-sfGFP N150TAG together with either pAJE-*Ma*DafRS, pDule2-CNF, or pEVOL-PylT-N346A/C348A/Y384F. Single colonies were selected on LB agar plates supplemented with the appropriate antibiotics: chloramphenicol for pEVOL-PylT-N346A/C348A/Y384F, spectinomycin for pAJE-*Ma*DafRS and pDule2-CNF, and kanamycin for pET28a-sfGFP WT and pET28a-sfGFP N150TAG. Colonies were inoculated into LB medium containing the corresponding antibiotics and cultured overnight at 37 °C with shaking at 180 rpm. Overnight cultures were diluted 1:100 into TB medium supplemented with antibiotics and incubated at 30 °C with shaking at 180 rpm until the optical density at 600 nm (OD₆₀₀) reached 0.4–0.5. DAF, prepared as a 0.25 M stock solution in methanol, was added to the appropriate cultures to a final concentration of 0.25 μM. After 30 min, protein expression was induced by the addition of IPTG and arabinose to all cultures, including DAF-positive, DAF-negative, and WT controls. Cultures were further incubated for 20 h at 25 °C with shaking at 180 rpm.

Cells were harvested the following day, and OD₆₀₀ values were measured after 20-fold dilution in PBS. The lowest OD₆₀₀ value obtained among all samples (Mj without DAF; OD₆₀₀ = 5.64) was used for normalization. All samples were adjusted to OD₆₀₀ = 5.64 in 1 mL PBS (pH 7.4). Cell suspensions were transferred to black 96-well plates, and fluorescence intensity was measured using a TECAN plate reader.

For protein purification, 15 mL overnight cultures of GFP WT and 30 mL cultures of all other samples were harvested by centrifugation. Cell pellets were resuspended in buffer containing sodium phosphate, 300 mM NaCl, and 5 mM imidazole, pH 7.5 and lysed by sonication. Lysates were clarified by centrifugation at 60,000g for 30 min at 4 °C. Supernatants were applied to 100 μL Ni-NTA agarose resin pre-equilibrated with same buffer. The resin was washed with 2 mL buffer, and bound proteins were eluted in six 100 μL fractions containing sodium phosphate, 300 mM NaCl, and 500 mM imidazole, pH 7.5. Purified proteins were analyzed by SDS–PAGE and imaged using an Azus 600 instrument with a Cy2 filter for GFP detection, followed by Coomassie Brilliant Blue staining. The pooled protein fractions were subsequently analyzed by mass spectrometry.

### 2.2. Expression and purification of EL222 Wild Type (17-225)

An ampicillin-resistant plasmid encoding EL222 WT (pEL222-TEV-6His_BamHI) was transformed into *Escherichia coli* BL21(DE3) by electroporation (2.5 kV for 5 ms in a 2 mm path-length cuvette) as described in ^9^. Transformed cells were grown in LB-Agar plate containing carbenicillin (50 µg/ml) antibiotic at 37°C overnight. A single colony was picked to make a starter culture in LB media and grown overnight at 37°C at 200 rpm. The starter culture was diluted 100 times in TB media supplemented with carbenicillin (50µg/ml). The bacteria were grown at 37°C at 180 rpm until an optical density of 0.5 at 600 nm at which point the temperature was subsequently reduced to 20°C. Then gene expression was induced by adding IPTG to a final concentration of 0.5 mM and 0.2% Arabinose. Cells were further grown for 22 hours in the dark.

The cells were harvested by centrifugation at 6,000 *×g* for 15 min. The cells in the pellet were resuspended in 50 mM Tris, 100 mM NaCl, pH 8 buffer, with 1 mM PMSF, and disrupted by sonication for 4 min on ice (50 mW). After repeating the centrifugation of the cell suspension at 60,000 *×g* for 30 min, the supernatant was purified by immobilized metal affinity chromatography with a Ni^2+-^NTA resin (purchased from ThermoFisher Scientific, Germany). 20 mM imidazole was used to wash away the loosely bound proteins, and 200 mM imidazole was used to elute the fusion proteins. The plasmid used for protein expression presented a C-terminal TE purification tag. The eluted protein was diluted in half with water to reduce Imidazole concentration which may denature TEV protease. EL222 concentration was estimated based on its FMN by checking absorbance at 450 nm assuming ε_450_= 13000 M^-1^cm^-1^ and 100% FMN loading. TEV was added to protein in 1:10 mass ratio and dialyzed against 50 mM Tris, pH 8; 0.5 mM EDTA and 1 mM DTT for 24 hours to remove this purification tag. A second dialysis was performed against 50 mM Tris, 100 mM NaCl, pH 8 to remove EDTA and DTT that may interfere with downstream process. Next a second affinity chromatography step was performed to remove the uncleaved proteins from the solution containing the cleaved proteins, which were further polished using SEC in a Superdex75 Increase 10/300 GL column (purchased from Cytiva) with 50 mM MES, 100 mM NaCl, pH 6.8 as running buffer. Fractions were analyzed by SDS-PAGE to identify EL222 (MW ∼ 25 kDa), pooled and concentrated using 10 kDa molecular-weight cut-off filters (Sartorius). The purified EL222 was flash-frozen in liquid nitrogen and stored at – 80 °C before its use.

### 2.3. Expression and purification of EL222 DAF and ETF Mutants

In EL222 WT plasmid (pEL222-TEV-6His_BamHI) each single residue leucine 35, methionine 151 and isoleucine 225 sense codon were recoded with amber codon TAG by site-directed mutagenesis to obtain plasmid EL222-L35TAG, EL222-M151TAG and EL222-I225TAG. DNA sequences were checked. To get an intramolecular crosslinked EL222 DAF mutant (EL222-L35DAF-V217C), the valine 217 residue was recoded with cysteine in the plasmid encoding EL222 L35TAG. Each obtained ampicillin-resistant EL222 plasmid bearing one TAG triplet at desired positions was co-transformed by electroporation into *Escherichia. coli* BL21 (DE3) with spectinomcyin-resistant plasmid pDule2-CNF encoding an OTS able to incorporate DAF/ETF in response to TAG codon except EL222-I225TAG, which was co-transformed with spectinomycin-resistant plasmid pAJE-*Ma*PylRS_DAF as described in section 2.2. Individual transformed cells were grown in LB-Agar plate containing carbenicillin (50 µg/ml) and spectinomycin (100 µg/ml) antibiotics at 37°C overnight. A single colony was picked to make a starter culture in LB media and grown overnight at 37°C at 180 rpm. The starter culture was diluted 100 times in TB media supplemented with carbenicillin (50µg/ml) and spectinomycin (100 µg/ml). The bacteria were grown at 37°C at 180 rpm until an optical density of 0.5 at 600 nm at which point the temperature was subsequently reduced to 20°C. In the culture, freshly prepared stock of either 0.5 M pDAF in 100% DMSO or 1 mM ETF dissolved in 3M NaOH was added to obtain the respective site-specifically labeled mutants EL222-L35DAF, EL222-M151DAF, EL222-I225DAF, EL222-L35DAF-V217C and EL222-M151ETF. Cells were incubated for additional 30 mins and then gene expression was induced by adding IPTG to a final concentration of 0.5 mM. Cells were further grown for 22 hours in the dark.

From cell harvesting to protein purification, the procedures were performed as described in section 2.2 with the only exception of using 10mM imidazole as washing buffer for EL222 mutants during first metal affinity chromatography.

### 2.4. Expression and purification of MBP-WT, MBP-V38DAF and MBP-V38ETF mutants

BL21(DE3) cells were electroporated with the plasmid pETH14F11-MBP to express wild-type MBP protein. For incorporation of noncanonical amino acids, BL21(DE3) cells were electroporated with pETH14F11-MBP_V38TAG together with either pAJE-*Ma*DafRS or pDule2-CNF to produce MBP V38DAF and MBP V38ETF, respectively. Single colonies selected on appropriate antibiotic containing agar plates (spectinomycin for plasmid pAJE-*Ma*DafRS and pDule2-CNF, while ampicillin for plasmid pETH14F11-MBP and pETH14F11-MBP_V38TAG) were inoculated into LB medium supplemented with the antibiotics and grown overnight at 37 °C with shaking at 180 rpm. The following day, overnight cultures were diluted 1% into 200 mL TB medium containing antibiotics and incubated at 30 °C with shaking at 180 rpm until the optical density (OD₆₀₀) reached 0.5–0.6. DAF, prepared as a 0.5 M stock solution in methanol, and ETF, prepared as a 1 M stock solution in 3 M NaOH, were added to the respective cultures to final concentrations of 0.5 mM and 1 mM. After 30 minutes, IPTG was added to all three cultures to induce protein expression for 22 hours at 25 °C with shaking at 180 rpm.

170 ml cells were harvested by centrifugation at 6,000 g for 15 min at 4 °C and resuspended in lysis buffer 50 mM Tris-HCl, 100 mM NaCl, 1 mM PMSF, pH8. Cells were lysed by sonication and lysates were clarified by centrifugation at 60,000 g for 30 min at 4 °C. The supernatant was applied to 300 µL Strep-Tactin^®^XT resin (IBA) pre-equilibrated in buffer 50 mM Tris-HCl, 100 mM NaCl, 1 mM EDTA, pH8. The resin was washed with 10 mL of the same buffer to remove unbound protein, and bound MBP proteins were eluted with 3 mL (10 column volumes) of Strep-Tactin^®^XT elution buffer (IBA). Eluted proteins were concentrated to 1 mL using a 10 kDa molecular weight cutoff concentrator at 6,000 g for 5 min at 4 °C and further clarified by centrifugation at 20,000 g for 20 min at 4 °C. Samples were subjected to size-exclusion chromatography on a Superdex 75 Increase 10/300 GL column equilibrated in buffer 50 mM Tris-HCl, 100 mM NaCl, pH 8. Peak fractions were collected and concentrated for subsequent labeling with dye.

### 2.5. Characterization of proteins

Concentration of all the proteins were quantified using UV-visible absorption spectroscopy assuming ε450 = 13,000 M^−1^cm^−1^ and 100% FMN loading for EL222 proteins and assuming ε488 = 55000 M^−1^cm^−1^ for GFP proteins, and assuming ε280 = 73340 M^−1^cm^−1^ for MBP proteins. Yields of the proteins are mentioned in **Table S4**.

All proteins were analyzed by mass spectrometry to determine the molecular weight and check for the presence of the ncAA residues. Prior to mass spectrometric analysis, proteins were desalted using an OPTI-TRAP Protein cartridge (Optimize Technologies). Samples were analyzed on a 15T solariX XR Fourier Transform Ion Cyclotron Resonance (FT-ICR) mass spectrometer (Bruker Daltonics) equipped with an Apollo II ESI ion source operated in positive ion mode. The samples were introduced into the ion source by direct infusion. Data were acquired with an acquisition size of 1 M and a mass range of 150–3000 m/z. The instrument was externally calibrated using sodium trifluoroacetate. Spectral data were processed using DataAnalysis 6.1 (Bruker Daltonics) with the SNAP algorithm and using UniDec software. ^74^

Mass spectrometry of the crosslinked EL222 was carried out with a bottom-up approach by digesting the proteins with trypsin and further analyzed by LC-MS/MS – timsTof Pro (Bruker Daltonics). DataAnalysis 6.1 (Bruker Daltonics) software was used to process the data.

## 3. Reactivity studies

### 3.1. Reactivity of DAF/ETF towards different functional groups

To test the reactivity of DAF towards thiol (-SH), di-sulfide (-S-S), carboxylic (-COOH), hydroxyl (-OH), azide (-N3), tetrazine and other noncanonical amino acids bearing either linear (-C≡C-) or cyclic alkyne (-C≡C-), and tetrazine groups, total twenty-one compounds (mentioned in following table) were selected. DAF was dissolved either in 100% Methanol or 100% DMSO. Benzyl azide, ETF, and Tet.2-Et, were dissolved in 100% DMSO whereas BCNK and SCOK were dissolved in 0.2 M NaOH + 15% DMSO. DTT, BME, Cysteine, Selenocysteine, Glutathione reduced, Glutathione oxidized, TCEP, Aspartic acid, Glutamic acid, Histidine, Methionine, Serine, and Tyrosine were dissolved in milliQ water. H-Tetrazine amine and Methyl-Tetrazine amine were dissolved in 100% Methanol. To perform the reaction, 15 mM DAF (dissolved in Methanol for reactions with H-Tetrazine amine and Met-Tetrazine amine and dissolved in 100% DMSO for reactions with rest of the compounds) was added to 20 mM each compound followed by incubation for 20 hours at room temperature. In parallel, ETF was dissolved in 100% DMSO and 15mM ETF was reacted with 20mM each of DTT, BME, Benzyl azide, H-Tetrazine amine and Methyl-Tetrazine amine for 20 hours at room temperature. After that, they were checked by MS to identify presence of any reacted compounds. Samples were diluted 200 x in 50% acetonitrile + 1% acetic acid and analyzed using 15T solariX XR Fourier Transform Ion Cyclotron Resonance (FT-ICR) mass spectrometer (Bruker Daltonics) equipped with an Apollo II ESI ion source operated in positive ion mode. The samples were introduced into the ion source by direct infusion. Data were acquired with an acquisition size of 2 M and a mass range of 150–1200 m/z. The instrument was externally calibrated using sodium trifluoroacetate. Spectral data were processed using DataAnalysis 6.1 (Bruker Daltonics).

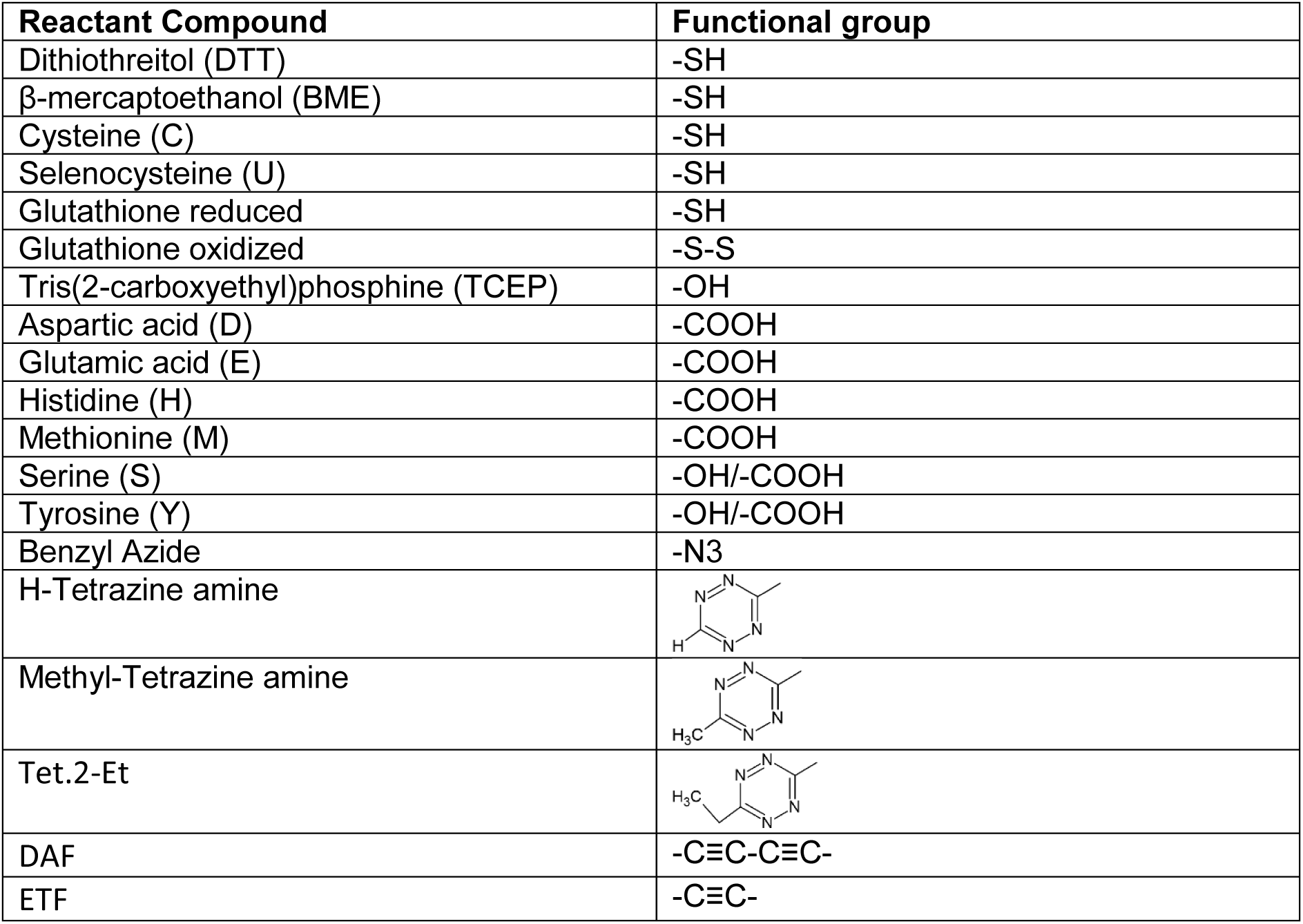
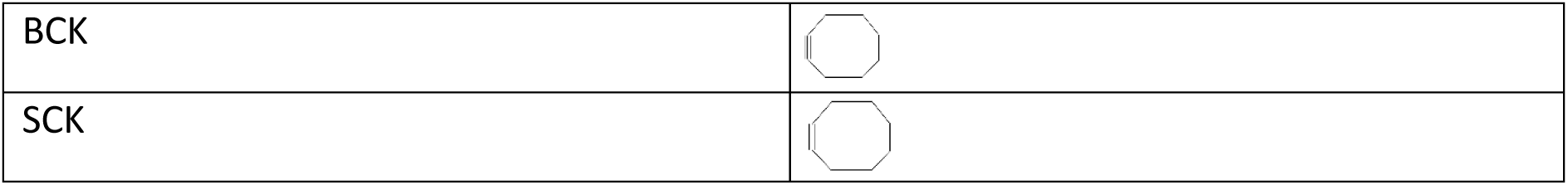

### 3.2. Set-up for steady-state fluorescence spectroscopy

The steady-state Fluorescence Spectroscopy to study either reaction kinetics or an intensity-based FRET measurement, was carried out using an Edinburgh Instruments FLS1000 spectrometer. The samples were excited by a 450 W ozone free continuous-wave xenon arc lamp providing a broad spectral range from 230 nm to >1000 nm wavelengths and the excitation beam with a desired wavelength was directed through a double grating Czerny-Turner monochromators with 325 mm (or 2 x 325 mm) focal length to gain high spectral purity, excellent stray light rejection and low temporal dispersion. Emission coming from the sample was collected in a right-angle detection geometry dispersed through a high-resolution emission monochromator followed by detection with red photomultiplier tube. Band-pass filter with 2 mm monochromator band width was employed to minimize higher-order diffraction and scattered excitation light. Polarizer was fixed at 55° angle with 100% light intensity. Dwell time was set to 0.5 s. All the measurements were conducted in 20°C temperature. Fluoracle 2.4.3 software was used for instrument control and data analysis.

### 3.3. Reaction kinetics between DAF/ETF and CalFluor Azide 647

To check the reaction kinetics of the reaction between DAF/ETF and fluorogenic dye CalFluor Azide 647,^75^ a concentrated stock of 100 mM DAF/ETF was prepared in Methanol. The dye was prepared in water. Kinetics reactions were performed by monitoring the change in fluorescence signal intensity over time at 20°C. For SPAAC reaction, kinetics measurements were started immediately after mixing 2 µM CalFluor azide dye and 200 µM DAF/ETF in PBS buffer pH=7.4 + 10% methanol and continued up to two hours for ETF reaction and seventeen hours for DAF reaction. CUAAC reaction required additional reagents.^75–77^ CuSO_4_ and THPTA stock solution was prepared in water. 2 µM CalFluor azide dye was added to a premixed solution of final concentration of 100 µM CuSO4 and 200 µM THPTA ligand in PBS buffer pH=7.4 + 10% methanol. The sodium ascorbate stock solution was prepared freshly and added at final concentration of 5 mM to initiate the CUAAC reaction and kinetics measurements were recorded immediately. Measurement time was followed same as the SPAAC reaction. Control experiment was done recording the kinetics of only 2 µM CalFluor azide dye in the same reaction solvent for seventeen hours.

### 3.4. Reaction kinetics between DAF/ETF/TCO and Atto488 H-Tetrazine

For reaction kinetics between either DAF, ETF or TCO with Atto488 H-Tetrazine dye,^78^ a concentrated stock of 100 mM of each ncAA was prepared in DMSO. The dye was dissolved in 100% DMSO. Kinetics reactions were initiated after mixing 2.5 µM of dye with 2.5 mM, either DAF or ETF and 2.5 µM TCO individually in 100% DMSO. Kinetics measurements were started immediately after reaction initiation and continued up to seventeen hours in case of DAF/ETF whereas two hours in case of TCO. Change in fluorescence signal intensity was monitored over time at 20°C. Similar to section .3., Control experiment was performed recording the kinetics of only 2.5 µM Atto488 H-Tetrazine dye in 100% DMSO for seventeen hours.

### 3.5. In-vitro labelling of MBP-V38DAF and MBP-V38ETF with Cy5 dyes

To check reactivity of MBP WT/DAF/ETF mutant proteins towards Cy5 dyes, 30 µM purified full length MBP proteins with N-terminal His tag and C-terminal Strep-tag II was labelled in 50 mM Tris, 100 mM NaCl buffer, pH 8.0 with 300 µM of either Cy5 H-Tetrazine or Cy5 Methyl Tetrazine, Cy5 Azide, or Cy5 Azide plus (a variant more reactive than its azide counterpart) in 50 µl reaction volume for 22h at room temp.^58,79^ Azide or Azide plus reactions were performed both with (CUAAC) and without (SPAAC) copper. For CUAAC labelling reaction, the following reagents were added in the order mentioned for setting up the labelling reaction: a)100 µM CuSO4, b) 200 µM THPTA ligand, c) 300 µM dye, d) 30 µM protein, and e) 500 µM freshly prepared sodium ascorbate in a total volume of 50 µl. After labeling, samples were directly loaded on a SDS-PAGE gel and analyzed for in-gel fluorescence on Azure c600 gel imager exciting with 650 nm laser and NIR detection filter settings. Afterwards, the gel was stained with Coomassie dye, destained and protein bands were analyzed by Azure 300 gel imager.

For the labelling of protein samples with Sun-Fluor SH-650, first 1 mM purified EL222 M151pDAF/EL222 M151pETF protein was labeled with 4 mM Thiol dye in 50 mM Tris, 100 mM NaCl buffer, pH 8.0 for 22h at room temperature in 50 µl reaction volume. The excess dye was removed by twice de-salting column chromatography using 5ml Zeba spin desalting column. The beads were washed and equilibrated thrice with 50mM Tris, 100mM NaCl, pH 8.0 buffer by centrifuging the column with 1000 g for 2 mins. The labelled protein was diluted to 700 µl and loaded into the beads. After the protein entered into the beads 100ul 50mM Tris, 100mM NaCl, pH 8.0 buffer was added as stacker. The column was centrifuged with 1000 g for 2 mins. The flow through was collected and again loaded onto a washed and equilibrated second column. The final flow through as a protein-dye adduct was collected. The UV-visible absorption spectra of the protein-dye adduct was checked. The concentration of the protein (based on extinction coefficient of FMN@450 nm) and concentration of dye (based on its extinction coefficient at its excitation wavelength, assuming ε642= 150000 M^-1^cm^-1^) present in adduct were calculated. The degree of dye labelling was found.

### 3.6. In-cell labelling of MBP V38DAF, V38ETF and controls

#### Preparation cell expressing protein

Cell samples were categorized based on plasmid content and ncAA supplementation. Cells lacking plasmids included BL21(DE3) cells grown either in standard TB medium or in TB medium supplemented with DAF or ETF. Cells harboring a single plasmid included: (i) BL21(DE3) cells transformed with plasmid pETH14F11-MBP, expressing wild-type MBP; (ii) cells transformed with pAJE-MaDafRS and cultured either in the presence or absence of DAF supplementation; and (iii) cells transformed with pDule2-CNF and cultured either in the presence or absence of ETF supplementation. Cells containing dual plasmids included: (i) BL21(DE3) cells co-transformed with plasmids pETH14F11-MBP_V38TAG and pAJE-MaDafRS, cultured either in the presence or absence of DAF supplementation; and (ii) cells co-transformed with plasmids pETH14F11-MBP_V38TAG and pDule2-CNF, and cultured either in the presence or absence of ETF supplementation.

BL21(DE3) cells were electro-transformed with either single or dual plasmids and plated on agar containing the appropriate antibiotics. Single colonies were inoculated into LB medium supplemented with antibiotics and grown overnight at 37 °C with shaking at 180 rpm. The following day, cultures were diluted 1% into 20 mL TB medium containing antibiotics and incubated at 37 °C with shaking until the optical density (OD₆₀₀) reached 0.5–0.6. Depending on the sample, DAF (0.5 M stock in methanol) or ETF (1 M stock in 3 M NaOH) was added to final concentrations of 0.5 mM and 1 mM, respectively. After 30 minutes, IPTG was added to induce protein expression, and all cultures were incubated for an additional 22 hours at 25 °C with shaking at 180 rpm.

#### Cell washing and storage

15 ml overnight cultures were centrifuged at 6000 g for 15 min at 4 °C to obtain cell pellets. The pellets were first resuspended in 30 mL of 0.1% methanol in PBS pH 7.4 incubated at 37 °C with shaking at 180 rpm for 30 min, followed by centrifugation at 6000 g for 5 min at 4 °C. The pellets were then washed twice by resuspension in 30 mL of 10% DMSO in PBS pH 7.4, each time incubating at 37 °C with shaking at 180 rpm for 30 min, followed by centrifugation at 6000 g for 5 min at 4 °C. Cells were subsequently resuspended in 10 mL PBS for OD₆₀₀ measurement. Appropriate volumes corresponding to OD₆₀₀ = 2 were aliquoted into microcentrifuge tubes and pelleted by centrifugation at 6000 g for 4 min at 4 °C. Finally, cell pellets were resuspended in 1 mL non-induce media (NIM)^80^ supplemented with 25% glycerol and stored at −80 °C until labeling.^66^

#### Cell labeling

Frozen E. coli aliquots were rapidly thawed in a warm water bath, followed by centrifugation at 6000 g for 4 minutes. The resulting cell pellet was resuspended in 1 ml PBS, followed by centrifugation at 6000 g for 4 minutes. The pellet was then resuspended in 1 mL NIM media in a 15 mL falcon tube and incubated at 37 °C with shaking at 180 rpm for 45 min, followed by centrifugation at 6000 g for 5 minutes. The cells were resuspended in 200 µL PBS and transferred into 800 µL PBS in an microcentrifuge tube, followed by centrifugation at 6000 × g for 4 minutes.

For fluorescent labeling, the cell pellet was resuspended in 100 µL of either 250 uM AzDye 488 Tetrazine in PBS; or 50 uM SunFluor SH488 in PBS; or 100 uM AZDye 488 Azide plus supplemented with 100 uM CuSO4, 200 uM THPTA, 500 uM sodium ascorbate final concentration in PBS. Cells with dye were incubated at 37 °C with shaking at 180 rpm for 30 minutes, followed by centrifugation at 6000 g for 4 minutes. For access dye washing out, cells were resuspended in 200 µL PBS, transfer into 800 µL PBS in microcentrifuge tube without incubation, followed by centrifugation at 6000 g for 4 minutes. Next, cells are resuspended in 200 µL PBS, transfer into 800 µL PBS in 15 ml falcon tube and incubated at 37 °C with shaking at 180 rpm for 30 min. After repeating the washing step one more time, the cells are resuspended in 40 ul PBS and ready for flow cytometry analysis.

#### Flow cytometry

Flow cytometric analysis was performed using a BD LSRFortessa™ SORP flow cytometer (BD Biosciences), and data were analysed using Kaluza Analysis Software version 2.0 (Beckman Coulter). Escherichia coli cells were fluorescently labelled with AZDye488 tetrazine, AZDye488 AzidePlus, or SunFluor488 SH, as described above. Fluorescence from the 488 nm-excitable dyes was excited with the 488 nm blue laser and collected through a 530/30 nm band-pass filter.

The bacterial population was initially identified on an FSC-H versus SSC-H dot plot. An empty medium sample was used to define background events and to distinguish the E. coli population from non-cellular particles and instrument noise. Subsequently, gated bacterial populations were visualized either as fluorescence histograms or as scatter versus fluorescence dot plots.

Positivity gates were set using the corresponding negative controls, and data were evaluated as the percentage of fluorescent-positive cells and as mean fluorescence intensity (MFI).

#### Confocal Fluorescence Microscopy

A 10 ul aliquot of bacteria suspension labeled with a 488 nm excitable fluorophore (AZDye488 and SunFluor488) was mounted on glass slides. Bacteria cells were imaged using Leica TCS SP8 confocal microscope (Leica Microsystems) equipped with a 63x/1.20 NA water-immersion objective (HC PL APO CS2) at room temperature. Samples were imaged in water immersion (refractive index 1.33). For confocal imaging scanning zoom was set to 15x. Bidirectional scanning was performed using harmonic scanner at frequency of 600 Hz. Fluorescence excitation was performed using the white light laser (WLL) tuned to 488 nm. Emission was collected over 500–650 nm using the hybrid/photomultiplier tube detector. Images were recorded at a resolution of 1024 × 1024 pixels with a pixel size of approximately 0.012 µm, 8-bit intensity depth. Line averaging was set to 16 to improve signal-to-noise ratio. The pinhole was set to 1 Airy unit. Z-stacks were acquired using system-optimized step sizes when applicable. Image processing and analysis were performed by Fiji software. Only linear adjustments of brightness and contrast of fluorescence channel were applied uniformly across all images (display range 0–255).

## 4. Front-illuminated surface plasmon resonance (fiSPR) biosensor

### 4.1. Set-up

The fiSPR biosensor used in this study was developed at the Institute of Photonics and Electronics (IPE), Prague, Czech Republic, as described in our previous work ^50^. The optical platform is based on the Kretschmann configuration and spectral modulation. In this configuration, a polychromatic light beam is collimated and directed onto the SPR chip *via* a prism coupler to excite surface plasmons. The SPR chip used in this study was obtained by depositing titanium (1.5 nm) and gold (50 nm) onto a glass substrate using electron-beam evaporation in vacuum. The sensor response is expressed as the shift in wavelength at which surface plasmon excitation occurs. This response is sensitive to changes in the refractive index caused by the binding of molecules to the surface of the SPR chip. For this biosensor, a sensor response of 1 nm corresponds to a protein surface density of approximately 17 ng/cm^2^. The above-described optical platform is integrated with a custom-designed microfluidic cartridge and a light module oriented orthogonally to the SPR chip from the front side, enabling *in situ* illumination of the analyzed sample. The microfluidic cartridge comprises a manifold with a central glass window that is adherent to the SPR chip using a double-sided adhesive gasket. A dispersionless microfluidic system and a specialized channel network are employed for sample injection. The channel network is organized in a detection and a reference channel, each of which presents a region used for the preliminary illumination of the sample (light-incubation area), and a region used for sensing (sensing area), where the sample flows under continuous illumination and interacts with the biomolecular receptors.^50^ The light module on the front side consists of a collimated light-emitting diode (LED), which is aligned with the central glass window of the microfluidic cartridge and emits at a wavelength of 450 nm with an intensity of 3.3 mW/cm². In all the fiSPR experiments, the volumetric flow rate and temperature were maintained at 20 μl/min and 25 °C, unless otherwise specified.

### 4.2. SPR chip functionalization

The light-incubation areas of the SPR chip were passivated with BSA, while the sensing areas were functionalized with the consensus double-stranded DNA (dsDNA), as reported in Ref. ^50^. Briefly, the gold surface was modified with a self-assembled monolayer (SAM) of alkanethiols, obtained by immersing the SPR chip overnight in an ethanolic solution of carboxy- and hydroxy-terminated alkanethiols with a total concentration of 200 µM and a molar ratio of 3:7. The SPR chip was washed with ethanol and Q water, dried with a stream of nitrogen and mounted into the fiSPR biosensor. The carboxylic groups were then activated *in situ* with an aqueous mixture of 25 mM NHS and 95.3 mM EDC for 10 minutes at 5 µl/min. To immobilize the proteins on the sensor surface, 50 μg/ml streptavidin and 100 μg/ml BSA diluted in 10 mM sodium acetate buffer, pH 5 were injected into the sensing and incubation areas, respectively, for 15 minutes. The non-covalently attached proteins were washed away by the injection of a phosphate buffer containing 1.4 mM KH_2_PO_4_, 8 mM Na_2_HPO_4_, 2.7 mM KCl, and 750 mM NaCl, pH 7.4 for 5 minutes. The unreacted carboxylic groups were deactivated by the injection of 0.5 M EA for 5 minutes. The DNA oligonucleotides were subsequently attached to the streptavidin immobilized in the sensing areas *via* streptavidin–biotin interactions. 10 nM biotinylated single-stranded (b-ssDNA) diluted in PBS was injected into the sensing areas of both detection and reference channels to reach a sensor response of ∼0.2 nm (corresponding to a surface density of ∼2 · 10^11^ b-ssDNA/cm^2^). To form the double-stranded DNA (dsDNA), a 40 nM solution of a complementary DNA sequence (cDNA) was injected into the sensing area of the detection channel until the maximum sensor response was reached (at approximately 10 minutes, with a surface density of ∼1.7 · 10^11^ dsDNA/cm^2^). All the DNA sequences used in this study are reported in **Supplementary Note S4**.

### 4.3. Interaction study of Interaction study of wild-type and cross-linked EL222 with dsDNA by fiSPR

To characterize the interaction of EL222 (including WT-EL222, and cross-linked EL222) with dsDNA, dsDNA20 was immobilized on the sensor surface of SPR. The cross-linked mutant of EL222 was compared to the EL222 WT under identical experimental conditions (using the same SPR chip). EL222 variants were injected into both detection and reference channels for 10 minutes at 1.4 μl/min. EL222 was flowed over the incubation areas of both channels for approximately 4 minutes, during which EL222 proteins were exposed to illumination from the front side prior to being injected into the sensing areas. For the on-chip comparison of EL222 WT and EL222 mutant, both proteins were diluted in 50 mM MES, 150 mM NaCl buffer pH 7.4 supplemented with 250 µg/ml BSA, to a concentration of 6 μM and injected into the detection and reference channels, both with and without *in situ* illumination supplied at a wavelength of 450 nm, a light intensity of 3.3 mW/cm^2^, and an illumination time of 260 s. For the kinetic characterization of crosslinked EL222–dsDNA interaction, protein was diluted in the same 50 mM MES, 150 mM NaCl buffer pH 7.4 supplemented with 250 µg/ml BSA to concentrations of 6, 8, 10, and 15 μM EL222 and injected into both detection and reference channels under same *in situ* illumination. All the sensor responses reported hereafter were reference-compensated by subtracting the sensor response of the reference channel from that of the detection channel (using Origin software, version 9.1). The kinetic fitting was performed using BIAevaluation software version 4.1, following the protocol detailed in our previous study.^50^ Briefly, assuming the same dimerization affinity for both EL222 variants, the total concentration of crosslinked EL222 dimers was calculated at equilibrium ([D]_eq_) and applied the 1:1 Langmuir model to fit the sensor responses to the binding of [D]_eq_ to dsDNA on the sensor surface. Additional SPR experiments are shown in **Supplementary Note S14**.

## 5. FTIR spectroscopy

### 5.1. Set-up

The stationary infrared spectra of the protein samples were recorded using a Bruker Vertex 70v FTIR spectrometer equipped with a globar source, KBr beam splitter and a liquid nitrogen cooled mercury cadmium telluride (MCT) detector. During FTIR spectral measurements the aperture was adjusted to obtain the maximum signal intensity while preventing detector saturation.^9^

### 5.2. Steady-state FTIR Spectroscopy with EL222 WT and L35DAFV217C mutant

Following the protocol as described in ^9^, both EL222 WT (WT-EL222) and cross-linked mutant (XL-EL222) prepared in H_2_O buffer (50 mM MES, 100 mM NaCl, pH 6.8) were transferred to D_2_O buffer (50 mM MES, 100 mM NaCl, pD 6.8) by centrifugal concentration method using 10 kDa cutoff filters. 30 μL of 1 mM each sample was loaded into a temperature controllable demountable liquid cell with CaF2 windows and a 50 μm Teflon spacer. For each sample, three spectra (dark state, lit state and difference) were recorded at 20°C and 200–300 scans were averaged in the spectral range of 4,000 to 900 cm^-1^ with a resolution of 2 cm^-1^. To measure the proteins in lit state the samples were irradiated by 450 nm LED with power of 25 mW/cm^2^ on throughout the measurement. Dark and lit samples were measured against buffer, and the difference spectra were recorded as the lit spectra against the dark spectra. The resulting spectra were processed with Spectra-Gryph and OriginPro (OriginLab 2024).

## 6. FRET studies

### 6.1. Set-up for Time-Resolved Fluorescence Spectroscopy

Experimental Fluorescence Lifetime spectra of the samples were measured using FLS1000 Photoluminescence Spectrometer with a time-correlated single photon counting (TCSPC) module. Excitation was provided by SuperK EXTREME, a supercontinuum white light laser (NKT Photonics) delivering broadband picosecond pulse from which 450 nm excitation wavelength was chosen using an excitation monochromator SuperK VARIA with a band width 2 mm. The repetition rate of 15.58 MHz was fixed with a pulse peaker and power intensity was 100%. The emitted photons from the proteins were collected in a 90° detection geometry, passed through a high-resolution emission monochromator for 528 nm and detected using red photomultiplier tube detector. Band-pass filter with 2 mm monochromator was used and polarizer was fixed at 55° angle as described in previous section. The time delay between excitation and emission was recorded to ten thousand counts by time-correlated single photon counting (TCSPC) to construct the histogram of photon arrival times. Time range and channels were set to 50 ns and 2048 respectively. All measurements were carried out in the single-photon counting regime to avoid pile-up effects. Fuoracle 2.4.3 software was used during measurements and analysis for tail decay fitting.

### 6.2. FRET with EL222 I225DAF

For FRET experiments, EL222 I225DAF-594 Tetrazine was prepared by labelling EL222 I225DAF with AZDye 594 Tetrazine. Initially, 1 mM unlabeled protein was labelled in 50 mM Tris, 100 mM NaCl buffer, pH 8.0 with 5 mM dye for 24 hours and labelling reaction was continued with additional 2 mM dye for another 24 hours. After a total of 48 hours of reaction, the excess dye was removed by loading the sample onto a 5ml Zeba spin desalting column equilibrated with 50 mM Tris, 100 mM NaCl buffer, pH 8.0 as described in section 4.5. The flow through was collected. This de-salting column chromatography was repeated once more with the flow through to ensure the complete excess dye removal. The flow through from the 2^nd^ desalting column as the protein-dye adduct was checked by the UV-visible absorption spectroscopy. The concentration of the protein (For EL222, based on extinction coefficient of FMN at 450 nm) and concentration of dye (Based on its extinction coefficient at its excitation wavelength, assuming ε590= 92000 M^-1^cm^-1^) present in adduct were calculated. The degree of dye labelling was found 76%.

For lifetime-based measurements, the donor protein sample EL222^WT^ and donor + acceptor protein sample EL222-I225DAF-594Tetrazine prepared in H_2_O buffer (50 mM Tris, 100 mM NaCl, pH 8.0) were transferred to D_2_O buffer (50 mM Tris, 100 mM NaCl, pD 8.4) by centrifugal concentration method using the 10 kDa cutoff filters. 60 µL of 15.5 µM each sample was loaded into a quartz cuvette with 3 mm path length. Decay curves were recorded at 20°C exciting the FMN within the protein samples. The decays corresponding to the “dark” state were measured in the absence of external illumination (apart from the laser source used to excite the FMN, which we found not be powerful enough to drive EL222 into the lit state). Simultaneous LED excitation and TCSPC data acquisition were found incompatible. Therefore, to capture the lit state, four different strategies were followed. First, the samples were irradiated by 450 nm LED with power of 35 mW/cm^2^ for 1 minute and then the measurement was started immediately measurement. Since the time required to acquire 10000 photon counts at the top of the decay curve was approximately 250 seconds, this condition is called *lit 250 s*. For the rest of the three strategies we call interval irradiation, samples were irradiated with 455 nm LED (35 mW) for 1 minute, measurements were started and after each 40 s/20 s/10 s interval, light was irradiated for 1 min/30 s/30 s respectively till 100% completion of the measurements. These samples are named *lit 40 s*, *lit 20 s*, and *lit 10 s*, respectively. The instrument response function (IRF) was measured separately using Ludox with 1000 times dilution in the same D_2_O buffer used for samples in scattering mode. The analysis of the decay curves is detailed in **Supplementary Note S15**.

## 7. Femtosecond-stimulated Raman Spectroscopy (FSRS)

### 7.1. Set-up

The FSRS spectroscopy experiment was conducted on a homebuilt setup constructed around a femtosecond Titanium sapphire amplifier, Femtopower (Spectra Physics), generating 4.2 mJ pulses of ∼20 fs duration at a repetitive rate of 1 kHz. Another Solstice amplifier (Spectra Physics) shares the fs oscillator. Both the Amplifiers were synchronized by electronic and optical delay to seed before amplification. The experiment is based on controlled time overall of three pulses in the sample, denoted as Probe (Probing transient absorption and Raman transition), Raman pump (Rp driving sample into vibration coherence with the Probe), and actinic pump (Ap – trigger the desired photoreaction).

The probe pulse is generated by splitting the laser output of 0.4 mJ of the laser and used to pump a two-stage optical parametric amplifier to generate 1450 nm pulses of ∼40 fs duration. This output was used as a pump for a single filament supercontinuum generated by a 2 mm CaF2 plate, resulting in a white light with the spectrum being from 370 nm to 1700 nm, in the near infrared region. The wavelength 1450 nm was chosen to have an undesired spike in the probe intensity from the white light driving the pump, matching the first peak of water infrared absorption. A 1 cm cuvette with water was used as an efficient notch filter, thus removing the 1450 nm spike, resulting in a flat white light covering the whole sensitivity range of the CCD detector. The Probe was imaged onto the sample by a spherical mirror to a 50 µm diameter spot, from the sample to the detection apparatus.

In the detection apparatus, the Probe is split into two beams, where one part is sent to a pair of homebuilt grating-based high-resolution imaging spectrographs for Raman analysis in the Stokes region 750 nm to 950 nm and anti-Stokes region 650 nm to 850 nm. The other part was directed to the prism spectrograph to obtain transient absorption (TA) spectra in 370 nm to 1200 nm range. In all three spectrographs, a 58×1024 pixel CCD (Entwicklungsbuero Stresing) was used as a linear image sensor via operation in a vertical-binning mode. Cameras were triggered from the lasers at 1 kHz and provided complete shot-to-shot detection with a dynamic range exceeding 30000:1. Despite the low intensity of the single filament supercontinuum (∼pJ/nm), it was possible to saturate the dynamic range of the sensors thoroughly. At the saturation level, the readout noise of this detector is two orders of magnitude lower than optical shot noise at a given intensity, so all presented measurements can be considered only the optical shot noise limit.

The Raman pump was generated from ∼1.5mJ of the laser output transmitted via a 4f pulse shaper, where the spherical disc with 96 shifted apertures was spinning at 10 Hz synchronized with the laser. At a time, 15 cm^-1^ intervals of wavelength were transmitted by each aperture, allowing for the production of 96 Raman pulses for each 100 incoming pulses, where each one was shifted by 5 cm^-1^ from each other. The shifted signals are then numerically combined to reduce fix pattern noise and facilitate baseline correction. The Raman pulses were generated in the interval from 770 to 795 nm, and the resulting Raman spectra represents an average of the signal from all Raman experiments conducted over this interval. Four pulses out of 100 were fully blocked to produce a pure transient absorption sequence along with the Raman experiment, leading to the cyclic scheme where 96% of the time the FSRS signal is measured and 4% of the time a pure TA signal is measured. The Raman pulses were guided via an optical delay line and then focused by a lens to ∼100 µm diameter spot overlapped with the Probe. Temporal overlap between Raman pump and Probe was adjusted to achieve maximal stimulated Raman gain while maintaining good spectral resolution. Average Raman pulse energy at the sample was ∼10-11 µJ.

The actinic pulse was generated from 1.5 mJ of the laser output, pumping a two-stage OPA combined with sum frequency generation (TOPAS, Light Conversion). The 475-490 nm output depending on different samples was guided via a motorized optical delay line and focused on the sample via a lens. The actinic pulse energy at the sample was adjusted from 1.15 µJ to 2 µJ.

### 7.2. Steady-state Raman Spectroscopy with non-canonical amino acids (ncAAS)

100 mM DAF was prepared dissolving it in 100% methanol and Stationary Raman spectra were measured in concentration dependent manner from 100 mM to 0.125 mM. The Raman signal intensity of 100 mM DAF was compared with same concentration of EDU as a reference compound and nine other noncanonical amino acids (ncAAs) bearing alkyne or azide moieties. These noncanonical amino acids are CNF, ETF, AZF, PRY, AZK, PRK, SCK, BCK and HPG. EDU was dissolved in milliQ water and heated up at 70°C, CNF was dissolved in milliQ water, ETF, AZF, PRY, AZK, PRK and HPG were dissolved in 3 M NaOH while SCK and BCK were dissolved in 0.2 M NaOH + 15% DMSO. To observe the response of the C≡C and C≡C-C≡C vibrations to the effect of diverse environments, vibrational solvatochromism studies on ETF and DAF respectively were implemented in solvents of different polarity. These two ncAAs were individually dissolved in different DMSO:Water mixtures and their Raman spectra were measured. In such studies the main parameters that were tracked are the mean stretching peak position and change in peak areas.

### 7.3. Steady-state Raman Spectroscopy with EL222 DAF/ETF mutants

The static Raman spectra of all the EL222 variants in 50 mM MES, 100 mM NaCl, pH 6.8 buffer were measured in a 2 mm quartz cuvette at room temperature recording two spectra for each sample, one in dark state and another in light state. The concentration of all the proteins measured in static FSRS is mentioned in the following table. Both the spectra were recorded in the spectral range of 3000 to 200 cm^-1^. The Protein lit state was induced by illuminating blue light of 455 nm LED (purchased from Thorlabs, M455L3) with power of 20 mW/cm^2^ on the sample throughout the measurement.

EL222 L35DAF was measured in denaturing condition as well after mixing it to urea with final concentration of protein and urea 0.9 mM and 4 mM respectively.

The resulting spectra were processed and difference spectra between either lit and dark states or non-denaturing and denaturing conditions were obtained with MATLAB and OriginPro (OriginLab).

### 7.4. Time-resolved (TR) Raman Spectroscopy with EL222 DAF/ETF mutants

Time resolved Raman spectra of EL222 mutants were recorded for dark to lit state transition with a time resolution of tens of milliseconds were recorded on the spectrophotometer as described in section 7.3. During measurements actinic pump was used to accumulate the lit state. Concentration of each sample is mentioned in **Table S4**. Spectra were taken with 75 delays. Raman spectra were measured both at Amide (fingerprint) region from 950-1750 cm^-1^ and Alkyne region from 1751-2500 cm^-1^.

## 8. Conventional Raman spectroscopy

### 8.1. Set-up

Raman microscope (InVia Renishaw) was set up with 532 nm, 50x objective, 30 mW, 1 s integration time, 30 accumulations and spectra center 2100 cm^-1^ (transparent window region of Raman spectrum). The laser was focused a few microns below the cover slip (thus a few microns into the sample volume).

### 8.2. In-cell Raman imaging

Suspensions of *E. coli* cells over-expressing MBP-DAF mutant protein and its negative control (*E. coli* cells over expressing MBP without DAF) were measured in two batches. The first batch was measured directly in the NB medium. The second batch was centrifuged, NB medium discarded, and resuspended in PBS. The suspension was put into a chamber made of microscope slide, spacer, and cover slip. 5 Raman spectra were collected for each sample. Spectra were processed using Savitzky-Golay algorithm (order 2, frame length 7), rolling circle filtering (radius 100 cm-1, passes 10), averaging per sample, and normalization at 1656 cm^-1^ (Amide I band).

The Raman results are detailed in **Supplementary Note S20**.

## References

1 Kottke, T., Xie, A., Larsen, D. S. & Hoff, W. D. Photoreceptors Take Charge: Emerging Principles for Light Sensing. Annual Review of Biophysics 47, 291–313, doi:10.1146/annurev-biophys-070317-033047 (2018).

2 Nash, A. I. et al. Structural basis of photosensitivity in a bacterial light-oxygen-voltage/helix-turn-helix (LOV-HTH) DNA-binding protein. Proceedings of the National Academy of Sciences 108, 9449–9454, doi:10.1073/pnas.1100262108 (2011).

3 Conrad, K. S., Manahan, C. C. & Crane, B. R. Photochemistry of flavoprotein light sensors. Nature Chemical Biology 10, 801–809, doi:10.1038/nchembio.1633 (2014).

4 Iuliano, J. N. et al. Variation in LOV Photoreceptor Activation Dynamics Probed by Time-Resolved Infrared Spectroscopy. Biochemistry 57, 620–630, doi:10.1021/acs.biochem.7b01040 (2018).

5 Liu, Y. et al. Sub-Millisecond Photoinduced Dynamics of Free and EL222-Bound FMN by Stimulated Raman and Visible Absorption Spectroscopies. Biomolecules 13, doi:10.3390/biom13010161 (2023).

6 Andrikopoulos, P. C. et al. QM calculations predict the energetics and infrared spectra of transient glutamine isomers in LOV photoreceptors. Physical Chemistry Chemical Physics 23, 13934–13950, doi:10.1039/d1cp00447f (2021).

7 Herzog, R. E. et al. Evolution and design shape protein dynamics in LOV domains – spanning picoseconds to days. Journal of Molecular Biology 438, doi:10.1016/j.jmb.2025.169599 (2026).

8 Chaudhari, A. S. et al. Light-dependent flavin redox and adduct states control the conformation and DNA-binding activity of the transcription factor EL222. Nucleic Acids Research 53, doi:10.1093/nar/gkaf215 (2025).

9 Chaudhari, A. S. et al. Genetically encoded non-canonical amino acids reveal asynchronous dark reversion of chromophore, backbone, and side-chains in EL222. Protein Science 32, doi:10.1002/pro.4590 (2023).

10 Herzog, R. E. et al. Signal propagation in LOV-based multidomain proteins: time-resolved infrared spectroscopy reveals the complete photocycle of YF1 and PAL. Physical Chemistry Chemical Physics 28, 2847–2857, doi:10.1039/d5cp03982g (2026).

11 Bannister, S., Böhm, E., Zinn, T., Hellweg, T. & Kottke, T. Arguments for an additional long-lived intermediate in the photocycle of the full-length aureochrome 1c receptor: A time-resolved small-angle X-ray scattering study. Structural Dynamics 6, doi:10.1063/1.5095063 (2019).

12 Kim, C. et al. Structural dynamics of protein-protein association involved in the light-induced transition of Avena sativa LOV2 protein. Nature Communications 15, doi:10.1038/s41467-024-51461-z (2024).

13 Davis, L. & Chin, J. W. Designer proteins: applications of genetic code expansion in cell biology. Nature Reviews Molecular Cell Biology 13, 168–182, doi:10.1038/nrm3286 (2012).

14 Fischer, S., Natter Perdiguero, A., Lau, K. & Deliz Liang, A. Hydrophobic tuning with non-canonical amino acids in a copper metalloenzyme. Nature Chemistry, doi:10.1038/s41557-026-02116-7 (2026).

15 Jann, C., Giofré, S., Bhattacharjee, R. & Lemke, E. A. Cracking the Code: Reprogramming the Genetic Script in Prokaryotes and Eukaryotes to Harness the Power of Noncanonical Amino Acids. Chemical Reviews 124, 10281–10362, doi:10.1021/acs.chemrev.3c00878 (2024).

16 Lang, K. & Chin, J. W. Bioorthogonal Reactions for Labeling Proteins. ACS Chemical Biology 9, 16–20, doi:10.1021/cb4009292 (2014).

17 Liu, X. et al. Excited-state intermediates in a designer protein encoding a phototrigger caught by an X-ray free-electron laser. Nature Chemistry 14, 1054–1060, doi:10.1038/s41557-022-00992-3 (2022).

18 Yu, B. et al. De novo design of light-responsive protein–protein interactions enables reversible formation of protein assemblies. Nature Chemistry 17, 1910–1919, doi:10.1038/s41557-025-01929-2 (2025).

19 Smits, A. H., Borrmann, A., Roosjen, M., van Hest, J. C. M. & Vermeulen, M. Click-MS: Tagless Protein Enrichment Using Bioorthogonal Chemistry for Quantitative Proteomics. ACS Chemical Biology 11, 3245–3250, doi:10.1021/acschembio.6b00520 (2016).

20 Saca, V. R., Mattheisen, J. M., Huber, T. & Sakmar, T. P. in *Epitope Mapping Protocols Methods in Molecular Biology* Ch. Chapter 12, 203–215 (2025).

21 Hall, C. R. et al. Site-Specific Protein Dynamics Probed by Ultrafast Infrared Spectroscopy of a Noncanonical Amino Acid. The Journal of Physical Chemistry B 123, 9592–9597, doi:10.1021/acs.jpcb.9b09425 (2019).

22 Ye, S., Huber, T., Vogel, R. & Sakmar, T. P. FTIR analysis of GPCR activation using azido probes. Nature Chemical Biology 5, 397–399, doi:10.1038/nchembio.167 (2009).

23 Ye, S. et al. Tracking G-protein-coupled receptor activation using genetically encoded infrared probes. Nature 464, 1386–1389, doi:10.1038/nature08948 (2010).

24 Krause, B. S. et al. Tracking Pore Hydration in Channelrhodopsin by Site-Directed Infrared-Active Azido Probes. Biochemistry 58, 1275–1286, doi:10.1021/acs.biochem.8b01211 (2019).

25 Kurttila, M. et al. Site-by-site tracking of signal transduction in an azidophenylalanine-labeled bacteriophytochrome with step-scan FTIR spectroscopy. Physical Chemistry Chemical Physics 23, 5615–5628, doi:10.1039/d0cp06553f (2021).

26 La Greca, M. et al. Propagation of Photoinduced Electric Field Changes Through Phytochrome and their Impact on Conformational Transitions. ChemPhysChem, doi:10.1002/cphc.202500595 (2025).

27 Kraskov, A. et al. Local Electric Field Changes during the Photoconversion of the Bathy Phytochrome Agp2. Biochemistry 60, 2967–2977, doi:10.1021/acs.biochem.1c00426 (2021).

28 Gil, A. A. et al. Photoactivation of the BLUF Protein PixD Probed by the Site-Specific Incorporation of Fluorotyrosine Residues. Journal of the American Chemical Society 139, 14638–14648, doi:10.1021/jacs.7b07849 (2017).

29 Gil, A. A. et al. Mechanism of the AppABLUF Photocycle Probed by Site-Specific Incorporation of Fluorotyrosine Residues: Effect of the Y21 pKa on the Forward and Reverse Ground-State Reactions. Journal of the American Chemical Society 138, 926–935, doi:10.1021/jacs.5b11115 (2016).

30 Moldenhauer, M. et al. Parameterization of a single H-bond in Orange Carotenoid Protein by atomic mutation reveals principles of evolutionary design of complex chemical photosystems. Frontiers in Molecular Biosciences 10, doi:10.3389/fmolb.2023.1072606 (2023).

31 Shi, W. & Lei, A. 1,3-Diyne chemistry: synthesis and derivations. Tetrahedron Letters 55, 2763–2772, doi:10.1016/j.tetlet.2014.03.022 (2014).

32 Wang, H., Du, J., Lee, D. & Wei, L. in Stimulated Raman Scattering Microscopy 289–310 (2022).

33 Qian, N. et al. Illuminating life processes by vibrational probes. Nature Methods 22, 928–944, doi:10.1038/s41592-025-02689-0 (2025).

34 Bae, K., Zheng, W., Ma, Y. & Huang, Z. Real-Time Monitoring of Pharmacokinetics of Mitochondria-Targeting Molecules in Live Cells with Bioorthogonal Hyperspectral Stimulated Raman Scattering Microscopy. Analytical Chemistry 92, 740–748, doi:10.1021/acs.analchem.9b02838 (2019).

35 Lee, H. J. et al. Assessing Cholesterol Storage in Live Cells and C. elegans by Stimulated Raman Scattering Imaging of Phenyl-Diyne Cholesterol. Scientific Reports 5, doi:10.1038/srep07930 (2015).

36 Zhang, J. et al. Small Unnatural Amino Acid Carried Raman Tag for Molecular Imaging of Genetically Targeted Proteins. The Journal of Physical Chemistry Letters 9, 4679–4685, doi:10.1021/acs.jpclett.8b01991 (2018).

37 Cai, E. et al. Imaging specific proteins in living cells with small unnatural amino acid attached Raman reporters. The Analyst 149, 5476–5481, doi:10.1039/d4an00758a (2024).

38 Flynn, J. D., Gimmen, M. Y., Dean, D. N., Lacy, S. M. & Lee, J. C. Terminal Alkynes as Raman Probes of α-Synuclein in Solution and in Cells. ChemBioChem 21, 1582–1586, doi:10.1002/cbic.202000026 (2020).

39 Wei, L. et al. Live-cell imaging of alkyne-tagged small biomolecules by stimulated Raman scattering. Nature Methods 11, 410–412, doi:10.1038/nmeth.2878 (2014).

40 Watson, M. D. & Lee, J. C. Genetically Encoded Aryl Alkyne for Raman Spectral Imaging of Intracellular α-Synuclein Fibrils. Journal of Molecular Biology 435, doi:10.1016/j.jmb.2022.167716 (2023).

41 Chen, Y. et al. Novel Vibrational Proteins. Analytical Chemistry 96, 16481–16486, doi:10.1021/acs.analchem.4c01569 (2024).

42 Meldal, M. & Tornøe, C. W. Cu-Catalyzed Azide−Alkyne Cycloaddition. Chemical Reviews 108, 2952–3015, doi:10.1021/cr0783479 (2008).

43 Massi, A. & Nanni, D. Thiol–yne coupling: revisiting old concepts as a breakthrough for up-to-date applications. Organic & Biomolecular Chemistry 10, doi:10.1039/c2ob25217a (2012).

44 Young, D. D. et al. An Evolved Aminoacyl-tRNA Synthetase with Atypical Polysubstrate Specificity. Biochemistry 50, 1894–1900, doi:10.1021/bi101929e (2011).

45 Wang, Y.-S., Fang, X., Wallace, A. L., Wu, B. & Liu, W. R. A Rationally Designed Pyrrolysyl-tRNA Synthetase Mutant with a Broad Substrate Spectrum. Journal of the American Chemical Society 134, 2950–2953, doi:10.1021/ja211972x (2012).

46 Beránek, V., Willis, J. C. W. & Chin, J. W. An Evolved Methanomethylophilus alvus Pyrrolysyl-tRNA Synthetase/tRNA Pair Is Highly Active and Orthogonal in Mammalian Cells. Biochemistry 58, 387–390, doi:10.1021/acs.biochem.8b00808 (2018).

47 Cheng, L., Wang, Y., Guo, Y., Zhang, S. S. & Xiao, H. Advancing protein therapeutics through proximity-induced chemistry. Cell Chemical Biology 31, 428–445, doi:10.1016/j.chembiol.2023.09.004 (2024).

48 Xiang, Z. et al. Adding an unnatural covalent bond to proteins through proximity-enhanced bioreactivity. Nature Methods 10, 885–888, doi:10.1038/nmeth.2595 (2013).

49 Lorenz-Fonfria, V. A. Infrared Difference Spectroscopy of Proteins: From Bands to Bonds. Chemical Reviews 120, 3466–3576, doi:10.1021/acs.chemrev.9b00449 (2020).

50 Finocchiaro, G. et al. Front-illuminated surface plasmon resonance biosensor for the study of light-responsive proteins and their interactions. Biosensors and Bioelectronics 291, doi:10.1016/j.bios.2025.117998 (2026).

51 Zoltowski, B. D., Nash, A. I. & Gardner, K. H. Variations in Protein–Flavin Hydrogen Bonding in a Light, Oxygen, Voltage Domain Produce Non-Arrhenius Kinetics of Adduct Decay. Biochemistry 50, 8771–8779, doi:10.1021/bi200976a (2011).

52 Andrikopoulos, P. C. et al. Femtosecond-to-nanosecond dynamics of flavin mononucleotide monitored by stimulated Raman spectroscopy and simulations. Physical Chemistry Chemical Physics 22, 6538–6552, doi:10.1039/c9cp04918e (2020).

53 Konold, P. E. et al. Unfolding of the C-Terminal Jα Helix in the LOV2 Photoreceptor Domain Observed by Time-Resolved Vibrational Spectroscopy. The Journal of Physical Chemistry Letters 7, 3472–3476, doi:10.1021/acs.jpclett.6b01484 (2016).

54 Konold, P. E. et al. Microsecond time-resolved X-ray scattering by utilizing MHz repetition rate at second-generation XFELs. Nature Methods 21, 1608–1611, doi:10.1038/s41592-024-02344-0 (2024).

55 Bednar, R. M., Karplus, P. A. & Mehl, R. A. Site-specific dual encoding and labeling of proteins via genetic code expansion. Cell Chemical Biology 30, 343–361, doi:10.1016/j.chembiol.2023.03.004 (2023).

56 Plass, T. et al. Amino Acids for Diels-Alder Reactions in Living Cells. Angewandte Chemie International Edition 51, 4166–4170, doi:10.1002/anie.201108231 (2012).

57 Plass, T., Milles, S., Koehler, C., Schultz, C. & Lemke, E. A. Genetically Encoded Copper-Free Click Chemistry. Angewandte Chemie International Edition 50, 3878–3881, doi:10.1002/anie.201008178 (2011).

58 Nikić, I. et al. Minimal Tags for Rapid Dual-Color Live-Cell Labeling and Super-Resolution Microscopy. Angewandte Chemie International Edition 53, 2245–2249, doi:10.1002/anie.201309847 (2014).

59 Nguyen, D. P. et al. Genetic Encoding and Labeling of Aliphatic Azides and Alkynes in Recombinant Proteins via a Pyrrolysyl-tRNA Synthetase/tRNACUA Pair and Click Chemistry. Journal of the American Chemical Society 131, 8720–8721, doi:10.1021/ja900553w (2009).

60 Milles, S. et al. Click Strategies for Single-Molecule Protein Fluorescence. Journal of the American Chemical Society 134, 5187–5195, doi:10.1021/ja210587q (2012).

61 Blizzard, R. J. et al. Ideal Bioorthogonal Reactions Using A Site-Specifically Encoded Tetrazine Amino Acid. Journal of the American Chemical Society 137, 10044–10047, doi:10.1021/jacs.5b03275 (2015).

62 Eddins, A. J. et al. Quantitative Protein Labeling in Live Cells by Controlling the Redox State of Encoded Tetrazines. Journal of the American Chemical Society 147, 23625–23634, doi:10.1021/jacs.5c04605 (2025).

63 Kozma, E. et al. Hydrophilic trans-Cyclooctenylated Noncanonical Amino Acids for Fast Intracellular Protein Labeling. ChemBioChem 17, 1518–1524, doi:10.1002/cbic.201600284 (2016).

64 Kumar, G. S., Racioppi, S., Zurek, E. & Lin, Q. Superfast Tetrazole–BCN Cycloaddition Reaction for Bioorthogonal Protein Labeling on Live Cells. Journal of the American Chemical Society 144, 57–62, doi:10.1021/jacs.1c10354 (2021).

65 Eddins, A., Pung, A., Cooley, R. & Mehl, R. Tetrazine Amino Acid Encoding for Rapid and Complete Protein Bioconjugation. Bio-Protocol 14, doi:10.21769/BioProtoc.5048 (2024).

66 Ilievski, F. et al. Optimization of the genetic code expansion technology for intracellular labelling and single-molecule tracking of proteins in genomically re-coded E. coli. RSC Chemical Biology, doi:10.1039/d5cb00221d (2025).

67 Yee, E. F. et al. Signal transduction in light–oxygen–voltage receptors lacking the adduct-forming cysteine residue. Nature Communications 6, doi:10.1038/ncomms10079 (2015).

68 Kopka, B. et al. Electron transfer pathways in a light, oxygen, voltage (LOV) protein devoid of the photoactive cysteine. Scientific Reports 7, doi:10.1038/s41598-017-13420-1 (2017).

69 Iuliano, J. N. et al. Unraveling the Mechanism of a LOV Domain Optogenetic Sensor: A Glutamine Lever Induces Unfolding of the Jα Helix. ACS Chemical Biology 15, 2752–2765, doi:10.1021/acschembio.0c00543 (2020).

70 Yao, X., Rosen, M. K. & Gardner, K. H. Estimation of the available free energy in a LOV2-Jα photoswitch. Nature Chemical Biology 4, 491–497, doi:10.1038/nchembio.99 (2008).

71 Löffler, J. G., Deniz, E., Feid, C., Franz, V. G. & Bredenbeck, J. Versatile Vibrational Energy Sensors for Proteins. Angewandte Chemie International Edition 61, doi:10.1002/anie.202200648 (2022).

72 Deniz, E. et al. Through bonds or contacts? Mapping protein vibrational energy transfer using non-canonical amino acids. Nature Communications 12, doi:10.1038/s41467-021-23591-1 (2021).

73 Maximov, O. B. & Pantiukhina, L. S. Thin-layer partition chromatography of benzenecarboxylic and hydroxybenzenecarboxylic acids. Journal of Chromatography A 20, 160–162, doi:10.1016/s0021-9673(01)97382-0 (1965).

74 Marty, M. T. et al. Bayesian Deconvolution of Mass and Ion Mobility Spectra: From Binary Interactions to Polydisperse Ensembles. Analytical Chemistry 87, 4370–4376, doi:10.1021/acs.analchem.5b00140 (2015).

75 Shieh, P. et al. CalFluors: A Universal Motif for Fluorogenic Azide Probes across the Visible Spectrum. Journal of the American Chemical Society 137, 7145–7151, doi:10.1021/jacs.5b02383 (2015).

76 Presolski, S. I., Hong, V. P. & Finn, M. G. Copper-Catalyzed Azide–Alkyne Click Chemistry for Bioconjugation. Current Protocols in Chemical Biology 3, 153–162, doi:10.1002/9780470559277.ch110148 (2011).

77 Tyagi, S. & Lemke, E. A. in Laboratory Methods in Cell Biology - Imaging Methods in Cell Biology 169–187 (2013).

78 Hild, F., Werther, P., Yserentant, K., Wombacher, R. & Herten, D.-P. A dark intermediate in the fluorogenic reaction between tetrazine fluorophores and trans-cyclooctene. Biophysical Reports 2, doi:10.1016/j.bpr.2022.100084 (2022).

79 Mayer, S. V., Murnauer, A., von Wrisberg, M. K., Jokisch, M. L. & Lang, K. Photo-induced and Rapid Labeling of Tetrazine-Bearing Proteins via Cyclopropenone-Caged Bicyclononynes. Angewandte Chemie International Edition 58, 15876–15882, doi:10.1002/anie.201908209 (2019).

80 Alexander, N. D. et al. Selecting aminoacyl-tRNA synthetase/tRNA pairs for efficient genetic encoding of noncanonical amino acids into proteins. Nat Protoc 21, 1374–1428, doi:10.1038/s41596-025-01241-w (2026).

