## Supplementary information for "A chemically reactive and Raman-active non-canonical amino acid reveals photocycle complexity in a blue-light receptor"

Note S2. Chemical reactivity of DAF. ....15-33

Note S3. Raman spectroscopy of "transparent window" probes. ....34-36

Note S4. Sequences of all constructs used in the present study. ....37-39

Note S19. Analysis of time-resolved absorption datasets. ....66-67

Note S20. Raman microspectroscopy of DAF. ....68-69

Supplementary references....70-71

#### Note S1. DAF synthesis.

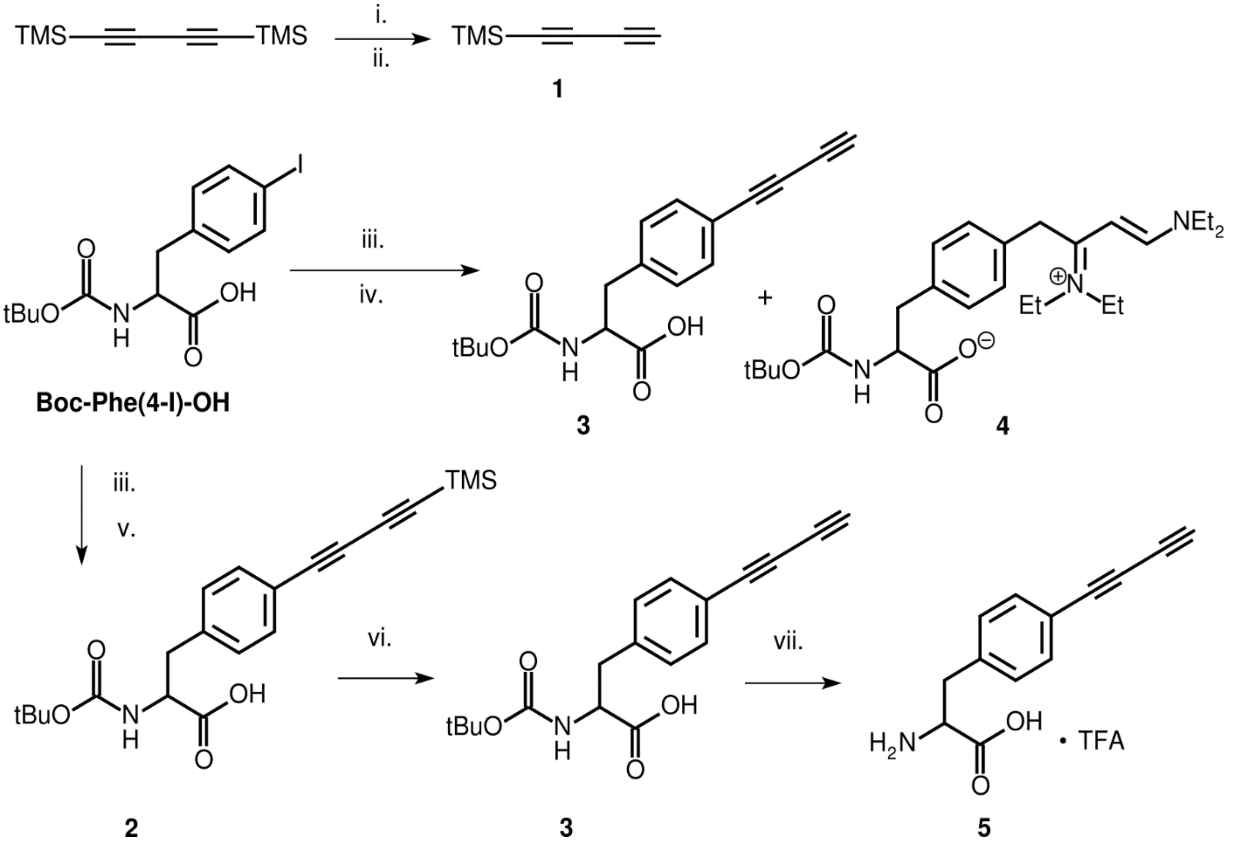

**Fig. S1. Synthesis of the novel ncAA DAF (5).**

i.) MeLi (3.1 M in $EtOCH_2OEt$), $Et_2O$. ii.) $NH_4Cl$; 53%. iii.) compound **1**, $PdCl_2(PPh_3)_2$, CuI, $PPh_3$, TEA, toluene, 60° C, overnight, 100% conversion. iv.) column chromatography EtOAc/DEA/MeOH (75:38:24) $SiO_2$; **3** 69%, **4** ~7%. v.) column chromatography $Et_2O$; ~100%. vi.) TBAF/$H_2O$/THF room temperature, 58 min, 97%. vii.) TFA/Anisole/DCM (25/8/19), 0° C, 46 min, 97%.

#### 1-(Trimethylsilyl)buta-1,3-diyn (1) (Compound characterization graph S1)

A published procedure[1] was slightly adapted using commercially available solution of MeLi. Briefly, to solid TMS-$C_4$-TMS (12.2 g, ~ 60 mmol), freshly dry ether (100 mL) was transferred by cannula using overpressure of nitrogen. The stirred solution was cooled down by bath at 0°C. Commercial solution of MeLi (3.1 M, 23 mL, 72 mmol, in $EtOCH_2OEt$) was added dropwise to reaction mixture. After complete addition, the reaction mixture was stirred overnight at room temperature. Next day, the reaction mixture was cooling down to -72°C by dry ice/ethanol bath, and ~60 mL of saturated $NH_4Cl$ solution (27 g $NH_4Cl$ in 61 g od water) was added dropwise. After addition, the reaction was allowed to warm up, and ether layer was separated. The aqueous phases were washed with ether (3×70 mL). Combined organic phases were dried with $MgSO_4$, filtered and gently evaporated 270 mbar/20°C. The residuum was fractionally distilled 130→72 mbar, bath temp. 22°C, #1; 72 mbar, bath temp. 22-60°C, #2 contaminated with water; 72 mbar, bath temp. 60-92°C, #3, 7.95 g, ~9.6 ml, ϱ 0.828 g/mL. For comparison, $EtOCH_2OEt$ has ϱ 0.831 g/mL. For NMR, 143 mg was diluted with 1.13 g of $CDCl_3$. NMR content of alkyne is 49% m/m. Yield 53% of colorless solution (**1**), which turned yellow upon standing. **$^1$H-NMR** (400 MHz, $CDCl_3$) δ 2.06 (s, 1H, ≡C-**H**), 0.12 (s, 9H, C**H**$_{3,TMS}$). $EtOCH_2OEt$ δ 4.58 (s, 2H, OC**H**$_2$O, ~1.22 × molar equivalent of the alkyne), 3.51 (q, J = 7.1 Hz, 2H, C**H**$_2$), 1.13 (t, J = 7.1 Hz, 3H, C**H**$_3$). **$^{13}$C-NMR** (100 MHz, $CDCl_3$) δ 87.54 (≡**C**-), 84.50 (≡**C**-TMS), 68.34 (≡**C**-), 66.76 (≡**C**-H), -0.56 (**C**H$_{3,TMS}$). $EtOCH_2OEt$ δ 94.88 (O**C**H$_2$O), 63.11 (**C**H$_2$), 15.21 (**C**H$_3$). Raman (785 nm, $CDCl_3$, $cm^{-1}$): ν(C≡C) 2184. The data are in accordance with those reported in the literature.[1] Fraction #2 was dried with $Na_2SO_4$ and filtered to vial. NMR content of alkyne 13.6% m/m. But it contained also some other impurities.

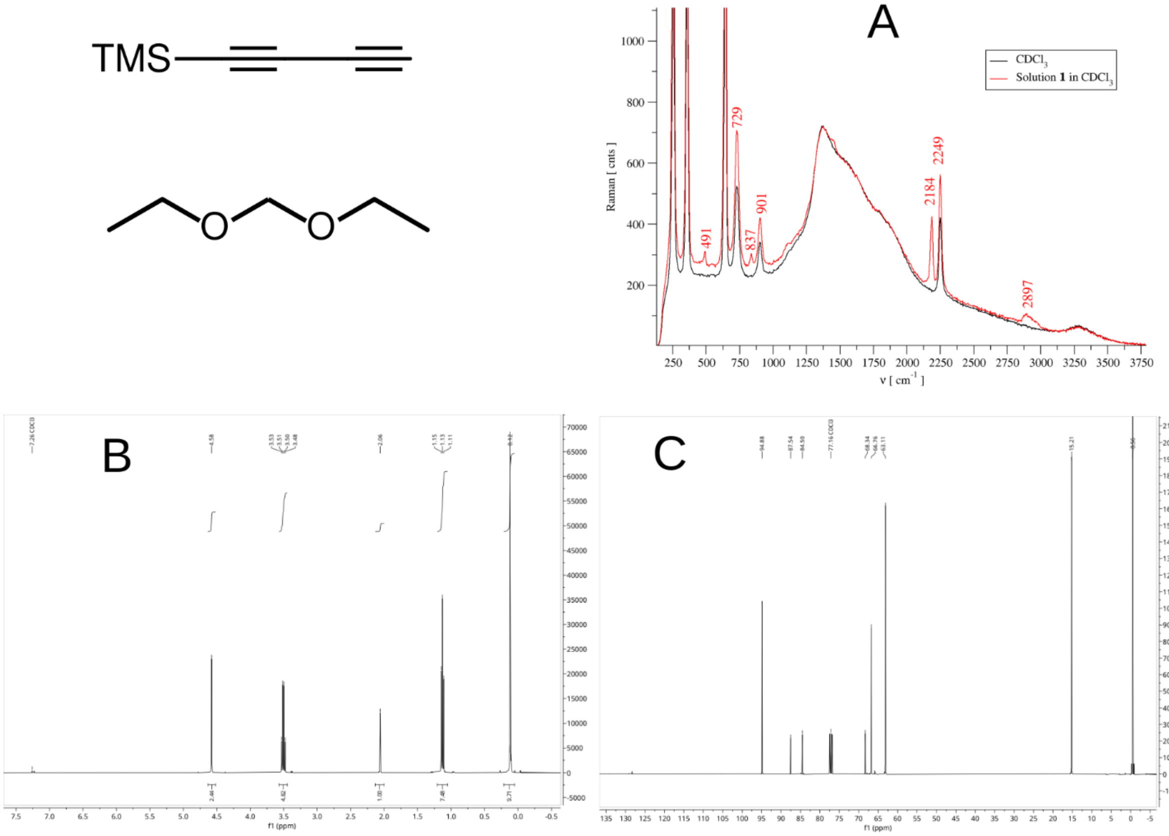

**Compound characterization graph S1.** Formulas of compound **1** (up) and its solvent (down). A) Raman spectra after NMR measurement. B) $^1$H-NMR and C) $^{13}$C-NMR spectra.

#### L-2-((*terc*-butoxycarbonyl)amino)-3-(4-((trimethylsilyl)buta-1,3-diyn-1-yl)phenyl)propanoic acid (2) (Compound characterization graph S2)

To solids Boc-Phe(4-I)-OH (1.3 g, 3.32 mmol), $PdCl_2(PPh_3)_2$ (206 mg, 0.29 mmol), CuI (66 mg, 0.35 mmol), $PPh_3$ (82 mg, 0.31 mmol) in nitrogen atmosphere, the freshly dried toluene (60 mL) was loaded by cannula using overpressure of nitrogen. At 65°C, the TMS-$C_4$H (**1**, 1.2 mL, 49% m/m in $(EtO)_2CH_2$, 3.98 mmol) and dry TEA (4.4 mL, 31.6 mmol) were added dropwise within 15 min. Reaction mixture was stirred for 24 h at 60°C. After cooling down, the reaction mixture was filtered through Celite. The flask was washed with addition toluene (30 mL and 60 mL). After first washing, the filtrate was colorless. Toluene was evaporated 30 mbar/24°C (beware, do not overheat sample to avoid significant formation of tars). The residuum was redissolved using EtOAc (25 mL) and 1M $KHSO_4$ (25 mL) and transferred to separation funnel. The flask was washed with additional EtOAc (10 mL) and 1M $KHSO_4$ (10 mL); and EtOAc (20 mL). Combined organic phases were washed 1M KHSO4 (2×25 mL), and dried with $Na_2SO_4$. After filtration, the EtOAc was evaporated (94 mbar/22°C), and dried for 1 h (3 mbar/22°C). The residuum was suspended in $Et_2O$ (10.5 mL) and filtered, the solid debris was washed with additional $Et_2O$ (6 mL). $Et_2O$ was evaporated (430 mbar/22°C) and the residuum dried (3 mbar/22°C) for 1h, 1.536 g. The debris from filter was dissolved in MeOH and evaporated to dryness, 47 mg. The residuum (1.536 g) was purified using column chromatography $SiO_2$ (81.6 g), $Et_2O$ (250 mL). It was dissolved in $Et_2O$ (5 mL) by help of sonication and led for awhile to separate insoluble part. Sample was loading to column carefully, to avoid use of settled parts. The solid was washed by additional $Et_2O$ (5 mL), and loaded to the column. The insoluble part was left to evaporated, 35 mg. Debris obtained before column chromatography (a) and during column loading (b) are insoluble in MeOH and acetone, for HPLC analysis they were dissolved in DMF. The column after soaking of soluble part was washed with $Et_2O$ (3×5 mL), and then chromatography with additional $Et_2O$ (305 mL) was carried out. Fractions were evaporated to dryness (300 mbar/25°C→5 mbar/24°C) and analyzed by TLC ($CHCl_3$:MeOH 5:1) and HPLC. #0 (30 mg); #3-8 (1.075 g); #9-15 (212 mg). #9-15 is more pure i.e. contained less tars visible at 254 nm. Combined fraction #3-15, yield 1.28 g (3.32 mmol, ~100%) of **2**. **TLC**: $R_f$ 0.35 ($CHCl_3$:MeOH 5:1). **HPLC**: $R_T$ 8.4 min. **ESI-HRMS** (m/z): for $[M+Na]^+$ $C_{21}H_{27}O_4NNaSi$ calcd. 408.16016; found 408.16052 (0.90 ppm); $[M-H]^-$ $C_{21}H_{26}O_4NSi$ calcd. 384.16366; found 384.16361 (-0.12 ppm). **$^1$H-NMR** (400 MHz, DMSO-*d6*) δ 12.65 (br, 1H, COO**H**), 7.49 (d, J = 8.2 Hz, 2H, **H**$3_{,Ph}$), 7.29 (d, J = 8.4 Hz, 2H, **H**$2_{,Ph}$), 7.14 (d, J = 8.5 Hz, 1H, CON**H**), 4.10 (td, J = 9.6, 4.2 Hz, 1H, **H**α), 3.05 (dd, J = 13.8, 4.5 Hz, 1H, **H**β), 2.84 (dd, J = 13.9, 10.5 Hz, 1H, **H**β), 1.30 (s, 9H, C**H**$_{3,Boc}$), 0.21 (s, 9H, C**H**$_{3,TMS}$). **$^{13}$C-NMR** (100 MHz, DMSO-*d6*) δ 173.31 (**C**OOH), 155.42 (**C**ONH), 140.68 (**C**$1_{,Ph}$), 132.43 (**C**$3_{,Ph}$), 129.72 (**C**$2_{,Ph}$), 117.86 (**C**$4_{,Ph}$), 91.04 (≡**C**-TMS), 87.94 (≡**C**-), 78.10 (**C**$_{Boc}$), 77.17 (≡**C**-Ph), 73.51 (≡**C**-), 54.73 (**C**α), 36.40 (**C**β), 28.12 (**C**H$_{3,Boc}$), -0.58 (**C**H$_{3,TMS}$). **IR** (film on $CaF_2$, $cm^{-1}$): ν(C≡C-C≡C)$_{symm}$ 2205, ν(C≡C-C≡C)$_{as}$ 2105, ν(C=O) 1718.

Computational model for explanation of two visible vibration of C≡C-C≡C system (Fig S3G). Top image shows one phase of these vibrations; whereas the bottom one shows another phase of the vibrations. At computed 2190 cm-1, one triple bond is shortening, whereas the second one is extending. At computed 2313 cm-1, both triple bonds are shortening or extending simultaneously.

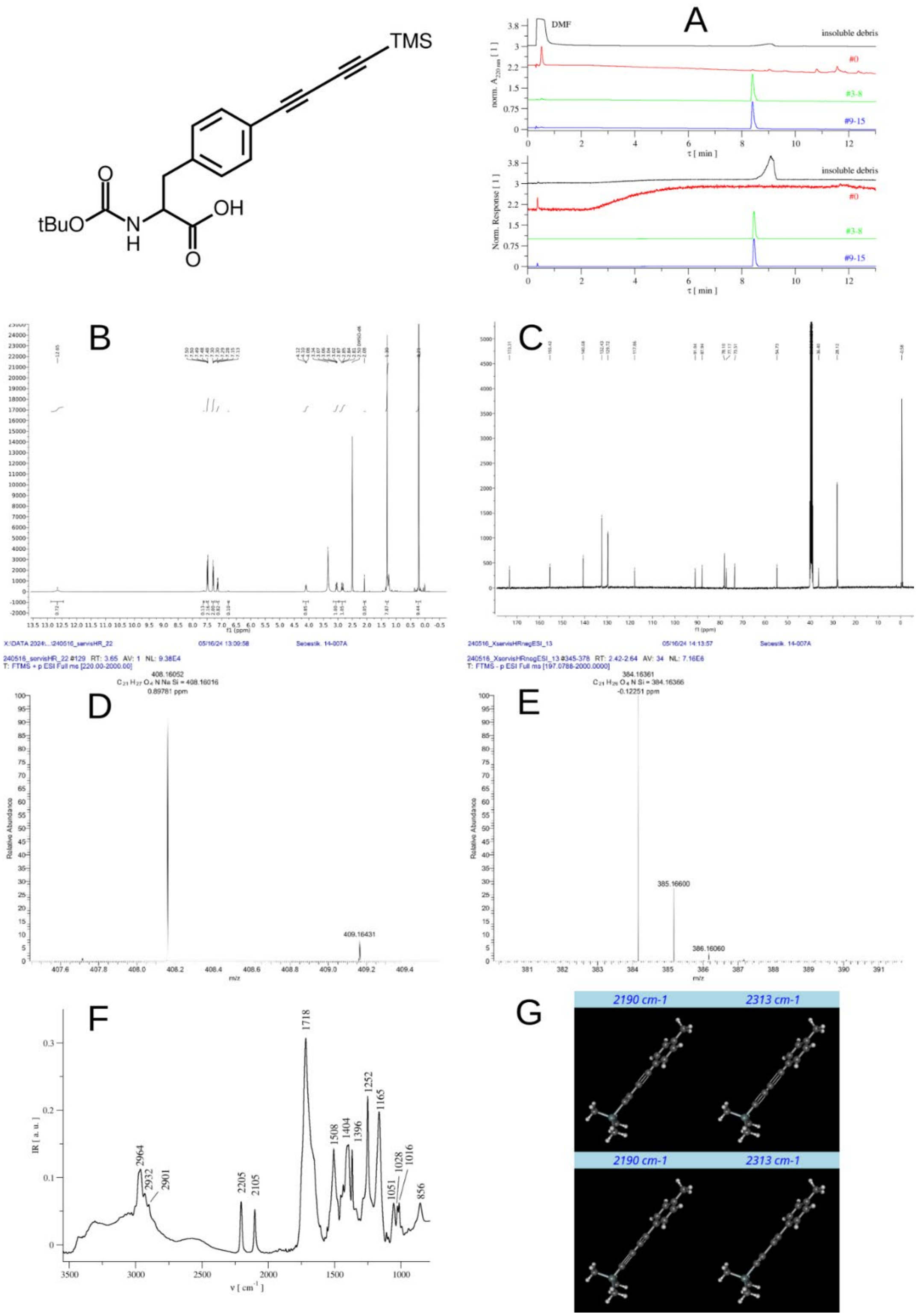

**Compound characterization graph S2.** Formula of compound **2**. A) Analytical HPLC records of fractions from column preparation of compound **2**. DAD detector (top) and ELSD (bottom). B) $^{1}$H-NMR spectrum. C) $^{13}$C-NMR spectrum. D) HRMS-spectrum. E) HRMS-spectrum. F) IR spectrum of compound **2**, film on $CaF_2$. G) DFT simulation of Ph-C≡C-C≡C-H vibrations.

##### L-3-(4- (buta-1,3-diyn-1-yl)phenyl-2-((*tert*-butoxycarbonyl)amino)propanoic acid (3) – method A (Compound characterization graph S3)

To solids Boc-Phe(4-I)-OH (497 mg, 1.27 mmol), $PdCl_2(PPh_3)_2$ (71.7 mg, 0.10 mmol), CuI (56 mg, 0.29 mmol), $PPh_3$ (32 mg, 0.12 mmol) in nitrogen atmosphere, the freshly dried toluene (20 mL) was loaded by cannula using overpressure of nitrogen. At 65°C, the TMS-$C_4$H (**1**, 450 µL, 49% m/m in $(EtO)_2CH_2$, 1.49 mmol) and dry TEA (1.42 mL, 10.2 mmol) were added dropwise within 15 min. Reaction mixture was stirred for 24 h at 60°C. After cooling down, the reaction mixture was filtered through Celite. The flask was washed with addition toluene (5 mL and 10 mL). After first washing, the filtrate was colorless. Toluene was evaporated 30 mbar/24°C (beware, do not overheat sample to avoid significant formation of tars). The residuum was redissolved using EtOAc (25 mL) and 1M $KHSO_4$ (25 mL) and transferred to separation funnel. The flask was washed with additional EtOAc (10 mL) and 1M $KHSO_4$ (10 mL); and EtOAc (20 mL). Combined organic phases were washed 1M $KHSO_4$ (2×25 mL), and dried with $Na_2SO_4$. After filtration, the EtOAc was evaporated (94 mbar/22°C), and dried for 1 h (3 mbar/22°C). Ca 620 mg of crude material, it was obtained. According to HPLC, it contained mostly compound **2**. The residuum (620 mg) was purified using column chromatography $SiO_2$ (43 g), EtOAc (375 mL), DEA (190 mL), and MeOH (120 mL) i.e. (75:38:24). The residuum was loaded on the column as EtOAc solution. The fractions were collected and evaporated to dryness, and dried for several hours (5 mbar/50°C): #13-15 (34 mg), #16-17 (107 mg), and #18-29 (274 mg). Fractions #18-29 were TLC pure. TLC $R_F$(EtOAc/DEA/MeOH 75:25:32) 0.41. Contamination with by-product **4** (detectable by HPLC) was removed by washing between EtOAc/1M $KHSO_4$. The organic phases were dried with $MgSO_4$, filtered and evaporated to dryness. Yield 136 mg (**3**, 34%). **HPLC**: $R_T$ 6.6 min. **ESI-HRMS** (m/z): for $[M+Na]^+$ $C_{18}H_{19}O_4NNa$ calcd. 336.12063; found 336.12085 (0.66 ppm). **$^1$H-NMR** (400 MHz, DMSO-*d6*) δ 12.65 (s, 1H, COO**H**), 7.50 (d, J = 7.9 Hz, 2H, **H**3,$_{Ph}$), 7.29 (d, J = 7.8 Hz, 2H, **H**2,$_{Ph}$), 7.13 (d, J = 8.4 Hz, 1H, CON**H**), 4.10 (ddd, J = 10.5, 8.5, 4.6 Hz, 1H, **H**α), 4.02 (s, 1H, ≡C-**H**), 3.05 (dd, J = 13.7, 4.6 Hz, 1H, **H**β), 2.84 (dd, J = 13.8, 10.5 Hz, 1H, **H**β), 1.30 (s, 9H, C**H**$_{3,Boc}$). **$^{13}$C-NMR** (100 MHz, DMSO-*d6*) δ 173.34 (**C**OOH), 155.43 (**C**ONH), 140.67 (**C**1,$_{Ph}$), 132.48 (**C**3,$_{Ph}$), 129.72 (**C**2,$_{Ph}$), 117.77 (**C**4,$_{Ph}$), 78.11 (**C**$_{Boc}$), 76.00 (≡**C**-H), 75.01 (≡**C**-Ph), 73.19 (≡**C**-), 67.69 (≡**C**-), 54.75 (**C**α), 36.40 (**C**β), 28.14 (**C**H$_{3,Boc}$). According to NMR data, the sample contained remaining solvents EtOAc and DEA from chromatography.

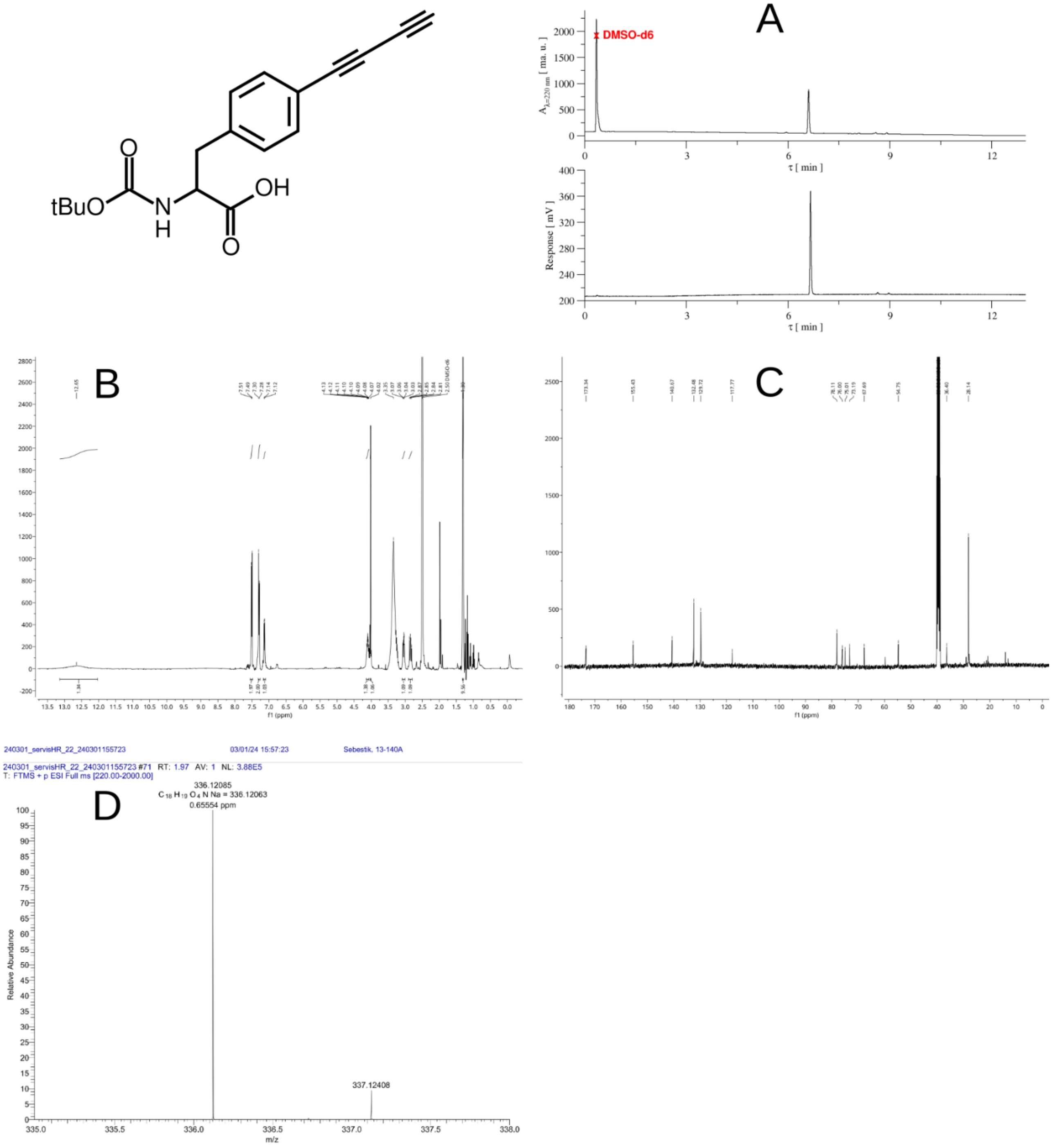

**Compound characterization graph S3.** Formula of compound **3**. A) Analytical HPLC records of compound **3**. DAD detector (top) and ELSD (bottom). B) $^{1}$H-NMR compound **3**. C) $^{13}$C-NMR compound **3**. D) HRMS-spectrum of compound **3**.

#### L-3-(4- (buta-1,3-diyn-1-yl)phenyl-2-((*tert*-butoxycarbonyl)amino)propanoic acid (3) – method B (Compound characterization graph S4)

To a stirred solution of Boc-Phe($C_4$TMS)-OH (**2**, 1.28 g, 3.32 mmol) in THF (20 mL), $Bu_4NF\cdot 3H_2O$ (1.59 g, 5.04 mmol) in $H_2O$ (530 µL) was added portion wise (10 µL additions). The reaction mixture was slightly bubbling (released TMS-F b.p. 16 °C). The addition was finished within 24 min. After 58 min, it was evaporated to dryness, redissolved in EtOAc (50 mL)/1M $KHSO_4$ (30 mL) and washed with 1M $KHSO_4$ (2×30 mL). After drying with $Na_2SO_4$ and filtration, EtOAc was evaporated and residuum dried 1.005 g (3.21 mmol, 97%). **HPLC**: $R_T$ 6.6 min. According to HPLC, the product prepared by method B was identical with those prepared by method A.

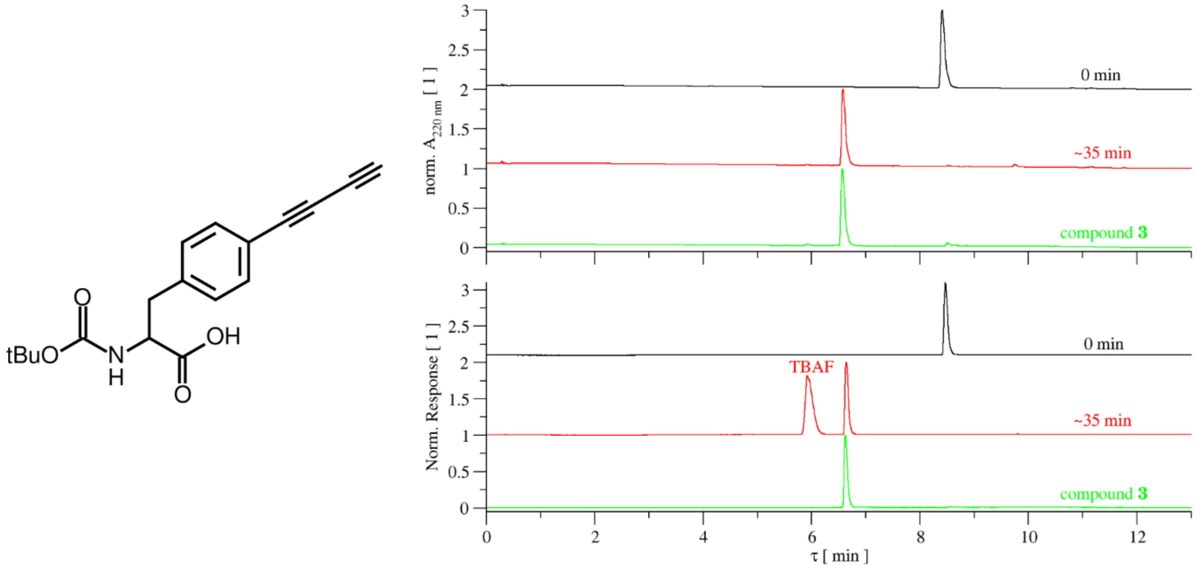

**Compound characterization graph S4.** Formula of compound **3**. Analytical HPLC records of TMS deprotection reaction and of obtained compound **3**. DAD detector (top) and ELSD (bottom).

#### L-3-(4- (4-(diethylamino)-2-(diethylammonio)buta-3-en-1-yl)phenyl-2-((*tert*-butoxycarbonyl)amino)propanoat (4)

It was obtained as a by-product of column chromatography during synthesis of the compound **3** by method B. After subsequent preparative TLC separation, during which original compound **3** completely vanished. TLC $R_F$(EtOAc/DEA/MeOH 75:25:32) 0.33. Yield **4** ~7%. **HPLC**: $R_T$ 5.7 min. **ESI-HRMS** (m/z): for $[M]^-$ $C_{26}H_{40}O_4N_3$ calcd. 458.30243; found 458.30237 (-0.12 ppm). **$^1$H-NMR** (400 MHz, DMSO-*d6*) δ 8.13 (d, J = 12.1 Hz, 1H, =C**H**-,$Bu_4$), 7.13 (d, J = 7.8 Hz, 2H, **H**2,$_{Ph}$), 7.00 (d, J = 7.8 Hz, 2H, **H**3,$_{Ph}$), 5.99 (d, J = 6.5 Hz, 1H, CON**H**), 5.46 (d, J = 12.2 Hz, 1H, =C**H**-,$Bu_3$), 4.11 (s, 2H, C**H**$_2$,$Bu_1$), 3.75 (q, J = 5.9 Hz, 1H, **H**α), 3.63 (q, J = 7.2 Hz, 2H, C**H**$_2$, $N^+Et_2$), 3.54 (dt, J = 14.3, 7.2 Hz, 4H, C**H**$_2$, $NEt_2$), 3.41 (q, J = 7.1 Hz, 2H, C**H**$_2$, $N^+Et_2$), 3.02 (dd, J = 13.3, 5.0 Hz, 1H, **H**β), 2.88 (dd, J = 13.2, 6.1 Hz, 1H, **H**β), 1.32 (s, 9H, C**H**$_{3,Boc}$), 1.20 (t, J = 7.0, 3H, C**H**$_3$, $N^+Et_2$), 1.18 (td, J = 7.3, 2.3 Hz, 6H, C**H**$_3$, $NEt_2$), 0.98 (t, J = 7.0 Hz, 3H, C**H**$_3$, $N^+Et_2$). **$^{13}$C-NMR** (100 MHz, DMSO-*d6*) δ 172.38 (**C**$OO^-$), 168.14 (**C**=$N^+$,$Bu_2$), 158.38 (=**C**H-N), 154.58 (**C**ONH), 138.01 (**C**1,$_{Ph}$), 133.33 (**C**4,$_{Ph}$), 130.14 (**C**2,$_{Ph}$), 126.77 (**C**3,$_{Ph}$), 90.67 (=**C**H-C=$N^+$), 77.16 (**C**$_{Boc}$), 55.88 (**C**α), 51.20 (**C**$H_2$, $NEt_2$), 46.55 (**C**$H_2$, $N^+Et_2$), 44.93 (**C**$H_2$, $N^+Et_2$), 42.84 (**C**$H_2$, $NEt_2$), 36.68 (**C**β), 32.53 (**C**$H_2$,$Bu_1$), 28.22 (**C**$H_{3,Boc}$), 14.48 (**C**$H_3$, $NEt_2$), 13.24 (**C**$H_3$, $N^+Et_2$), 11.86 (**C**$H_3$, $NEt_2$), 11.56 (**C**$H_3$, $N^+Et_2$). According to NMR data, compound was contaminated with DEA.

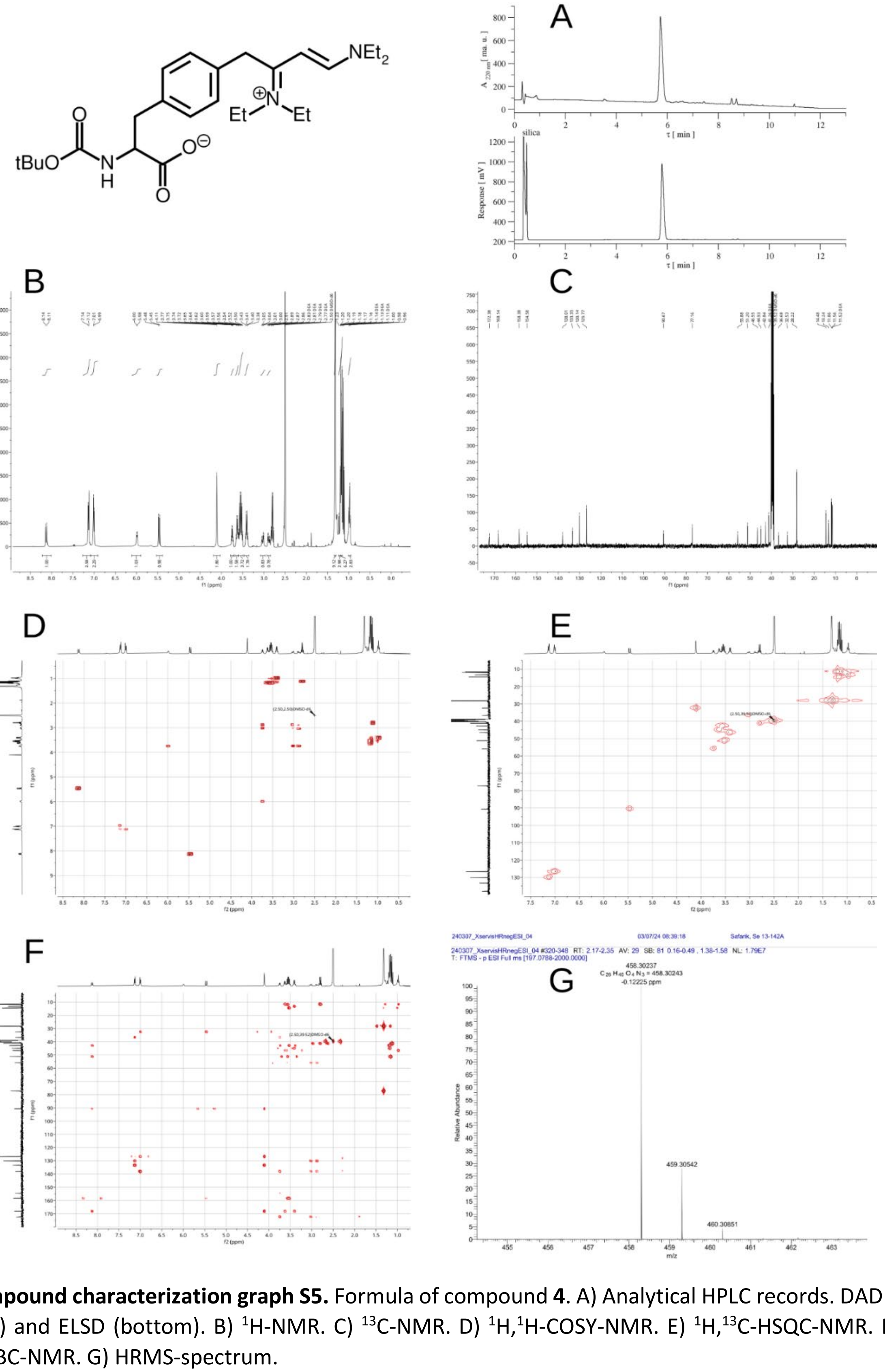

**Compound characterization graph S5.** Formula of compound **4**. A) Analytical HPLC records. DAD detector (top) and ELSD (bottom). B) $^{1}H$-NMR. C) $^{13}C$-NMR. D) $^{1}H$,$^{1}H$-COSY-NMR. E) $^{1}H$,$^{13}C$-HSQC-NMR. F) $^{1}H$,$^{13}C$-HMBC-NMR. G) HRMS-spectrum.

#### L-3-(4-(buta-1,3-diyn-1-yl)phenyl-1-carboxyethan-1-aminium trifluoroacetate (5) (Compound characterization graph S6)

Boc-Phe(C4H)-OH (1.005 g, 3), DCM (19 mL), and anisole (8 mL) was cooled down to 0°C, then TFA (25 mL) was added. The intensively green solution was formed. After 46 min, the excess of solvents was blown out by stream of N2 maintaining the cooling bath at 0°C. After 4 h, the remaining anisole was evaporated at 3 mbar/19°C (low temperature of bath was maintained by addition of ice) and dried for additional 2h Boc-Phe($C_4$H)-OH (1.005 g, **3**), DCM (19 mL), and anisole (8 mL) was cooled down to 0°C, then TFA (25 mL) was added. The intensively green solution was formed. After 46 min, the excess of solvents was blown out by stream of $N_2$ maintaining the cooling bath at 0°C. After 4 h, the remaining anisole was evaporated at 3 mbar/19°C (low temperature of bath was maintained by addition of ice) and dried for additional 2h. The weight of crude residuum was 1.509 g, and it contained some anisole according to HPLC analysis. The residuum was dissolved in 10% ACN and freeze dried over weekend. A black-brown powder (1.023 g, ~97%) was obtained. Freeze drying led to removal of anisole; however, the formation of tars impurity occurred (detectable by ELSD in HPLC). The quality of sample is sufficient for preparation of recombinant proteins with incorporated non-coding amino acids (see also section Stability of prepared butadiynes). Extra pure DAF (**5**) was obtained by semipreparative HPLC on the VYDAC 250 × 10 mm, 10 µm RP-18 column with a flow rate 3 mL/min using a 0–100% ACN (acetonitrile) gradient in 0.05% aqueous TFA. **HPLC**: $R_T$ 3.84 min. **ESI-HRMS** (m/z): for $[M+Na]^+$ $C_{13}H_{11}O_2NNa$ calcd. 236.06820; found 236.06843 (0.99 ppm); $[M+H]^+$ $C_{13}H_{12}O_2N$ calcd. 214.08626; found 214.08639 (0.65 ppm). **$^1$H-NMR** (400 MHz, DMSO-*d6*) δ 8.32 (s, 2H, N**H**$_2$), 7.56 (d, J = 8.4 Hz, 2H, **H**3,$_{Ph}$), 7.31 (d, J = 8.3 Hz, 2H, **H**2,$_{Ph}$), 4.18 (t, J = 6.5 Hz, 1H, **H**α), 4.04 (s, 1H, ≡C-**H**), 3.33 – 2.94 (m, 2H, **H**β). **$^{13}$C-NMR** (100 MHz, DMSO-*d6*) δ 170.29 (**C**OOH), 158.24 (q, J = 31.3 Hz, $CF_3$**C**OOH), 137.54 (**C**1,$_{Ph}$), 132.89 (**C**3,$_{Ph}$), 130.11 (**C**2,$_{Ph}$), 118.73 (**C**4,$_{Ph}$), 117.21 (q, J = 299 Hz, **C**$F_3$), 76.27 (≡**C**-H), 74.77 (≡**C**-Ph), 73.54 (≡**C**-), 67.64 (≡**C**-), 53.00 (**C**α), 35.81 (**C**β). $^{13}$C-NMR presence of trifluoroacetate corresponds well with our previous data for inubosin trifluoroacetate[2] i.e. two quartets were observed around 158 and 117 ppm, first with $J_{C-F}$ ~ 30 Hz and second with $J_{C-F}$ ~ 300 Hz. **$^{19}$F-NMR** (376 MHz, DMSO-*d6*) δ -76.55. **IR** (film on $CaF_2$, $cm^{-1}$): ν($N^+$H) 3337, ν(C≡C-C≡C)$_{symm}$ 2203, ν(C≡C-C≡C)$_{as}$ 2129, ν(C=O) 1747sh, ν(C=O) 1678.

**Compound characterization graph S6 (next page).**Formula of compound **5**. A) Analytical HPLC records of combined fractions from HPLC preparation of compound **5**. DAD detector (top) and ELSD (bottom). #E was rich on tar impurities. B) $^1$H-NMR compound **5**. C) $^{13}$C-NMR compound **5**. D) $^{19}$F-NMR compound **5**. E) $^1$H,$^1$H-COSY-NMR compound **5**. F) $^1$H,$^{13}$C-HSQC-NMR compound **5**. G) $^1$H,$^{13}$C-HMBC-NMR compound **5**. H) HRMS-spectrum. I) HRMS-spectrum. J) IR spectrum of compound **5**, film on $CaF_2$.

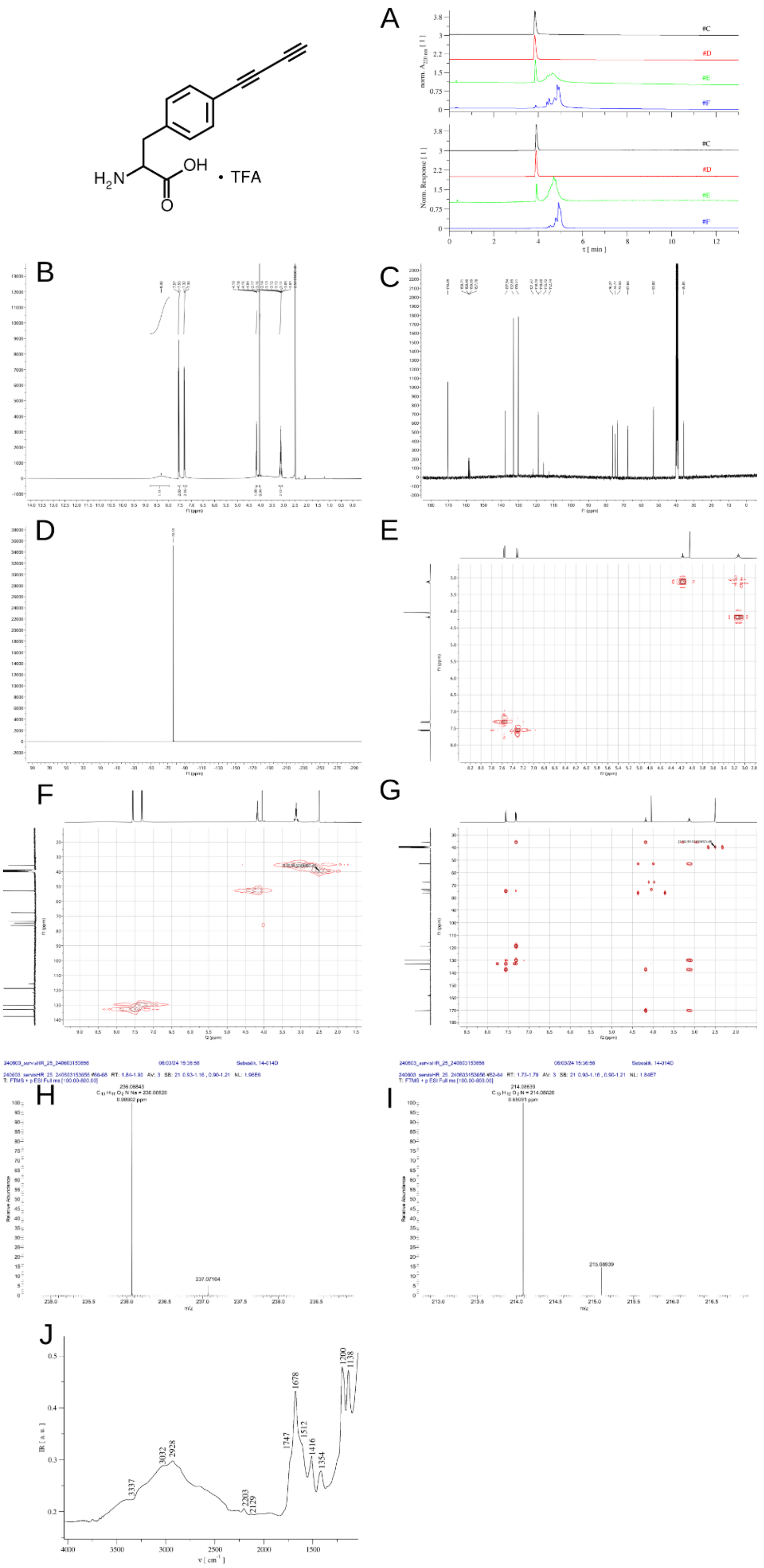
H2N
OH
O
• TFA
A
norm. A220 nm [ 1 ]
#C
#D
#E
#F
Norm. Response [ 1 ]
τ [ min ]
B
C
D
E
F
G
H
I
J
IR [ a. u. ]
ν [ cm⁻¹ ]
3337
3032
2928
2203
2129
1747
1678
1512
1416
1354
1200
1138

#### Stability of prepared butadiynes.

Originally these terminal dialkyne compounds should be colorless; however, short standing at room temperature lead to formation of dark brown impurities, which presence prevented measurement of slow Raman spectra due to strong fluorescence. For starting material **1**, it was already reported in literature,[1] where freshly distilled colorless product turned yellow or brown upon standing. The compounds are very sensitive to heating, thus, during preparations when evaporation is necessary, it is advisable to add ice to cool down evaporator bath. As shown in figure below, two months standing at room temperature led to almost complete decomposition of compound **5**. Most of the sample polymerized (signal at exclusion limit of the column in inject peak) i.e. ca ~3% DAF remained in the sample. Noteworthy, the main impurities obtained in batches of crude sample are those tars that were also formed by long time decomposition. Heating of samples led to higher abundance of these impurities. Therefore, we have tested compatibility of tars with living cells, and crude samples were suitable for protein expression. Due to thermal instability of DAF, any handling at room temperature led to worsening of sample quality, for instance, ionex chromatography was used for anion exchange $CF_3COO^-$ to $Cl^-$. It also led to higher abundance of various tar impurities and polymers.

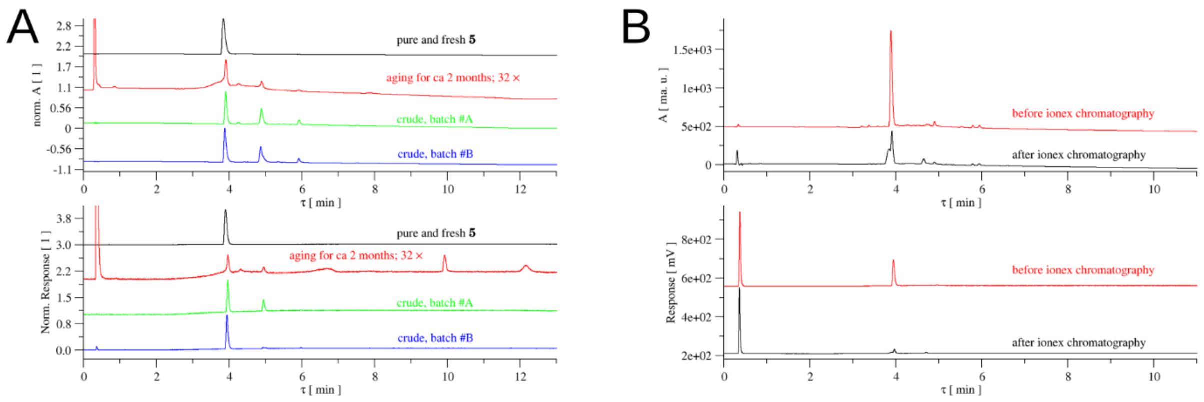

**Compound characterization graph S7.** A) Analytical HPLC records showing aging of DAF (**5**, cf. black and red records). Data for two batches of crude compound **5** are also shown (from the blue one, the pure compound was obtained). DAD detector (top) and ELSD (bottom). B) Analytical HPLC records showing changes of DAF quality by applying ionex chromatography TFA-HCl exchange. DAD detector (top) and ELSD (bottom).

#### Note S2. Chemical reactivity of DAF.

DAF and ETF reactivity were determined by mass spectrometry and fluorescence spectroscopy. In the case of probing reactivity with mass spectrometry, we look for the masses of adducts between DAF/ETF and the other compounds according to **Tables S1A-C** (**Fig. S2**).

As expected, we observed the masses corresponding to DAF-azide (**Fig. S2A**), DAF-thiol (**Fig. S2B**) adducts. In the former case, only in the presence of copper. Two DAF-tetrazine adducts were observed, one corresponding to the aggregate mass of the reacting compounds and another one with higher abundance having 28 Da less (**Fig. S2C**), which may be attributed to the release of nitrogen gas. Also, the adduct between DAF and TCEP was found (**Fig. S2F**). Apart from cysteine and selenocysteine, no reactivity was observed between DAF and canonical amino acids, though in case of selenocysteine, the masses corresponding to neither reactants nor the adduct were observed and formation of an intermediate was assumed (**Fig. S2D** and **Fig. S2F**). Between the two tripeptides: glutathione reduced and glutathione oxidized, DAF showed reactivity towards the former one through the adduct formation and but not towards the later. Interestingly, DAF evidenced dimer formation at higher concentration and formed adducts with other two non-canonical amino acids: ETF and Tet.2-Et, among the four non-canonical amino acids tested (**Fig. S2E**). The observed reactivity is not exclusive of DAF. The related non-canonical amino acid 4-ethynyl-phenylalanine (ETF, **Fig. 1**), which contains a single alkyne, can also react with azide, thiol, and H-tetrazine (with the mass corresponding to nitrogen gas release only) (**Fig. S2E**), albeit the product yields are somewhat lower compared to DAF, suggesting slower kinetic rates.

| Reactant with DAF | Expected mass (Dalton) [M+H]$^+$ | Observed mass (Dalton) [M+H]$^+$ | Reacted (adduct) | Did not react | Reacted (unknown) |
|---|---|---|---|---|---|
| **DTT** | 368.098475 | 368.0982 | ✓ | | |
| **BME** | 292.10019 | 292.1006 | ✓ | | |
| **H-Tetrazine Amine** | 401.17205[a] | 401.17205 | ✓ | | |
| | 373.165902[b] | 373.16586 | | | |
| **Methyl-Tetrazine Amine** | 415.1877[a] | 415.18763 | ✓ | | |
| | 387.181552[b] | 387.18149 | | | |
| **Benzyl Azide** | 347.150252 | 347.15554 | ✓ | | |
| **Cysteine** | 335.106003 | 335.1062 | ✓ | | |
| **Glutathione reduced** | 521.17006 | 521.1703 | ✓ | | |
| **TCEP** | 464.146879 | 464.1471 | ✓ | | |
| **Glutathione oxidized** | 826.238215 | | | ✓ | |
| **Selenocysteine** | 383.050455 | | | | ✓ |
| **Aspartic acid** | 347.123763 | | | ✓ | |
| **Glutamic acid** | 361.139413 | | | ✓ | |
| **Histidine** | 367.140082 | | | ✓ | |
| **Methionine** | 363.137304 | | | ✓ | |
| **Serine** | 319.128848 | | | ✓ | |
| **Tyrosine** | 395.160148 | | | ✓ | |
| **DAF** | 427.165234 | 427.16537 | ✓ | | |
| **ETF** | 403.165234 | 403.16532* | ✓ | | |
| **Tet.2-Et** | 487.20883 | 487.20906* | ✓ | | |
| **BCNK** | 536.275512 | | | ✓ | |
| **SCOK** | 510.259862 | | | ✓ | |

[a] Without $N_2$ release, [b] With $N_2$ release
*Significantly low abundance
**Table S1.A.**

| Reactant with DAF | Possible adducts between DAF and tetrazine amine | Expected mass of Product (Dalton) $[M+H]^+$ | Observed mass of Product (Dalton) |
|---|---|---|---|
| **H-Tetrazine Amine** | 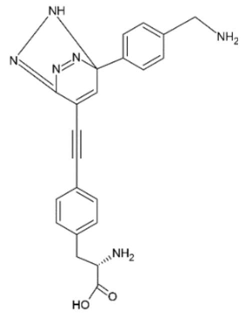 | 401.17205 | 401.17205 |
| | 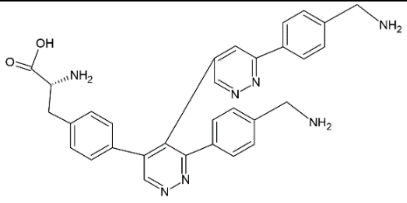 | 532.24555 | No |
| | 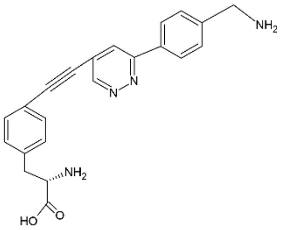 | 373.165902 | 373.16586 |
| | 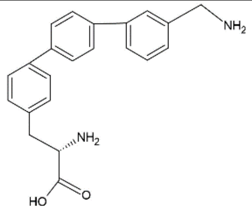 | 347.175404 | No |

| | | | |
|---|---|---|---|
| **Methyl-Tetrazine Amine** | 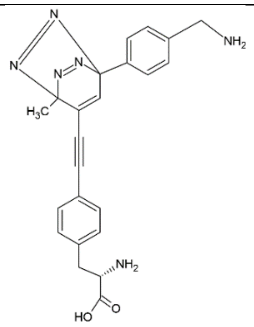 | 415.1877 | 415.18763 |
| | 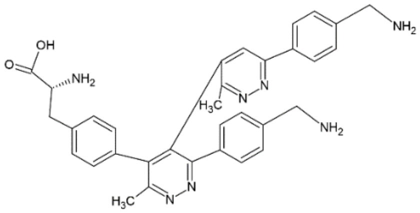 | 560.27685 | No |
| | 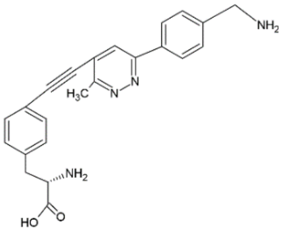 | 387.181552 | 387.18149 |
| | 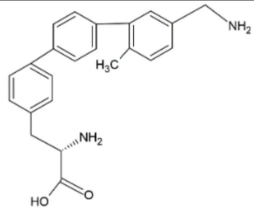 | 361.191054 | No |

**Table S1.B**

| Reactant with ETF | Expected mass of Product (Dalton) | Observed mass of Product (Dalton) | Reacted and formed adduct with DAF | Did not react with DAF |
|---|---|---|---|---|
| **DTT** | 343.098475 | 343.1654 | ✓ | |
| **BME** | 268.10019 | 268.10407 | ✓ | |
| **H-Tetrazine** | 377.17205[a] | - | | |
| | 349.165902[b] | 349.16599 | ✓ | |
| **Methyl-Tetrazine** | 391.1877[a] | - | | ✓ |
| | 363.181552[b] | - | | |
| **Benzyl Azide** | 323.150252 | 323.15501 | ✓ | |

[a] Without $N_2$ release, [b] With $N_2$ release

**Table S1.C.**

### (A) DAF + AZIDES

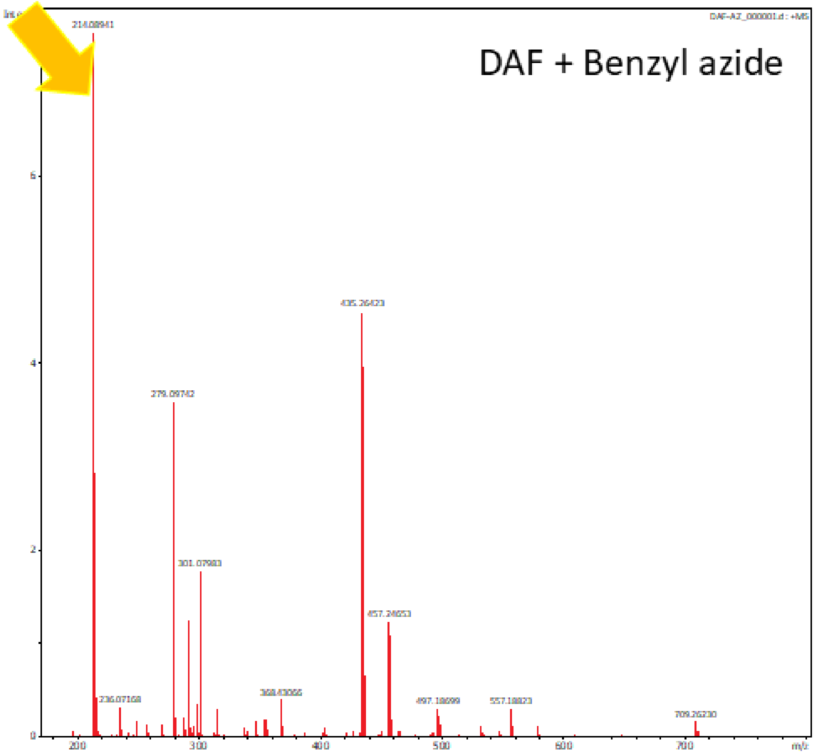

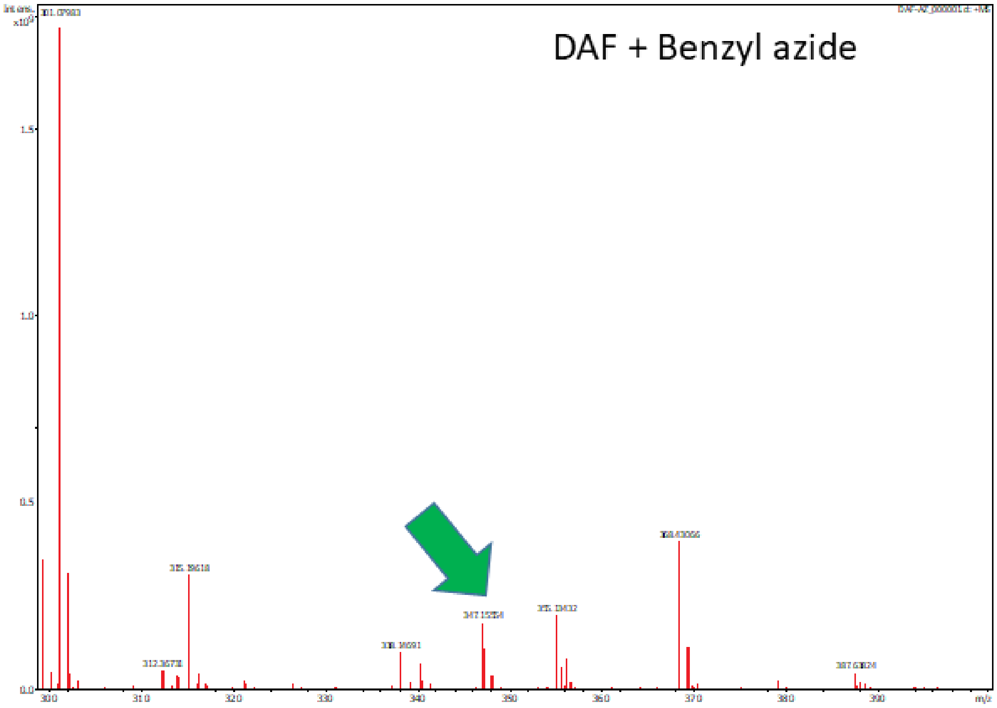

### (B) DAF + THIOLS

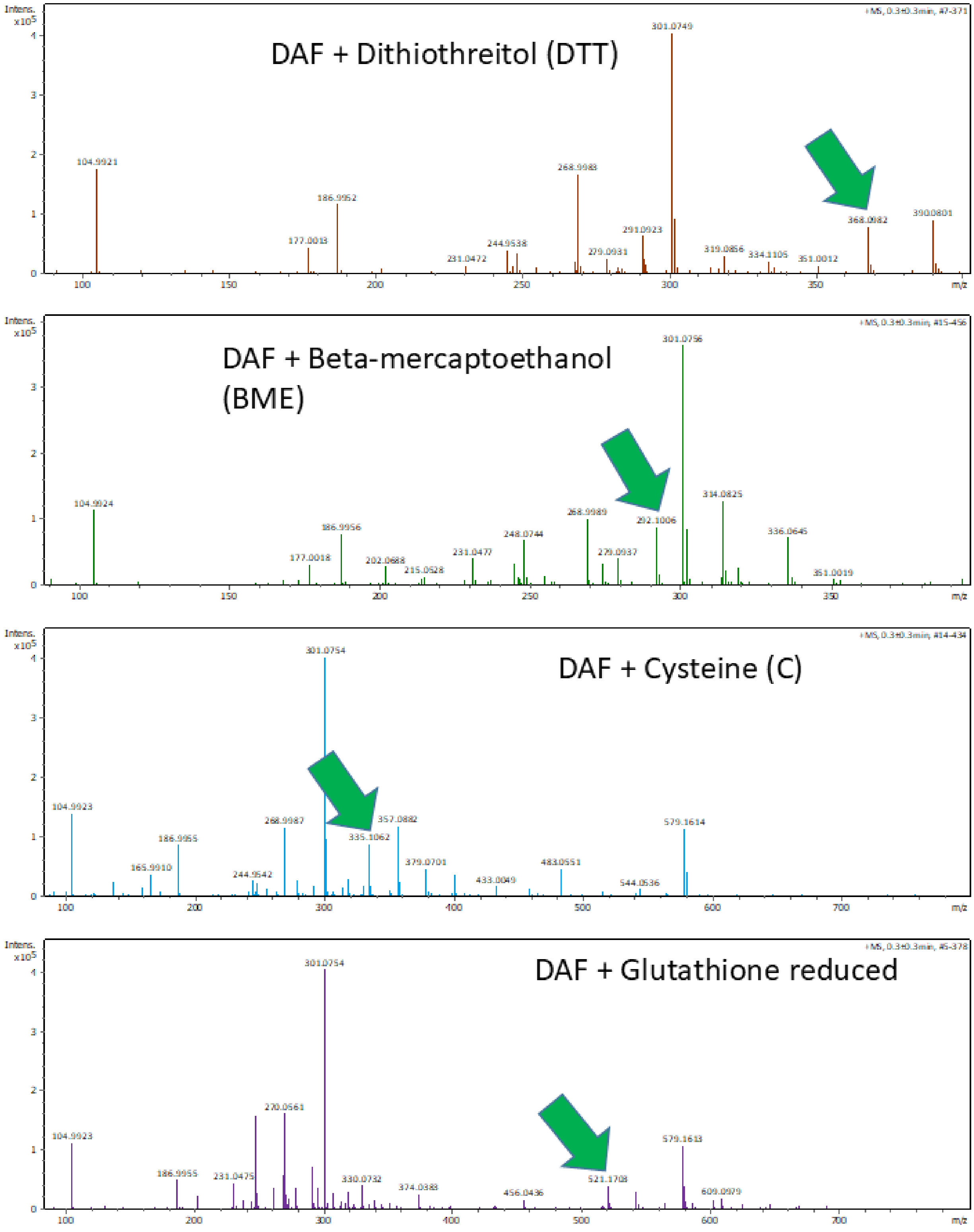
DAF + Dithiothreitol (DTT)
104.9921
177.0013
186.9952
231.0472
244.9538
268.9983
279.0931
291.0923
301.0749
319.0856
334.1105
351.0012
368.0982
390.0801
DAF + Beta-mercaptoethanol (BME)
104.9924
177.0018
186.9956
202.0688
215.0528
231.0477
248.0744
268.9989
279.0937
292.1006
301.0756
314.0825
336.0645
351.0019
DAF + Cysteine (C)
104.9923
165.9910
186.9955
244.9542
268.9987
301.0754
335.1062
357.0882
379.0701
433.0049
483.0551
544.0536
579.1614
DAF + Glutathione reduced
104.9923
186.9955
231.0475
270.0561
301.0754
330.0732
374.0383
456.0436
521.1708
579.1613
609.0979

### (B) DAF + THIOLS (Cont.)

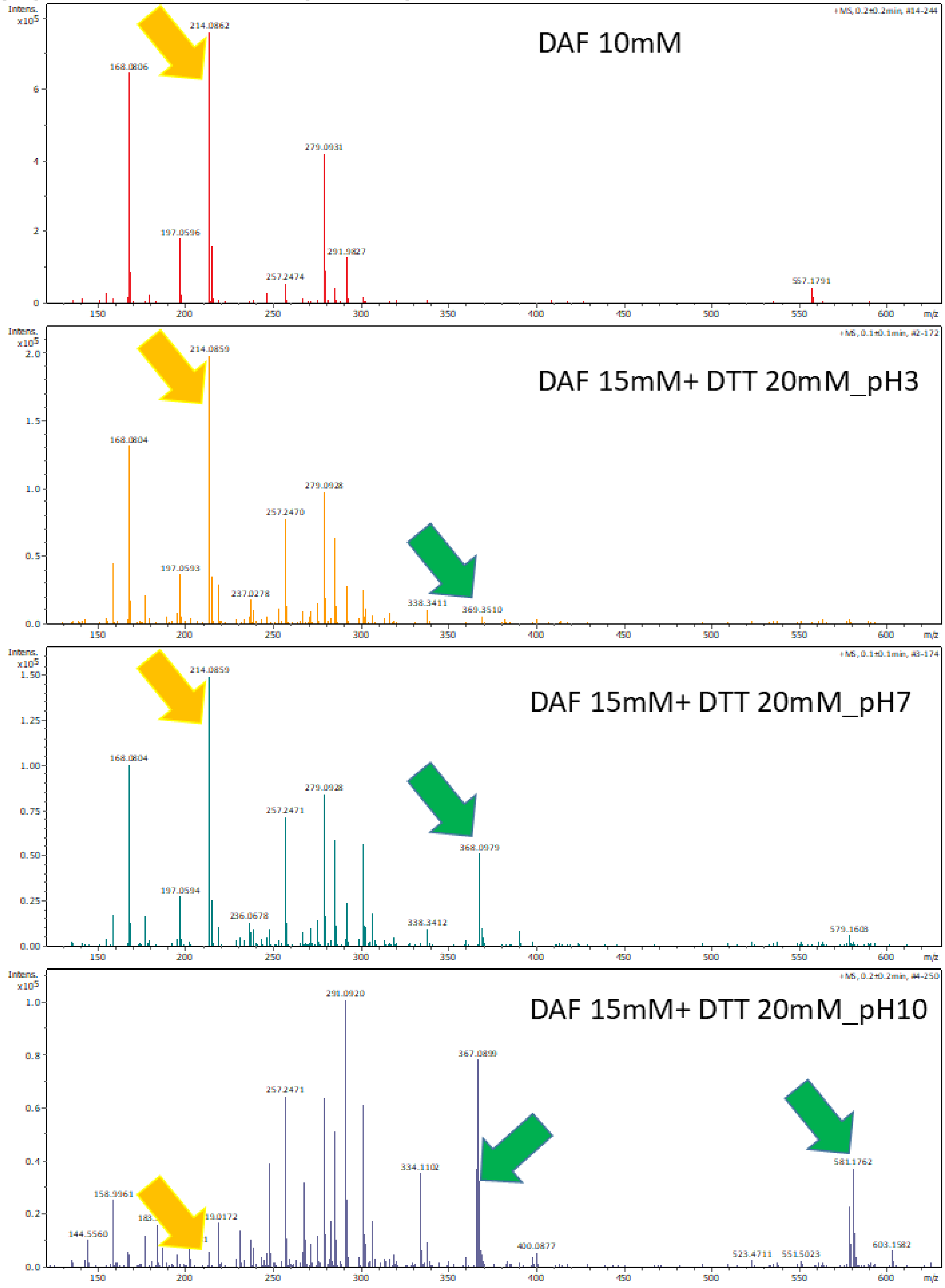

### (C) DAF + TETRAZINES

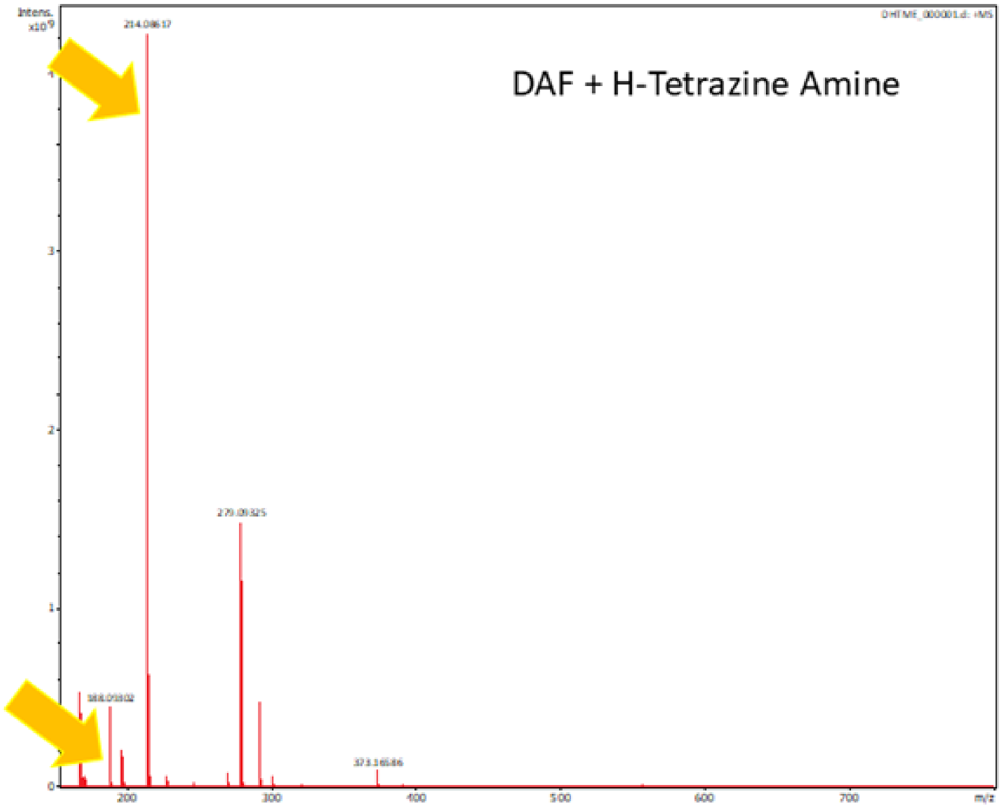

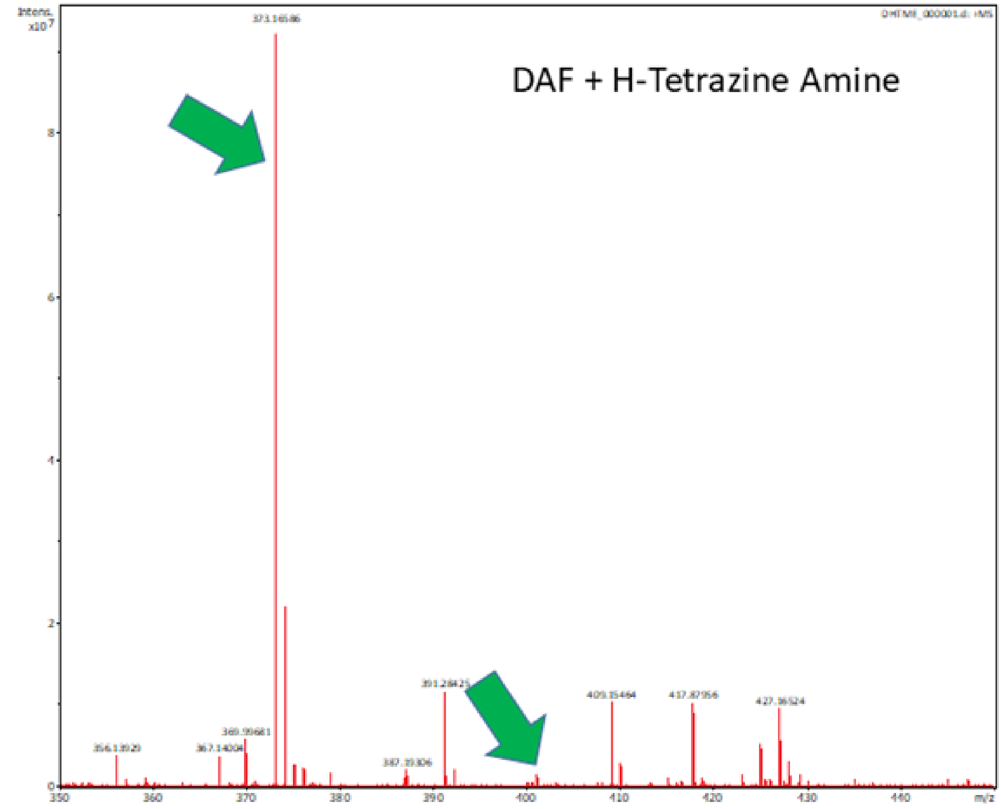

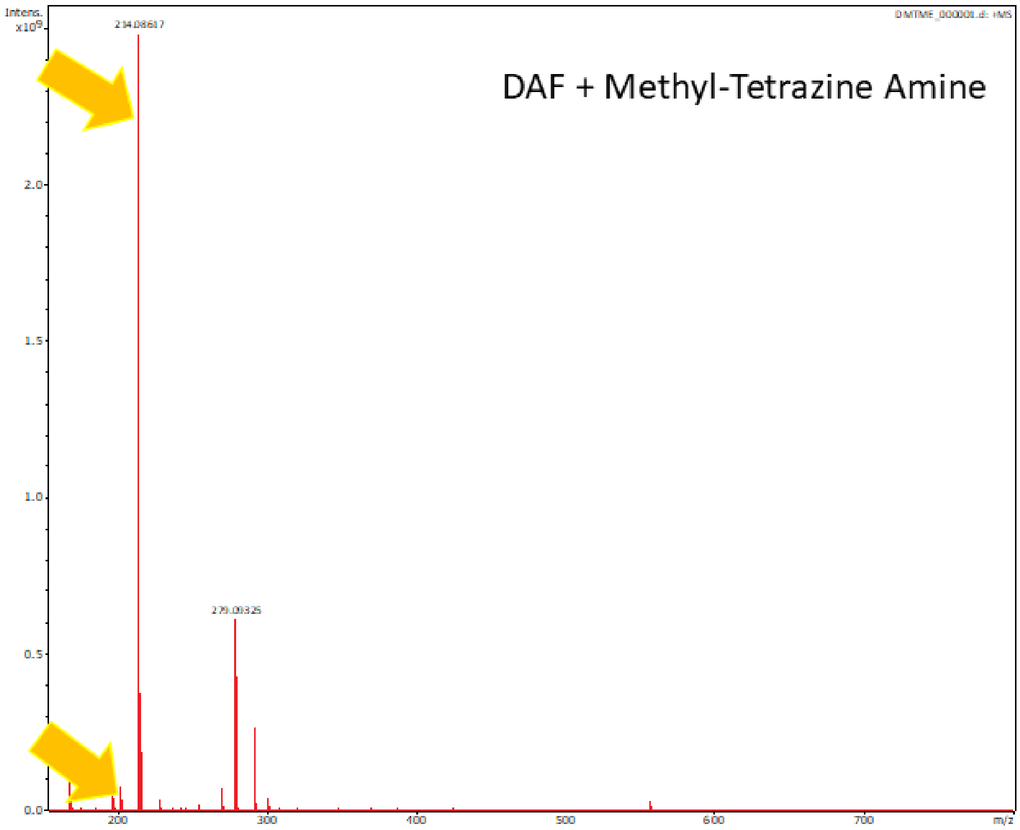
Intens.
x10^9
214.08617
DAF + Methyl-Tetrazine Amine
2.0
1.5
1.0
0.5
0.0
200
300
400
500
600
700
m/z

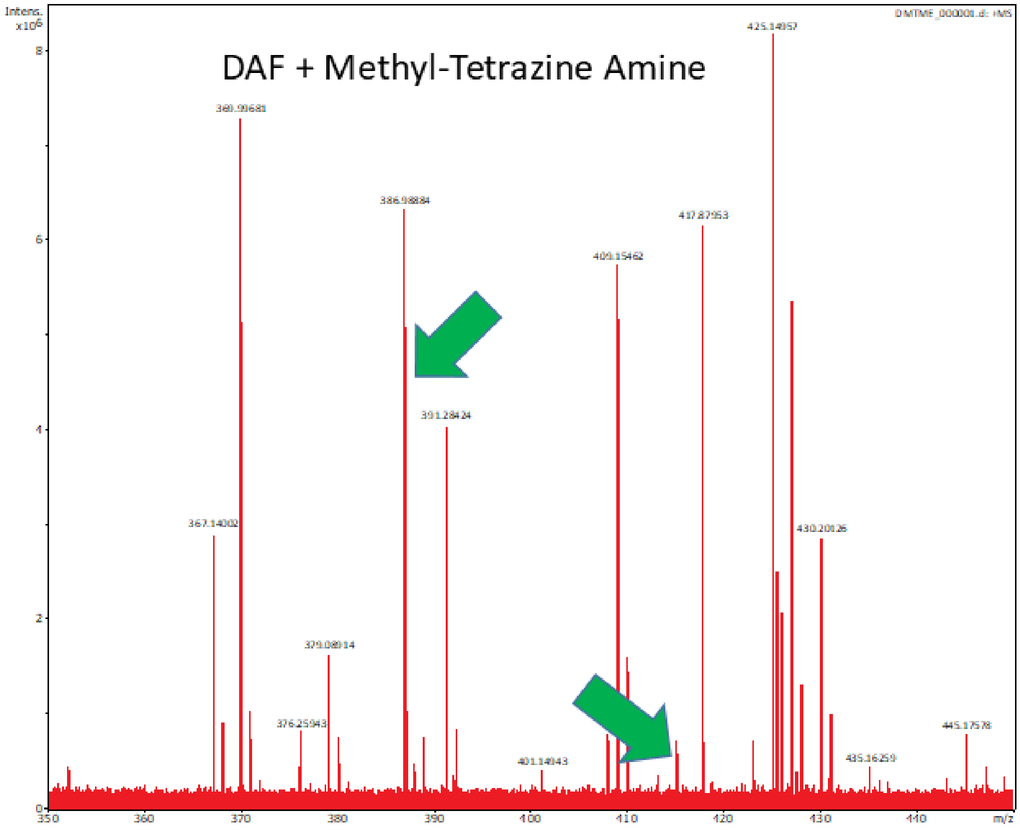
Intens.
x10^6
DAF + Methyl-Tetrazine Amine
8
6
4
2
0
350
360
370
380
390
400
410
420
430
440
m/z

### (D) DAF + AMINO ACIDS

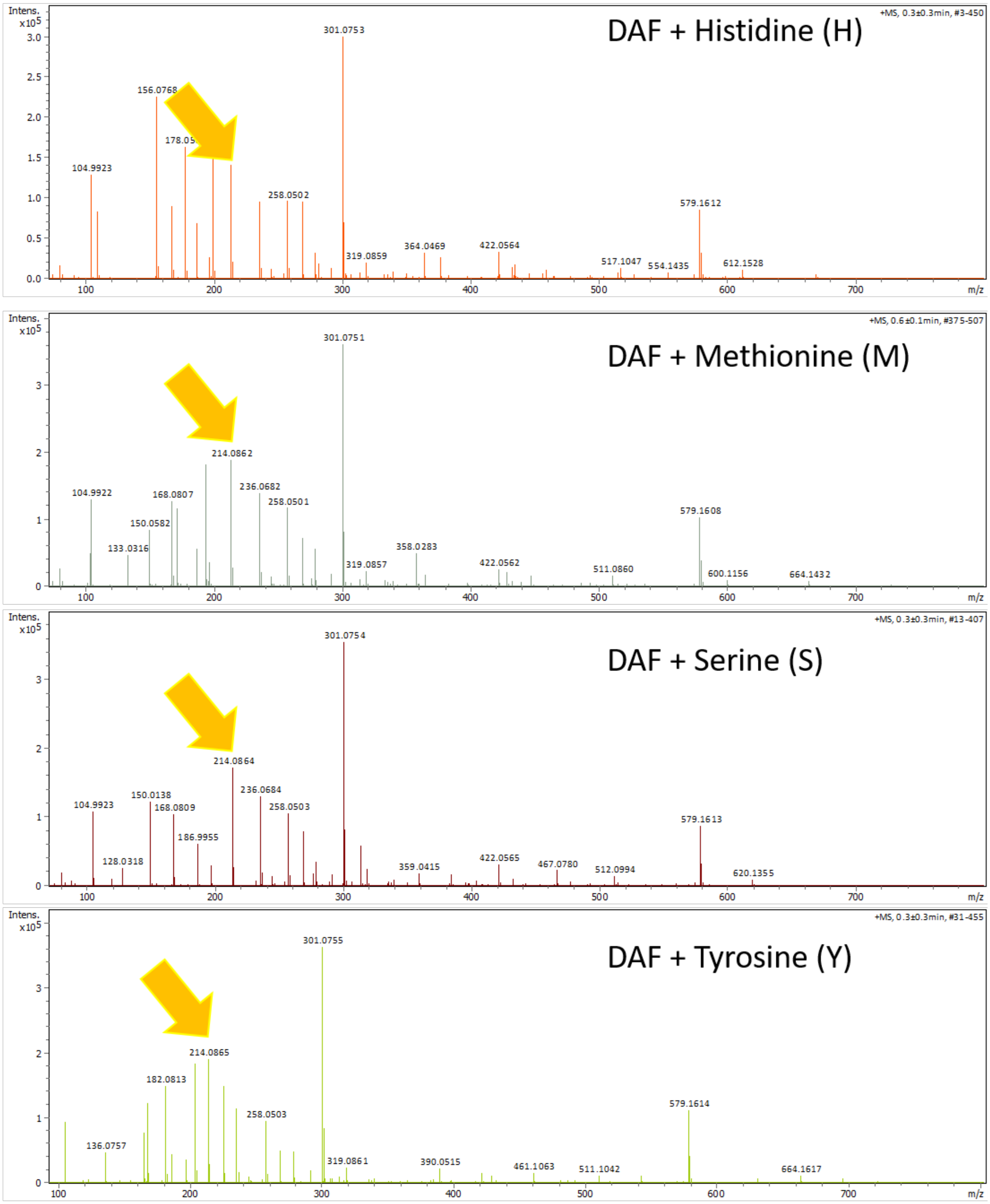

### (D) DAF + AMINO ACIDS (Cont.)

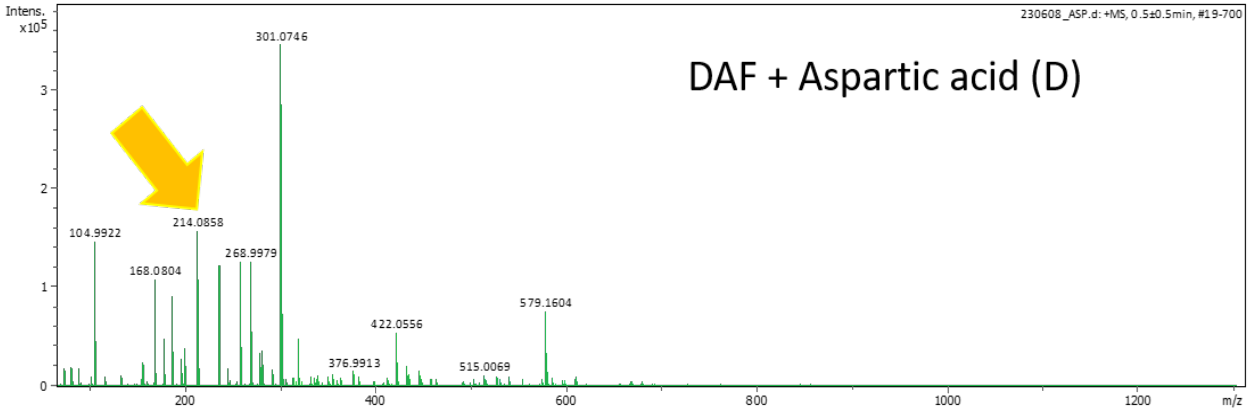

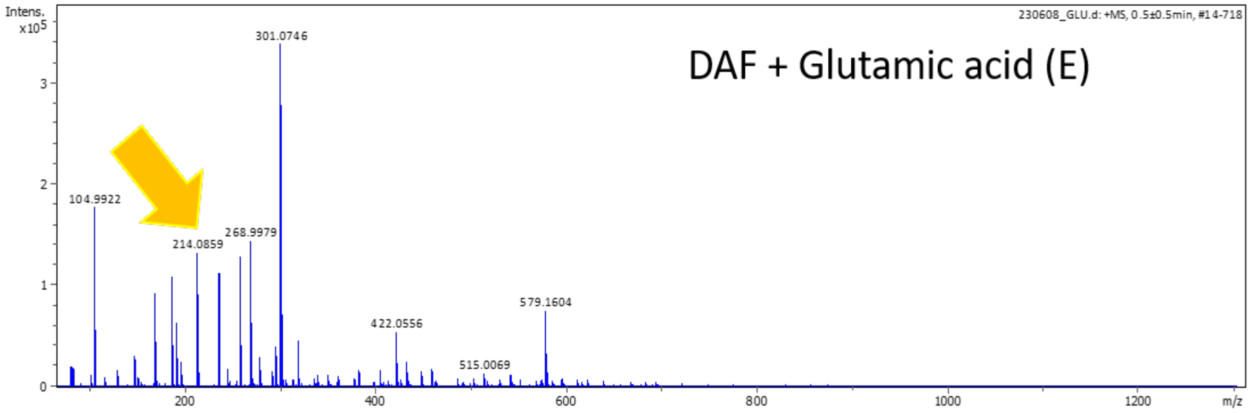

### (E) DAF + NON-CANONICAL AMINO ACIDS

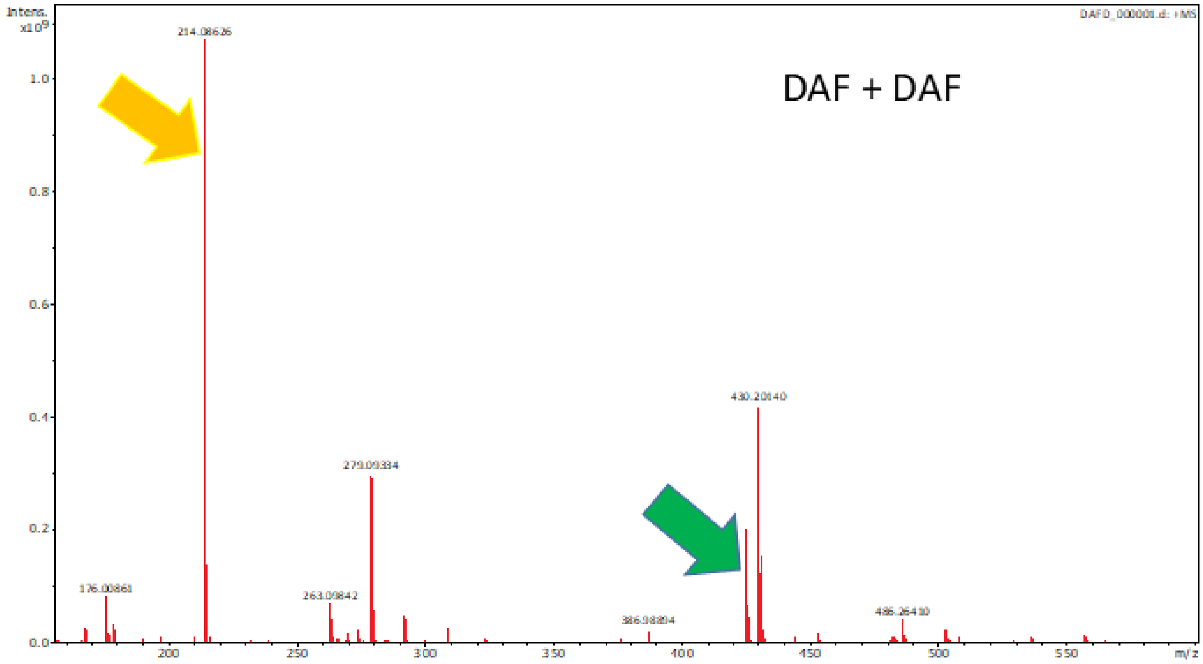

### (E) DAF + NON-CANONICAL AMINO ACIDS (Cont.)

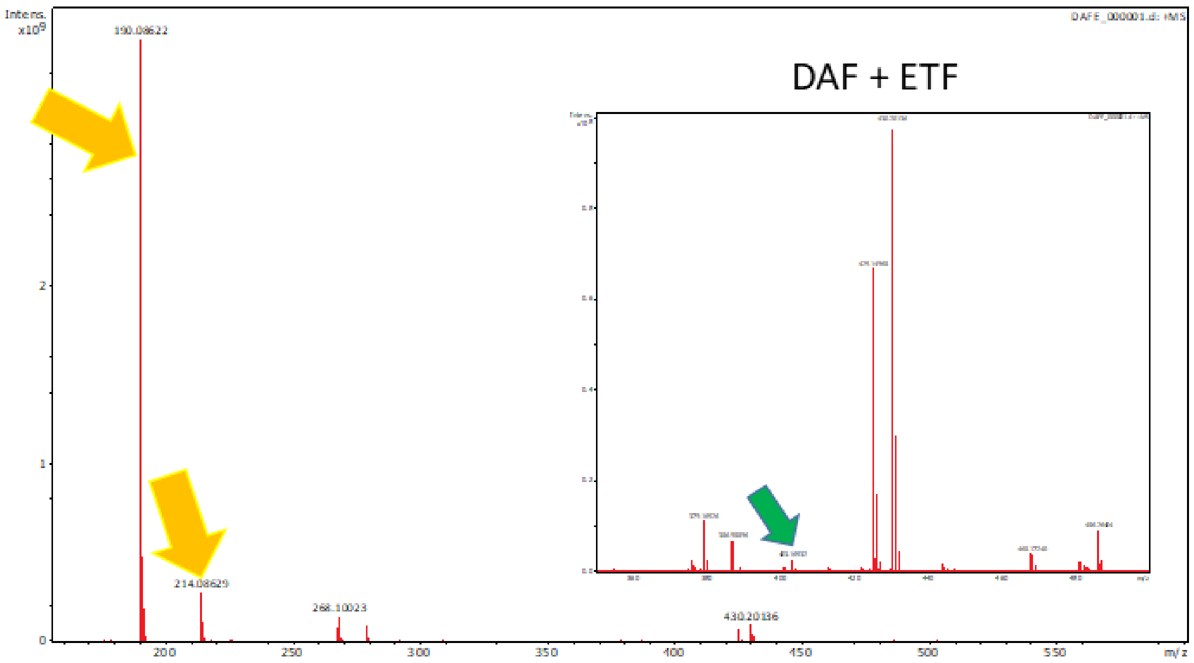

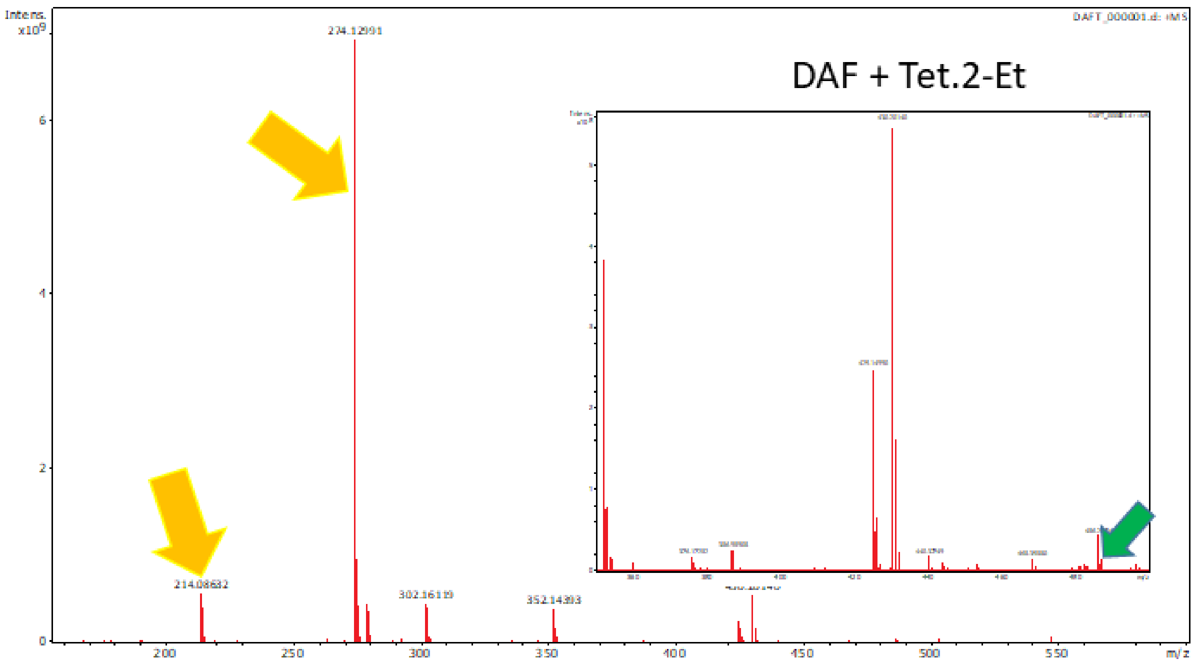

### (E) DAF + NON-CANONICAL AMINO ACIDS (Cont.)

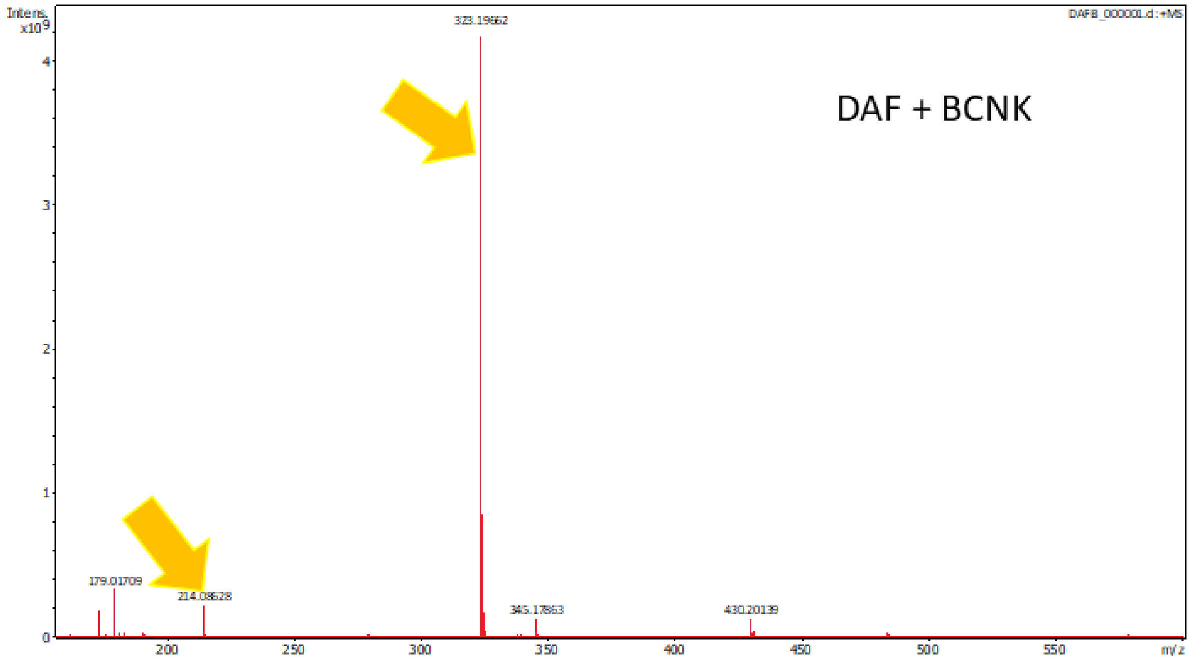

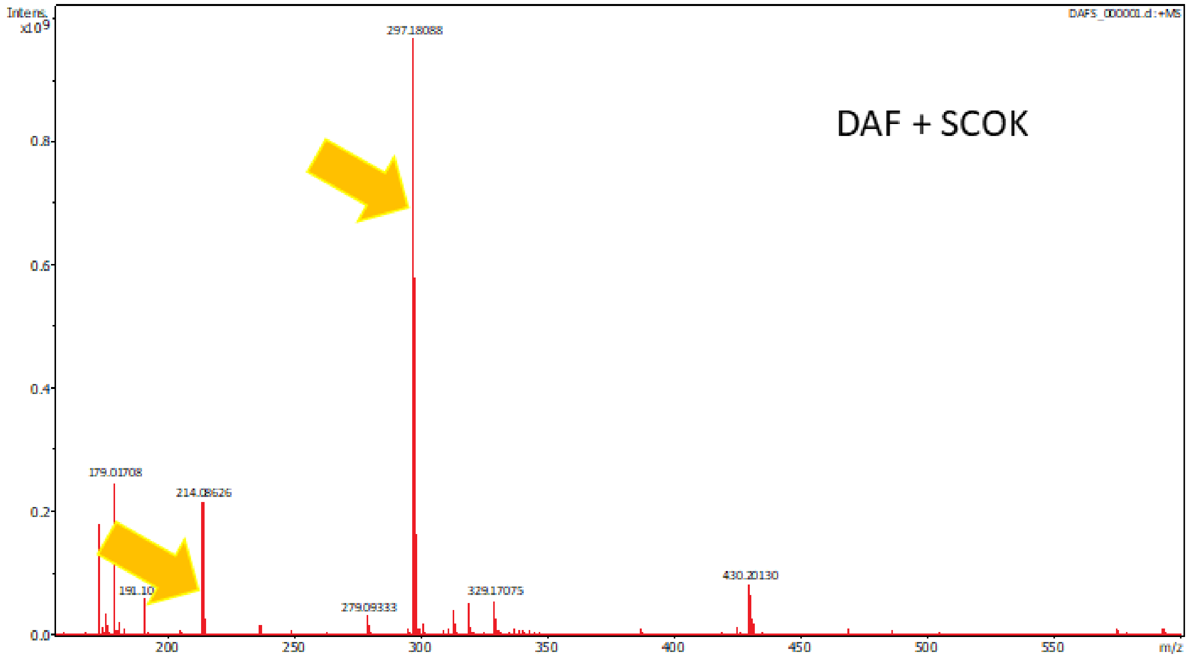

### (F) DAF + TERTIARY PHOSPHINE/DISULFIDE/SELENOL

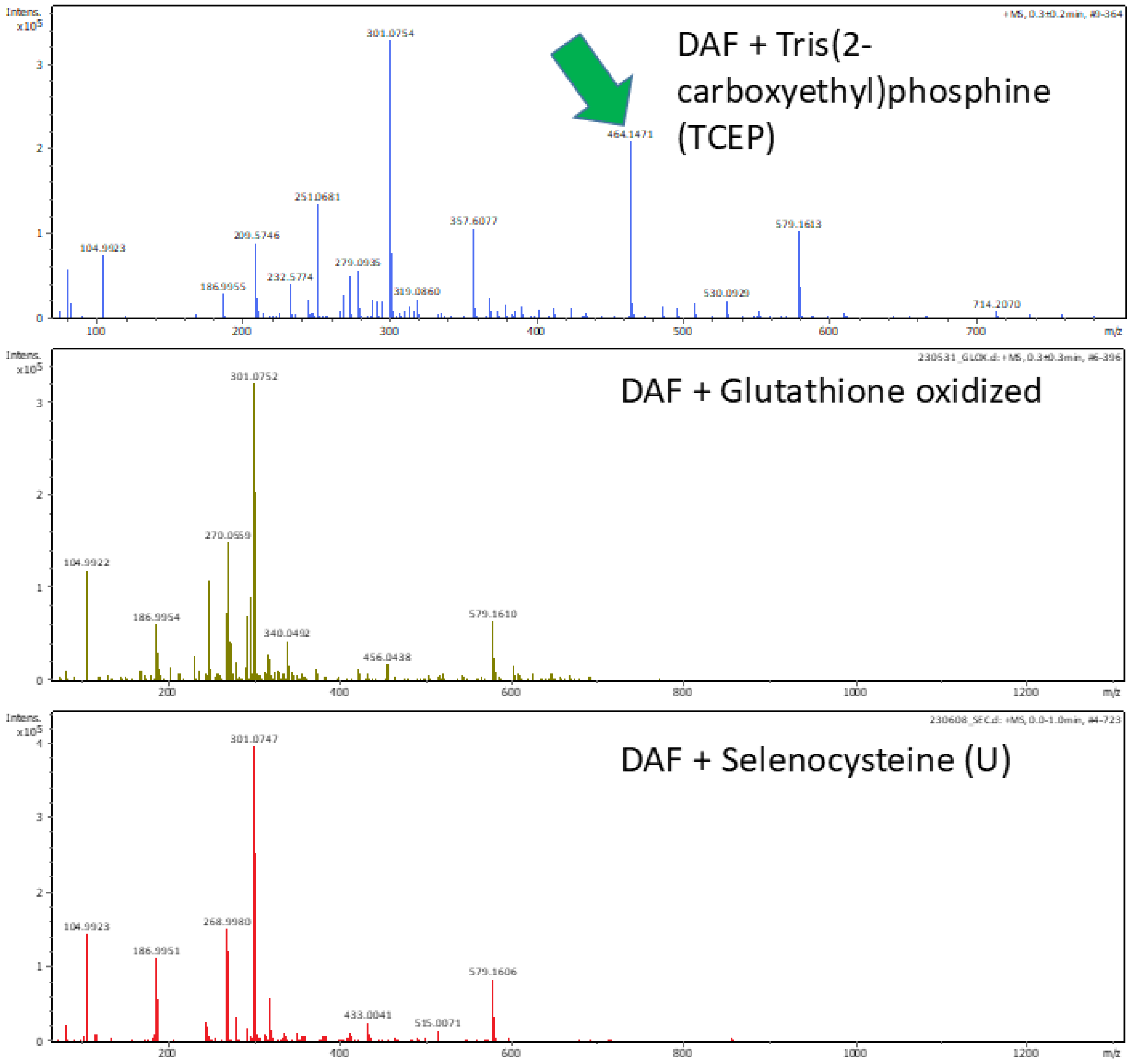

### (G) ETF + AZIDES/THIOLS/TETRAZINES

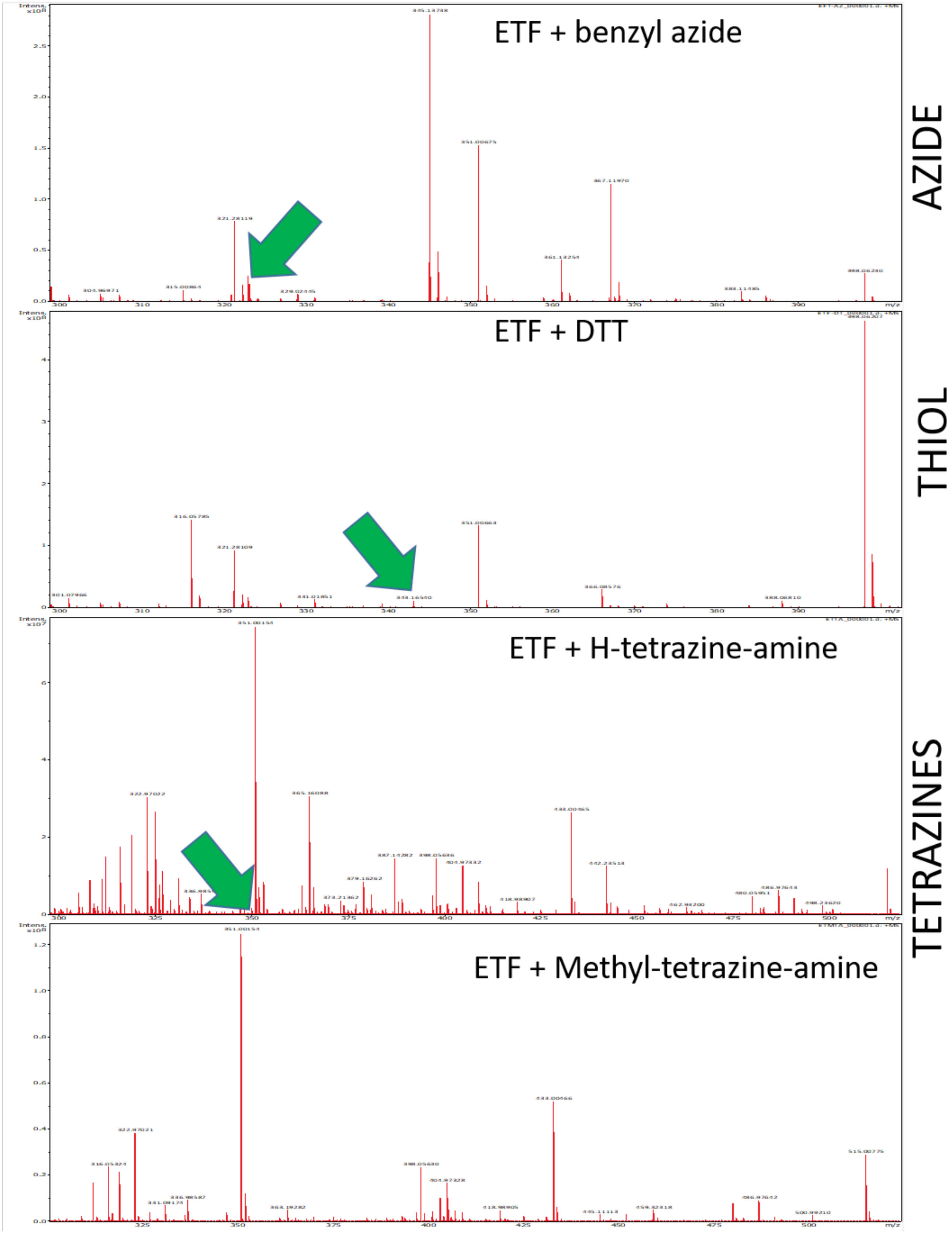

**Fig. S2. Reactivity properties of DAF assessed by mass spectrometry (pages 20-30).** Mass spectra of mixtures of 15 mM DAF with 20 mM of the indicated compounds: azides (A), thiols (B), tetrazines (C), other amino acids (D), other non-canonical amino acids (E) and other reactive moieties (F). In (G), the mass spectra of mixtures of 15 mM ETF with 20 mM of selected compounds are shown. See the method section for the solvents used to dissolve all the reactants. The yellow arrows point to the unreacted reactants, while the green arrows indicate the expected product.

To get semi-quantitative insight into the reaction kinetics, we used fluorogenic probes, which are not fluorescent until reacted. As fluorogenic azide probes, we utilized so called CalFluors, which are activated by copper-catalyzed or metal-free click reaction. In the case of fluorogenic tetrazine probes, we used common tetrazine variants (H-tetrazines), which are also activated upon reaction of the tetrazine (the unreacted tetrazine effectively quenches fluorophore fluorescence).[3] Some tetrazine-functionalized dyes (tetrazine-dyes) function as fluorogenic probes, meaning these compounds substantially increase fluorescence intensity upon reaction with strained dienophiles such as TCO.[4] This makes tetrazine-dyes especially interesting for live-cell labeling and fluorescence imaging applications since the fluorogenic reaction could lower background and potentially eliminate the need for washing out excess fluorophore. In summary, these probes (azide or tetrazine) are poorly fluorescent before they reacted.

The DAF-azide and DAF-tetrazine reactions were also followed by fluorescence spectroscopy. We recorded the "turn on" fluorescence enhancement kinetics upon mixing DAF with these fluorogenic probes (**Fig. S3**). A distinctive feature of DAF kinetics is their multiphasic character, including fast and slow phases. Overall, the reaction rates are relatively slow, typically requiring overnight incubations (16 hours) to reach steady-state levels. When compared to the gold standard transcyclooctene-lysine (TCO*K)[5] ncAA (**Fig. 1**), the tetrazine ligation reaction with DAF is 5-fold slower (compare **Fig. S3C and S3D**), suggesting a bimolecular second-order rate constant of ~1-10 $M^{-1}s^{-1}$. The copper catalyzed alkyne-azide cycloaddition with DAF was compared with that of ETF. DAF showed slower reaction rate than ETF.

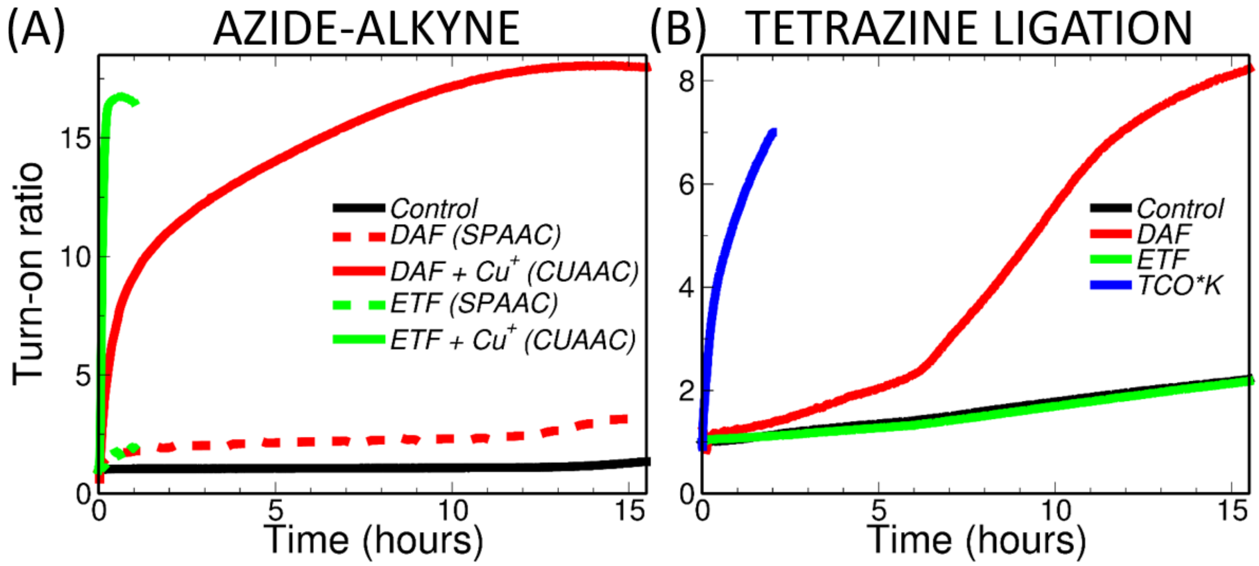

**Fig. S3. Reactivity properties of DAF assessed by fluorescence spectroscopy.** (A) "Turn on" fluorescence kinetics of mixtures of CalFluorAzide647 (2 μM) and the indicated compounds (200 μM). Azide-alkyne reactions were done in 90% PBS buffer pH=7.4 + 10% methanol. (B) "Turn on" fluorescence kinetics of mixtures of Atto488-H-tetrazine (2.5 μM) with the indicated compounds (2.5 mM except for TCO*K, which was added at 2.5 μM). Tetrazine ligations were done in 100% DMSO. In (A) and (B), the controls include the quenched dye only. All reactions were conducted at 20 °C.

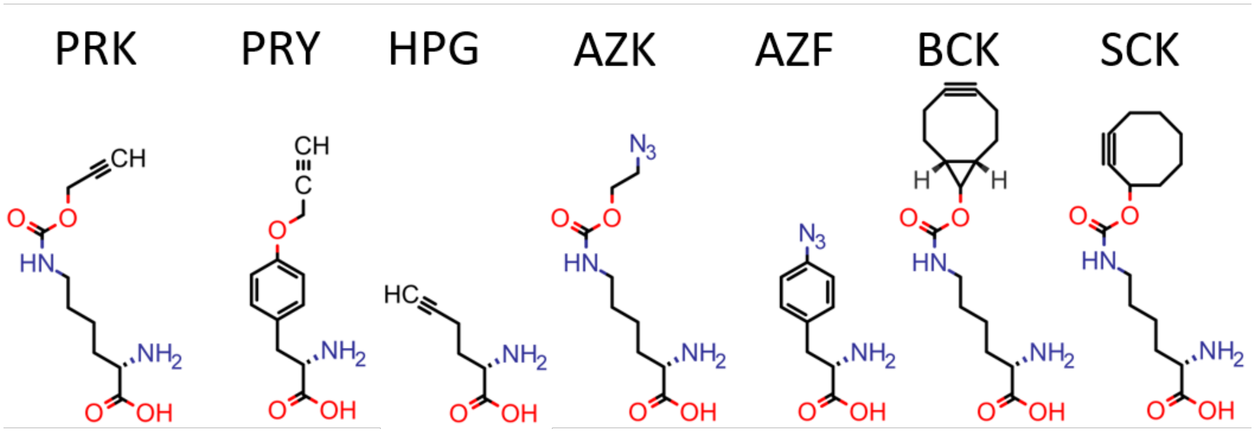

**Fig. S4. Non-canonical amino acids discussed in Fig. 2A**. PRK: Propargyl-lysine. PRY: Propargyl-tyrosine. HPG: Homopropargyl-glycine. AZK: Azido-lysine. AZF: 4-azido-phenylalanine. BCK: Bicyclononyne-lysine. SCK: Strained-cycloalkyne-lysine.

#### Note S3. Raman spectroscopy of "transparent window" probes.

The Raman intensity of an alkyne-containing molecular Raman probe is typically evaluated by its relative Raman intensity versus 2-ethynyl-deoxyuridine (EDU) or RIE. To account for the different number of repeating units, we next calculated the bond-normalized RIE value (RIE per C≡C bond) of various types of alkynes, including diynes, acetylenes, and cycloalkynes (**Fig. S5, Table S2**). We found that DAF has a similar bond-normalized RIE, 6, as the compound diphenylacetylene (DPA), but 6-fold more than phenylacetylene and 4-fold less than diphenylbutadiyne (DPY). Cyclic alkynes displayed the lowest relative Raman intensities among all the studied compounds.

**Table S2.**

| | **$\nu_{max}$ (cm$^{-1}$)** | | |
|---|---|---|---|
| **Compound<br>Solvent** | **PAE** | **DPA** | **DPY** |
| *Water* | 2177.283 | - | - |
| *DMSO* | 2170.399 | 2286.051 | 2286.445 |
| *Methanol* | - | 2289.578 | 2289.695 |
| *Ethanol* | 2178.809 | 2289.936 | 2289.636 |
| *Isopropanol* | 2179.886 | 2290.14 | 2290.104 |
| *Trifluoroethanol* | 2181.268 | 2289.917 | - |
| *Toluene* | 2178.134 | 2286.246 | 2286.08 |

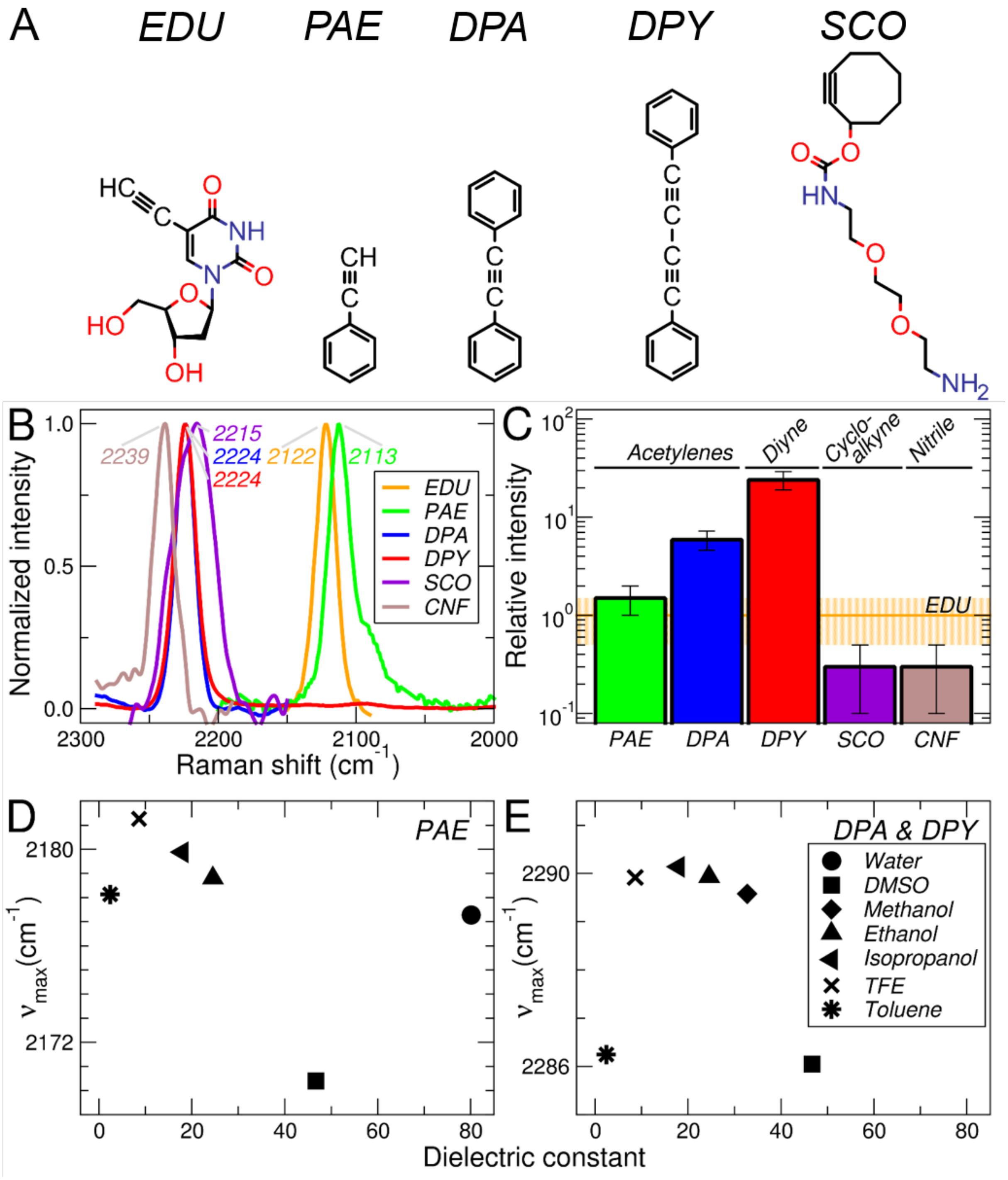

**Fig. S5. Raman activity of model compounds.** (A) Structure of the model compounds discussed in the paper. (B) Stimulated-Raman spectra. (C) Relative intensity vs. EDU (RIE). (D) Position of the Raman peak of PAE as a function of the solvent dielectric constant, E) Position of the Raman peak of DPA and DPY as a function of the solvent dielectric constant. EDU: ethynyl-deoxyuridine. PAE: phenylacetylene, DPA: diphenylacetylene. DPY: diphenylbutadiyne. SCO: Strained-cyclooctyne-PEG-amine.

The bond-normalized RIE values of DAF and alkyne-bearing ncAA (**Table S3**) suggest that aryl alkynes, like those in DAF and ETF, and nitriles, like that of CNF, provide sufficient signal-to-noise ratio for biological studies at millimolar concentrations. The peak positions ($\nu_{max}$) are also indicated in **Table S3**. Importantly, DAF and ETF display a notable frequency shift (~100 $cm^{-1}$ down-shift of ETF relative to DAF) suggesting that the two of them can be paired up in a single experiment.

Next, we measured the sensitivity of the alkyne frequency to the local chemical environment in mixed water/DMSO buffer solutions. Both nitriles and alkynes exhibit vibrational solvatochromism by shifting to lower energies with decreasing water content. We observed that DAF and ETF exhibit similar solvent-dependent shifts (7.3 $cm^{-1}$ in DAF vs. 7.6 $cm^{-1}$ in ETF, **Fig. 2D**). As a reference, the commonly used infrared reporter ncAA 4-cyano-phenylalanine (CNF) shifts by as much as 9 $cm^{-1}$ in going from water to DMSO (**Fig. 2D**). Interestingly, not only the peak position varied but also the peak area of DAF exhibited a clear increase as a function of the DMSO concentration (**Fig. 2E**). Such a response of DAF (and CNF) is in clear contrast with the behavior of ETF, which show similar spectral intensities regardless of the solvent (**Fig. 2E**).

Thus, two parameters can be extracted from the Raman spectra of DAF in the C≡C region: the peak positions (related to hydrogen bonding), and the peak areas. In the case of nitriles, their infrared peak areas are related to the local electric field.[6,7] Although the Raman spectra of alkynes has been extensively studied theoretically,[8] we are not aware of any quantitative interpretation of the intensity changes.

**Table S3.**

| **ncAA** | **Solvent** | **$\nu_{max}$ ($cm^{-1}$)** | **RIE** |
|---|---|---|---|
| *DAF* | Methanol | 2207 | 5.2 |
| *ETF* | 3 M NaOH | 2104 | 0.8 |
| *PRY* | 3 M NaOH | 2120 | 0.14 |
| *AZF* | 3 M NaOH | 2185 | 0.08 |
| *PRK* | 3 M NaOH | 2101 | 0.17 |
| *AZK* | 3 M NaOH | 2107 | 0.03 |
| *HPG* | 3 M NaOH | 2123 | 0.2 |
| *CNF* | Water | 2234 | 0.6 |
| *BCK* | 0.2 M NaOH + 15% DMSO | 2112 | 0.08 |
| *SCK* | 0.2 M NaOH + 15% DMSO | 2213 | 0.07 |
| *EDU* | water | 2115 | 1.0 |

#### Note S4. Sequences of all constructs used in the present study.

A) Orthogonal translation systems.

>*Mj*TyrRS* (Y32L, L65V, F108W, Q109M, D158G, I159P)

MDEFEMIKRNTSEIISEEELREVLKKDEKSALIGFEPSGKIHLGHYLQIKKMIDLQNAGFDIIIVLADLHAYLNQKGELDEIRK
IGDYNKKVFEAMGLKAKYVYGSEWMLDKDYTLNVYRLALKTTLKRARRSMELIAREDENPKVAEVIYPIMQVNGPHYL
GVDVAVGGMEQRKIHMLARELLPKKVVCIHNPVLTGLDGEGKMSSSKGNFIAVDDSPEEIRAKIKKAYCPAGVVEGNPI
MEIAKYFLEYPLTIKRPEKFGGDLTVNSYEELESLFKNKELHPMDLKNAVAEELIKILEPIRKRL

>*Mj*TyrtRNA$_{CUA}$

CCGGCGGTAGTTCAGCAGGGCAGAACGGCGGACTCTAAATCCGCATGGCGCTGGTTCAAATCCGGCCCGCCGGA
CCA

>*Mm*PylRS* (N346A, C348A, Y384F)

MDKKPLNTLISATGLWMSRTGTIHKIKHHEVSRSKIYIEMACGDHLVVNNSRSSRTARALRHHKYRKTCKRCRVSDEDL
NKFLTKANEDQTSVKVKVVSAPTRTKKAMPKSVARAPKPLENTEAAQAQPSGSKFSPAIPVSTQESVSVPASVSTSISSIS
TGATASALVKGNTNPITSMSAPVQASAPALTKSQTDRLEVLLNPKDEISLNSGKPFRELESELLSRRKKDLQQIYAEEREN
YLGKLEREITRFFVDRGFLEIKSPILIPLEYIERMGIDNDTELSKQIFRVDKNFCLRPMLAPNLYNYLRKLDRALPDPIKIFEIG
PCYRKESDGKEHLEEFTMLAFAQMGSGCTRENLESIITDFLNHLGIDFKIVGDSCMVFGDTLDVMHGDLELSSAVVGPIP
LDREWGIDKPWIGAGFGLERLLKVKHDFKNIKRAARSESYYNGISTNL

>*Mm*PyltRNA$_{CUA}$

GGAAACCTGATCATGTAGATCGAATGGACTCTAAATCCGTTCAGCCGGGTTAGATTCCCGGGGTTTCCGCCA

>*Ma*DafRS (named *Ma*PylRS* in **Fig. 1,** L125M, N166H, V168H, A223G, W239S)

MTVKYTDAQIQRLREYGNGTYEQKVFEDLASRDAAFSKEMSVASTDNEKKIKGMIANPSRHGLTQLMNDIADALVAE
GFIEVRTPIFISKDALARMTITEDKPLFKQVFWIDEKRALRPMLAPNMYSVMRDLRDHTDGPVKIFEMGSCFRKESHSG
MHLEEFTMLHLHDMGPRGDATEVLKNYISVVMKAAGLPDYDLVQEESDVYKETIDVEINGQEVCSAGVGPHYLDAAH
DVHEPSSGAGFGLERLLTIREKYSTVKKGGASISYLNGAKINSG

>*Ma*PyltRNA$_{CUA}$

AGATCTGGGGGACGGTCCGGCGACCAGCGGGTCTCTAAAACCTAGCATAGCGGGGTTCGACACCCCGGTCTCTCG

B) Green Fluorescent Protein (GFP) variants

>GFP-WT

MVSKGEELFTGVVPILVELDGDVNGHKFSVRGEGEGDATNGKLTLKFICTTGKLPVPWPTLVTTLTYGVQCFSRYPDHM
KRHDFFKSAMPEGYVQERTISFKDDGTYKTRAEVKFEGDTLVNRIELKGIDFKEDGNILGHKLEYNFNSHNVYITADKQK
NGIKANFKIRHNVEDGSVQLADHYQQNTPIGDGPVLLPDNHYLSTQSVLSKDPNEKRDHMVLLEFVTAAGITHGMDEL
YKGSHHHHHH

>GFP-N150DAF

MVSKGEELFTGVVPILVELDGDVNGHKFSVRGEGEGDATNGKLTLKFICTTGKLPVPWPTLVTTLTYGVQCFSRYPDHM
KRHDFFKSAMPEGYVQERTISFKDDGTYKTRAEVKFEGDTLVNRIELKGIDFKEDGNILGHKLEYNFNSH$VYITADKQK
NGIKANFKIRHNVEDGSVQLADHYQQNTPIGDGPVLLPDNHYLSTQSVLSKDPNEKRDHMVLLEFVTAAGITHGMDEL
YKGSHHHHHH

C) EL222 variants

>EL222-WT (named WT-EL222 in **Fig. 4**).

GADDTRVEVQPPAQWVLDLIEASPIASVVSDPRLADNPLIAINQAFTDLTGYSEEECVGRNCRFLAGSGTEPWLTDKIR
QGVREHKPVLVEILNYKKDGTPFRNAVLVAPIYDDDDELLYFLGSQVEVDDDQPNMGMARRERAAEMLKTLSPRQLE
VTTLVASGLRNKEVAARLGLSEKTVKMHRGLVMEKLNLKTSADLVRIAVEAGIGSENLYFQ

>EL222-L35DAF/V217C (named XL-EL222 in **Fig. 4**).

GADDTRVEVQPPAQWVLD**$**IEASPIASVVSDPRLADNPLIAINQAFTDLTGYSEEECVGRNCRFLAGSGTEPWLTDKIR
QGVREHKPVLVEILNYKKDGTPFRNAVLVAPIYDDDDELLYFLGSQVEVDDDQPNMGMARRERAAEMLKTLSPRQLE
VTTLVASGLRNKEVAARLGLSEKTVKMHRGLVMEKLNLKTSADLCRIAVEAGIGSENLYFQ

>EL222-L35DAF

GADDTRVEVQPPAQWVLD**$**IEASPIASVVSDPRLADNPLIAINQAFTDLTGYSEEECVGRNCRFLAGSGTEPWLTDKIR
QGVREHKPVLVEILNYKKDGTPFRNAVLVAPIYDDDDELLYFLGSQVEVDDDQPNMGMARRERAAEMLKTLSPRQLE
VTTLVASGLRNKEVAARLGLSEKTVKMHRGLVMEKLNLKTSADLVRIAVEAGIGSENLYFQ

>EL222-M151DAF

GADDTRVEVQPPAQWVLDLIEASPIASVVSDPRLADNPLIAINQAFTDLTGYSEEECVGRNCRFLAGSGTEPWLTDKIR
QGVREHKPVLVEILNYKKDGTPFRNAVLVAPIYDDDDELLYFLGSQVEVDDDQPN**$**GMARRERAAEMLKTLSPRQLEV
TTLVASGLRNKEVAARLGLSEKTVKMHRGLVMEKLNLKTSADLVRIAVEAGIGSENLYFQ

>EL222-I225DAF

GADDTRVEVQPPAQWVLDLIEASPIASVVSDPRLADNPLIAINQAFTDLTGYSEEECVGRNCRFLAGSGTEPWLTDKIR
QGVREHKPVLVEILNYKKDGTPFRNAVLVAPIYDDDDELLYFLGSQVEVDDDQPNMGMARRERAAEMLKTLSPRQLE
VTTLVASGLRNKEVAARLGLSEKTVKMHRGLVMEKLNLKTSADLVRIAVEAG**$**GSENLYFQ

>EL222-M151ETF

GADDTRVEVQPPAQWVLDLIEASPIASVVSDPRLADNPLIAINQAFTDLTGYSEEECVGRNCRFLAGSGTEPWLTDKIR
QGVREHKPVLVEILNYKKDGTPFRNAVLVAPIYDDDDELLYFLGSQVEVDDDQPNØGMARRERAAEMLKTLSPRQLEV
TTLVASGLRNKEVAARLGLSEKTVKMHRGLVMEKLNLKTSADLVRIAVEAGIGSENLYFQ

D) Maltose-binding protein (MBP) variants

>MBP-WT
MSKIKHHHHHHGTGMKIEEGKLVIWINGDKGYNGLAEVGKKFEKDTGIKVTVEHPDKLEEKFPQVAATGDGPDIIFWA
HDRFGGYAQSGLLAEITPDKAFQDKLYPFTWDAVRYNGKLIAYPIAVEALSLIYNKDLLPNPPKTWEEIPALDKELKAKGK
SALMFNLQEPYFTWPLIAADGGYAFKYENGKYDIKDVGVDNAGAKAGLTFLVDLIKNKHMNADTDYSIAEAAFNKGET
AMTINGPWAWSNIDTSKVNYGVTVLPTFKGQPSKPFVGVLSAGINAASPNKELAKEFLENYLLTDEGLEAVNKDKPLGA
VALKSYEEELAKDPRIAATMENAQKGEIMPNIPQMSAFWYAVRTAVINAASGRQTVDEALKDAQTTSGENLYFQGSSL
RIRAPSTKLWSHPQFEK

>MBP-V38DAF
MSKIKHHHHHHGTGMKIEEGKLVIWINGDKGYNGLAEVGKKFEKDTGIKVT**$**EHPDKLEEKFPQVAATGDGPDIIFWA
HDRFGGYAQSGLLAEITPDKAFQDKLYPFTWDAVRYNGKLIAYPIAVEALSLIYNKDLLPNPPKTWEEIPALDKELKAKGK
SALMFNLQEPYFTWPLIAADGGYAFKYENGKYDIKDVGVDNAGAKAGLTFLVDLIKNKHMNADTDYSIAEAAFNKGET
AMTINGPWAWSNIDTSKVNYGVTVLPTFKGQPSKPFVGVLSAGINAASPNKELAKEFLENYLLTDEGLEAVNKDKPLGA
VALKSYEEELAKDPRIAATMENAQKGEIMPNIPQMSAFWYAVRTAVINAASGRQTVDEALKDAQTTSGENLYFQGSSL
RIRAPSTKLWSHPQFEK

>MBP-V38ETF
MSKIKHHHHHHGTGMKIEEGKLVIWINGDKGYNGLAEVGKKFEKDTGIKVT**Ø**EHPDKLEEKFPQVAATGDGPDIIFWA
HDRFGGYAQSGLLAEITPDKAFQDKLYPFTWDAVRYNGKLIAYPIAVEALSLIYNKDLLPNPPKTWEEIPALDKELKAKGK
SALMFNLQEPYFTWPLIAADGGYAFKYENGKYDIKDVGVDNAGAKAGLTFLVDLIKNKHMNADTDYSIAEAAFNKGET
AMTINGPWAWSNIDTSKVNYGVTVLPTFKGQPSKPFVGVLSAGINAASPNKELAKEFLENYLLTDEGLEAVNKDKPLGA
VALKSYEEELAKDPRIAATMENAQKGEIMPNIPQMSAFWYAVRTAVINAASGRQTVDEALKDAQTTSGENLYFQGSSL
RIRAPSTKLWSHPQFEK

**$** and **Ø** represent here DAF and ETF respectively.
The 3 amino acids that form the chromophore of GFP upon maturation are shown in green

E) DNA

> dsDNA
Forward sequence or biotinylated single-stranded DNA (b-ssDNA):
biotin–5'–TTT TTT TTT TTT TTA GGT AGC CTT TAG TCC ATG–3'
Reverse or complementary sequence (cDNA): 5'–CAT GGA CTA AAG GCT ACC TA–3'

### Note S5. Protein quality control.

**Table S4. Masses and concentrations of the protein constructs used in this study.**

| Proteins | Expected mass (g/mol) | Observed mass (g/mol) $[M+H]^+$ | Yield (mg/L) | Concentration for steady-state FSRS (mM) | Concentration for time-resolved FSRS (mM) | Concentration for TA (μM) |
|---|---|---|---|---|---|---|
| EL222-WT | 24023.37 | 24024.3894 | 32.7 | 2.6 | N.A. | N.A. |
| EL222-L35DAF | 24105.3689 | 24106.4838 | 8.6 | 1.8 | 1.2 | 77 |
| EL222-M151DAF | 24087.4089 | 24088.4679 | 8.5 | 2.3 | 1.4 | 154 |
| EL222-M151DAF[a] | 23222.17 | 23375.1777* | | N.A. | N.A. | N.A. |
| EL222-M151ETF | 24063.54 | 24064.5440 | 9.8 | 1.5 | 1.1 | 77 |
| EL222-M151ETF[a] | 23196.16 | 23197.1629 | | N.A. | N.A. | N.A. |
| EL222-I225DAF | 24105.3689 | 24105.3398 | 4.8 | 1.5 | 1.3 | 22 |
| EL222-I225DAF-594Tetrazine | 24968.5905 | 24969.728 | N.A. | N.A | N.A | N.A |
| MBP-WT | 45254.13 | 45255.3542 | 28.2 | N.A. | N.A. | N.A. |
| MBP-V38DAF | 45350.147 | 45351.3723 | 8.8 | N.A. | N.A. | N.A. |
| MBP-V38ETF | 45326.76 | 45327.3766 | 10.4 | N.A. | N.A. | N.A. |

*This mass corresponds to the mass of an adduct between EL222M151DAF and DTT.

[a] From a construct containing an intein-CBD-12His tag

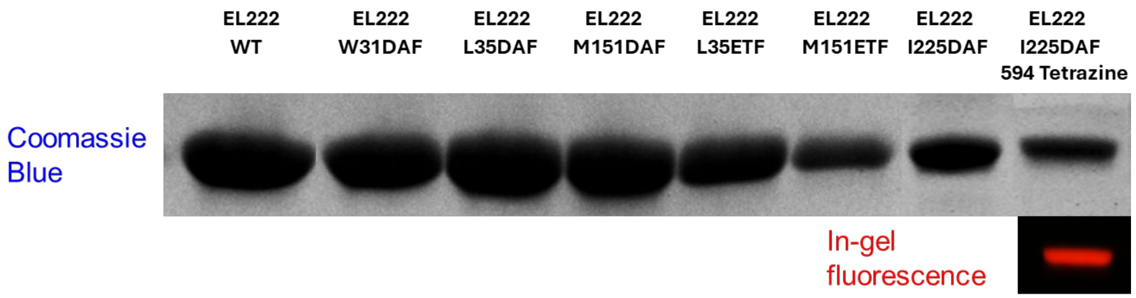

**Fig. S6. SDS-PAGE of the protein constructs used in this study.** (A) Coomassie Blue-stained SDS-PAGE of EL222 WT and EL222 mutants. (B) In-gel fluorescence of labeled EL222 I225DAF after incubation with AZDye594 Tetrazine and removal of excess dye. In-gel fluorescence was acquired with $\lambda_{excitation}$ = 650 nm, $\lambda_{emission}$ ~ 750 nm in order to visualize the fluorescence arising from AZDye594.

#### Note S6. Directed evolution of a DAF-specific synthetase.

The aminoacyl synthetase for incorporation DAF into protein was obtained from the life-death selection procedure on the 5-site library of *Methanomethylophilus alvus* pyrrolysyl-tRNA synthetase (*Ma*PylRS),[9] in which 5 residues at the amino acid binding pocket were randomized.[10] This election procedure includes one round positive and one round negative selection according to established protocol.[11] With the supplement of DAF in the positive selection, library members that can aminoacylate the orthogonal tRNA with either canonical amino acid or DAF present in the bacteria cell are collected. Subsequently, the negative selection in absence of DAF removes unwanted library members that aminoacylate the orthogonal tRNA with canonical amino acid. The fluorescence-activated cell sorting (FACS)-based screening is employed to assess how well the selective library accomplishes the ncAA-dependent suppression of TAG codons, based on comparing the fluorescent expression of superfolder GFPN150TAG protein in cell with and without DAF supplement. The bacteria containing the most promising *Ma*PylRS DAF variants are sorted and validated individually. As a result, the *Ma*PylRS DAF is achieved.

During positive selection, DH10B cells were electroporated with pBK-MaPylRS library plasmid (Addgene #224585) and pPOS plasmid (Addgene #197571), which contains a TAG mutation in chloramphenicol resistance gene. Transformed cells were grown on autoinduced media (AIM) agar plate supplemented with 0.5 mM DAF, chloramphenicol (Cam), kanamycin (Kan) and tetracycline (Tet) at 37 °C, overnight. All resulting colonies were collected and their pBK library plasmids were isolated. For negative selection, the isolated pBK plasmids were electroporated into DH10B cells harboring pNEG plasmid (Addgene #197572), which contains a TAG mutation in toxic barnase gene. Cells were grown on AIM agar plate supplemented with Cam, Kan at 37 °C, overnight. Colonies again were collected. The enriched pBK plasmids were isolated and electroporated into DH10B cell containing pALS3-sfGFP-N150TAG (addgene #212121). Recovered transformants were grown in non-induced media (NIM) supplemented with Kan, Tet at 37 °C, shaking 250 rpm, overnight. The following day, the protein expression was induced in AIM supplemented with either without DAF or with 0.5 mM DAF, Kan, Tet, at 37 °C, shaking 250 rpm, overnight. Cells of 1 mL induced culture were harvested, washed twice with 1 mL of 1X PBS buffer and resuspended to a final density of $10^6$–$10^7$ cells/mL in a total volume of 10 mL ready for FACS sorting analysis (**Fig. S7A)**. Cells supplemented with DAF and corresponding to the top 5% most fluorescent events were sorted and the mixture of pBK enriched plasmids and pALS3-sfGFP-N150TAG plasmid were extracted.

To evaluate the efficiency and fidelity of individual synthetases, the extracted plasmid mixture was electroporated into commercial electroMAX DH10B cells (Invitrogen™). In parallel, the pALS3-sfGFP-WT plasmid, generated by reverting the TAG mutation in pALS3-sfGFP-N150TAG to the wild-type sequence, was transformed as a positive control. Transformed cells were grown on AIM agar plate supplemented with either without DAF or with 0.5 mM DAF, Kan, Tet at 37 °C for 2 days until the colonies and their GFP fluorescence are seen by naked eyes under visible normal light. These plates were imaged for fluorescence using an Azure c600 imaging system (**Fig. S7B).**

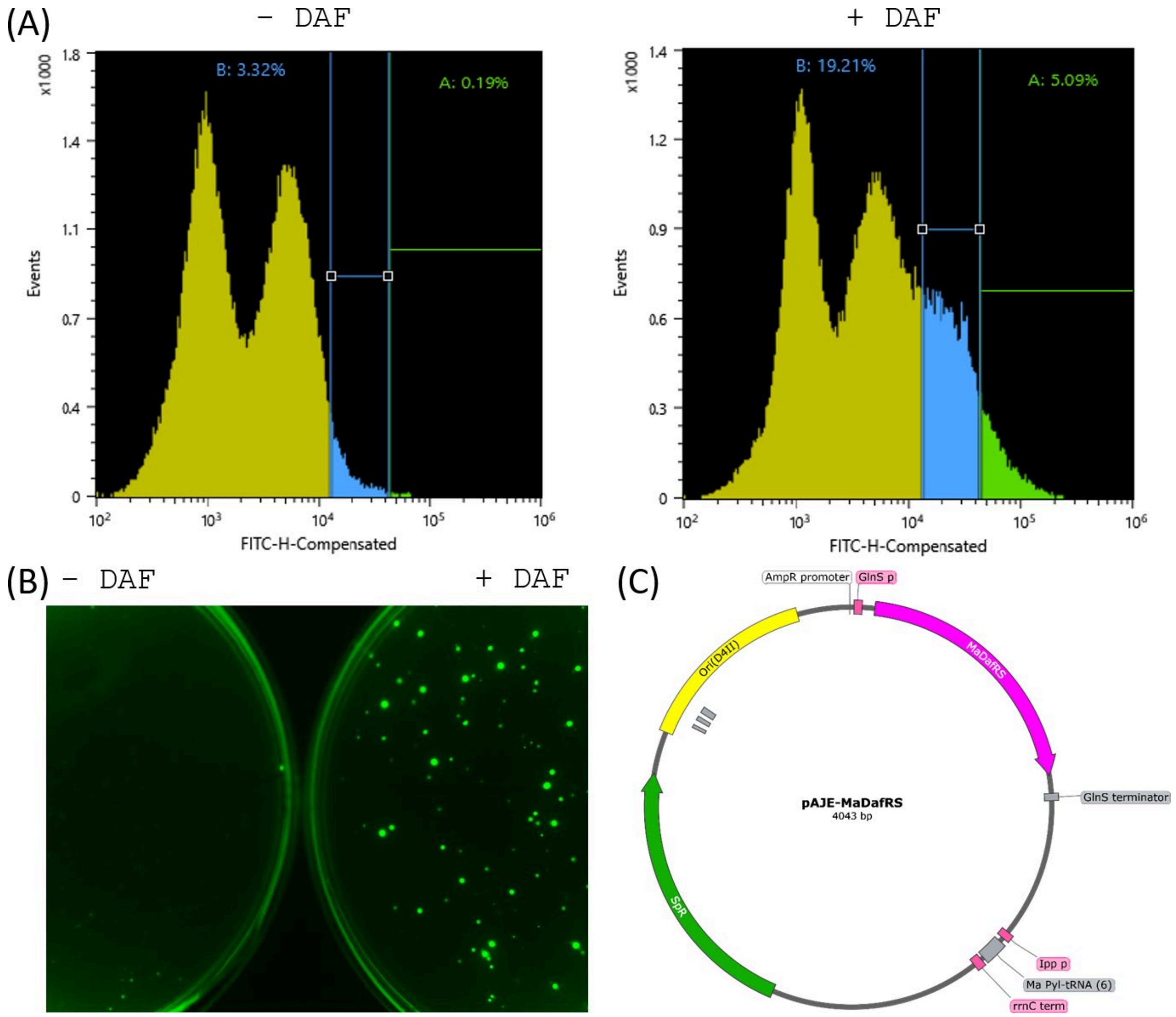

**Fig. S7. Directed evolution of a DAF-specific *Ma*PylRS-derived orthogonal translation system.** A) Fluorescence-activated cell sorting analysis of DH10B cells containing selected pBK-*Ma*PylRS variants and sfGFP-N150TAG exhibited fluorescence in autoinduction media (AIM) in the absence (*left*), and presence of DAF (*right*). B) Fluorescence image of *E. coli* colonies grown on AIM agar plates. DH10B cells harboring selected pBK-*Ma*PylRS variants and sfGFP-N150TAG exhibited virtually no fluorescence on AIM agar plates in the absence of DAF (*left*), whereas clear fluorescence was observed on plates containing DAF (*right*). C) Map of pAJE-*Ma*DafRS plasmid.

95 small and big colonies with brightest fluorescence on AIM supplemented with 0.5 mM DAF, Kan, Tet and a sfGFP-WT colony were picked and grown individually in 96-well block NIM supplemented with Kan, Tet at 37 °C, 250 rpm, overnight. Protein expression was induced in AIM supplemented with Kan and Tet, either in the presence or absence of 0.5 mM DAF, at 37 °C, shake 250 rpm, overnight. Cell suspensions were diluted 20-fold in PBS buffer, pH 7.4 to achieve an OD below 0.8 and transferred to black 96-well plates for fluorescence measurement using a CLARIOstar plate reader machine (BMG LABTECH). 17 promising clones exhibiting the highest fluorescence intensities were selected for pBK plasmid extraction and sequencing **(Table S5).**

**Table S5. Comparison of five amino acid–binding pocket residues among 17 MaPylRS variants exhibiting high fluorescence intensity, using the *Ma*PylRS sequence (WT) as the reference**. Eight individual colonies show same Group 1 sequence, eight individual colonies show same Group 2 sequence, and one colony shows Group 3 sequence.

| *Residue number* | 125 | 166 | 168 | 223 | 239 |
|---|---|---|---|---|---|
| ***WT*** | **L** | **N** | **V** | **A** | **W** |
| *Group 1 (8 individual clones)* | M | H | H | G | S |
| *Group 2 (8 individual clones)* | L | H | H | A | S |
| *Group 3 (1 clone)* | I | H | H | C | S |

Three synthetases representing 3 groups were further characterized in larger-scale 50 mL cultures. Firstly, the corresponding *Ma*PylRS variants were subcloned from the pBK backbone into the pAJE backbone (Addgene #225684) by NdeI and PstI restriction enzyme cloning. Each pAJE-*Ma*PylRS construct was then electroporated together with pET28a-sfGFP-N150TAG (Addgene #85493) into BL21AI cells for protein expression. Expression cultures were grown in TB medium supplemented with 0.25 mM DAF, IPTG, arabinose, Kan, and spectinomycin (Spec) at 37 °C, shaking at 180 rpm, for 20 h. The *Ma*DafRS (L125M, N166H, V168H, A223G, W239S) is our final synthetase for specific incorporation DAF into protein because it gives the highest yields and fidelities highest among all other synthetase (**Table S6**).

**Table S6.** Relative yields and fidelities of three selected synthetases.

| Parameter→<br>aaRS ↓ | Relative yield[1] (%) | Relative fidelity[2] (%) |
|---|---|---|
| *Ma*DafRS | 97 | 94 |
| *Mj*TyrRS* | 42 | 9 |
| *Mm*PylRS* | 8 | 0 |

[1]Relative yield is calculated as (mg or purified GFP-N150X in presence of DAF per liter of culture)/(mg of purified GFP-WT per liter of culture).
[2]Relative fidelity is estimated as (mg or purified GFP-N150X in presence of DAF per liter of culture - mg or purified GFP-N150X in the absence of DAF per liter of culture)/(mg of purified GFP-WT per liter of culture). X= DAF and/or F

The plasmid map encoding the *Ma*DafRS/*Ma*PyltRNA$_{CUA}$ pair is shown in **Fig. S7C** (the plasmid itself is available from Addgene #256615).

#### Note S7. Monitoring the thiol-yne reaction with mass spectrometry.

Protein–thiol adduct formation after in vitro thiol–yne reactions between purified EL222 M151DAF/ETF mutants and DTT was examined by mass spectrometry. The expected adduct for the DAF mutant was detected with 100% relative abundance, while the corresponding ETF mutant adduct was observed at only 3.4% relative abundance (**Fig. S8**).

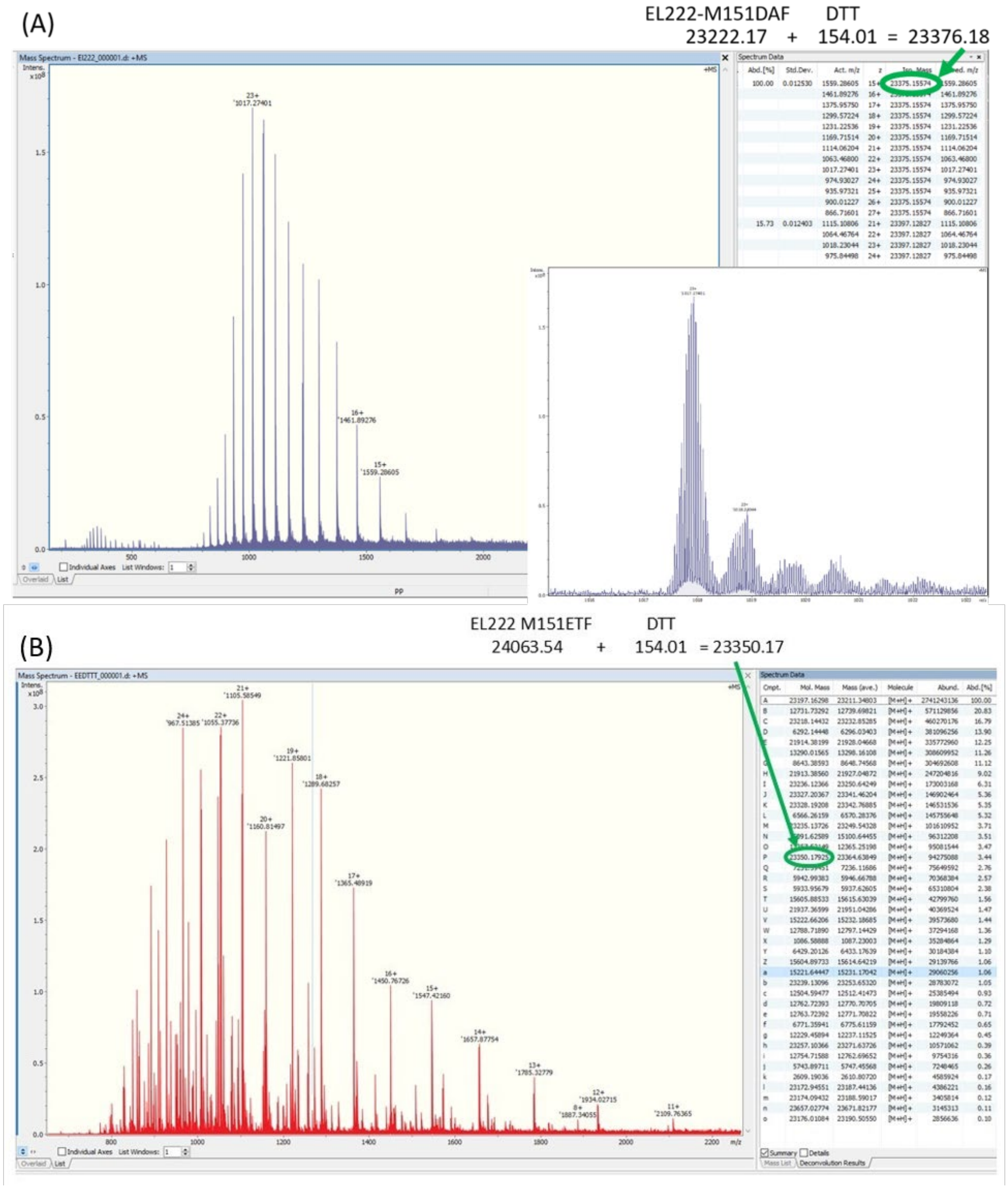

**Fig. S8. Mass spectra of EL222-M151X-Intein-CBD-12His after intein cleavage.** Reactions were carried out in the presence of 100 mM DTT for 48 hours at 23ºC. (A) X=DAF. (B) X=ETF.

#### Note S8. Labeling of ETF-containing proteins.

Similar to DAF, the labeling experiments in solution and in live *E. coli* cells were performed with purified MBPV38ETF. For in-vitro experiments, the protein was incubated with red fluorescent dyes Cy5 tetrazines (H-tetrazines and methyl-tetrazines), Cy5 azide (canonical and azide plus) and Cy5 thiol. The reactions were analyzed by SDS-PAGE followed by in-gel fluorescence (**Fig. S9B**). ETF mutant showed clear reactivity against AzidePlus in presence of catalyst and H-tetrazine dyes, but it was lower compared to that of DAF. However, no fluorescent band was observed upon reactions with methyl-tetrazines and azide. In case of thiol labelling with EL222M151ETF, additionally, after incubating the protein with dye, we removed the excess dye by gel filtration chromatography, and the UV/Visible spectrum was recorded (**Fig. S9B right**). The degree of labeling was estimated to be less than 10% which is again significantly lower compared to DAF (**Fig. S9C right**).

Next, we tested ETF reactivity in live bacterial cells by flow cytometry. *E. coli* cells expressing MBP-V38ETF proteins were incubated with green fluorescent dyes AZDye488 having three different reactive handles azidePlus, tetrazine and thiol. After incubation and washing out the excess dyes, Fluorescence -positive cells were quantified by flow cytometry. The higher percentage of fluorescent cells of cells expressing MBP-V38ETF relative to the negative controls were only found in case of CuAAC labelling with AzidePlus. However, labelling through H-tetrazines and thiols did not show significant higher fluorescence compared to the negative controls suggesting ETF is not an optimal handle for in-cell tetrazine or thiol ligation.

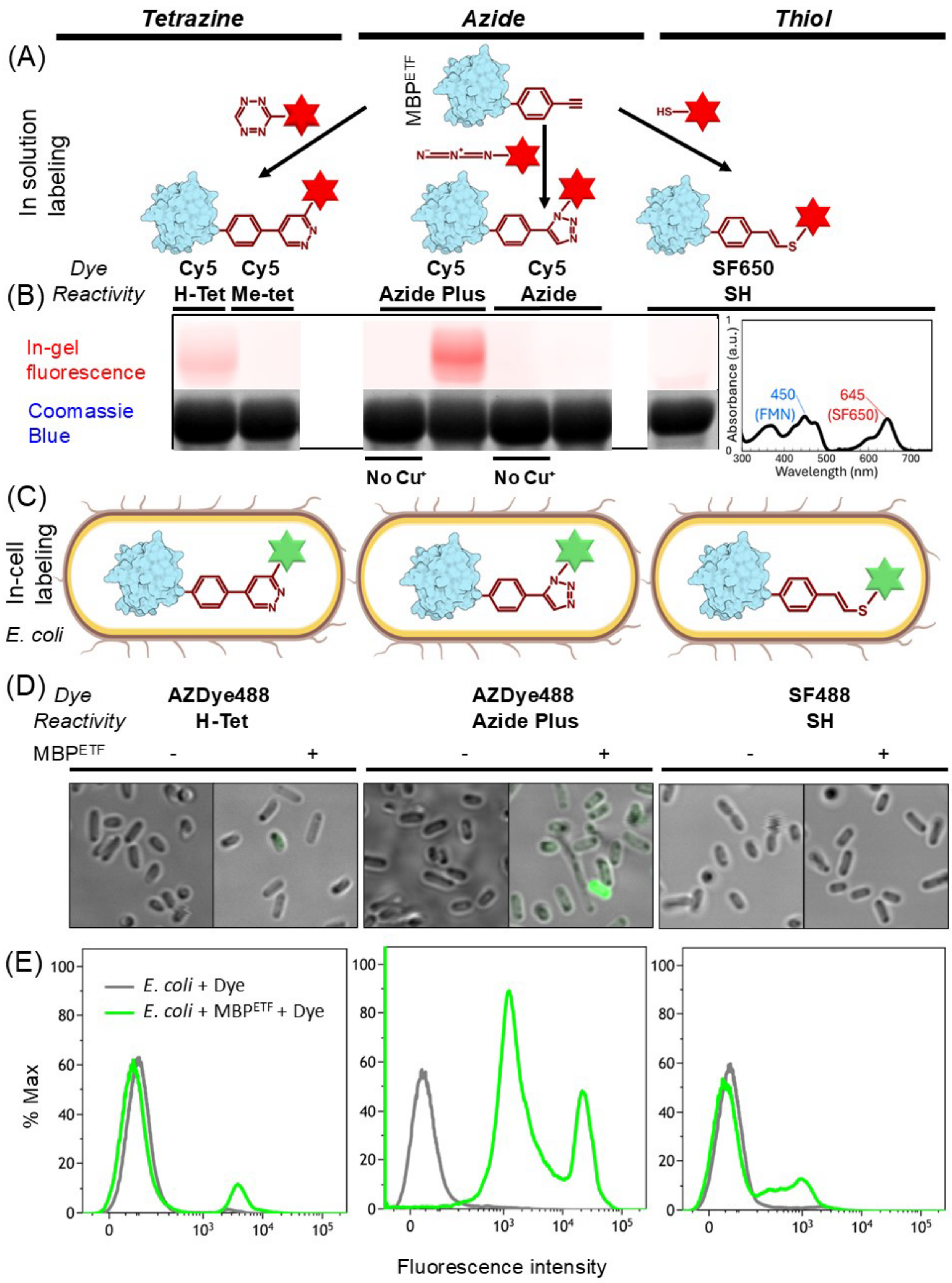

Tetrazine
Azide
Thiol
(A)
In solution labeling
MBPETF
Dye
Reactivity
Cy5 H-Tet
Cy5 Me-tet
Cy5 Azide Plus
Cy5 Azide
SF650 SH
(B)
In-gel fluorescence
Coomassie Blue
No Cu+
No Cu+
450 (FMN)
645 (SF650)
Absorbance (a.u.)
Wavelength (nm)
(C)
In-cell labeling
E. coli
(D)
AZDye488 H-Tet
AZDye488 Azide Plus
SF488 SH
MBPETF
(E)
E. coli + Dye
E. coli + MBPETF + Dye
% Max
Fluorescence intensity

**Fig. S9. Labeling of ETF-containing proteins with fluorescent dyes in solution and *in cellulo* (previous page).** (A) Scheme showing the reaction between purified MBP bearing ETF at residue 38 ($MBP^{ETF}$, originally expressed in *E. coli*, and a red-fluorescing dye carrying azide, tetrazine (H-tetrazine or Methyl-tetrazine), or thiol (SH) moieties. (B) Fluorescence image ($\lambda_{excitation}$ = 650 nm, $\lambda_{emission}$ ~ 750 nm) and corresponding Coomassie Blue-stained SDS-PAGE of 30 µM $MBP^{DAF}$ incubated with ten-fold excess of red-fluorescing dyes bearing distinct reactive handles for 16 hours. Azide reactions were done both in the absence and in the presence of copper ($Cu^+$) as a catalyst to check for SPAAC and CUAAC reaction mechanisms, respectively. In the case of thiol-yne reaction, the absorbance spectrum of 1 mM purified $EL222^{ETF}$ after incubation with four-fold excess of SF650-SH for 16 hours and removal of excess dye is shown on the right. The peak maxima in nanometer units of FMN and SF650 are indicated. (C) Scheme showing the reaction between maltose binding protein (MBP) bearing ETF at position 38 ($MBP^{DAF}$) recombinantly expressed in *E. coli* BL21(DE3) and exogenously added green-fluorescing dyes containing azide, tetrazine or sulfhydryl groups. (D) Confocal fluorescence images ($\lambda_{excitation}$ = 488 nm, $\lambda_{emission}$ ~575 nm) from of *E.coli* cells expressing $MBP^{ETF}$ incubated with green fluorescent dyes (250 µM H-tetrazine, 100 µM AzidePlus and 50 µM thiol) for 30 minutes. The scale bar is indicated on the left panel. (E) Flow cytometry histograms showing the fluorescence intensity ($\lambda_{excitation}$ = 488 nm, $\lambda_{emission}$ ~ 530 nm) distribution of cells within the gated *E.coli* population after the three labelling reactions depicted in (C). As a control to check the bio-orthogonality of the reactions, fluorescent dyes were also mixed with *E. coli* cells in the absence of DAF. The three tested reactions are shown in distinct sides of the panels: tetrazine-alkyne ligations (*left*), azide-alkyne cycloadditions (*middle*) and thiol-yne (*right*).

#### Note S9. Labelling of wild type proteins.

As a controlled experiment we performed the labeling of MBP WT in solution and in live *E. coli* cells (**Fig. S10** and **Fig. S11**) with a red-fluorescing dye carrying either azide, tetrazine, or thiol moieties. Only highly reactive Cy5 Tetrazine showed little reactivity towards the protein, but this reactivity is much slower than that of two EL222 mutants.

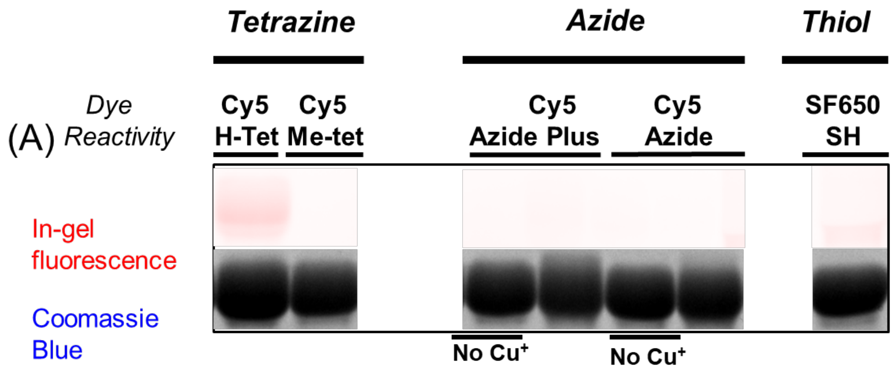

**Fig. S10. Labeling of WT proteins with fluorescent dyes in solution and *in cellulo*.** (A) Fluorescence image and corresponding Coomassie Blue-stained SDS-PAGE of MBP$^{WT}$ incubated with red-fluorescing dyes bearing distinct reactive handles for the indicated times.

#### Note S10. Additional flow cytometry experiments in *E. coli* cells.

DAF showed substantially higher reactivity toward H-tetrazine compared with ETF. Cells expressing MBP-DAF exhibited 45% labeled cells with a median fluorescence intensity of 6,667, whereas MBP-ETF cells showed only 13.08% labeled cells with a median fluorescence intensity of 3,821 (**Fig. S12 and table S7**). In contrast, all control samples displayed similar low-intensity peak patterns, with 0–14% labeled cells and median fluorescence intensities of approximately 3,000. Notably, MBP-WT cells showed ~12% labeled cells with a median intensity of 3,200, indicating background fluorescence, while cells without protein expression exhibited minimal labeling (2.8%). These results demonstrate that DAF reacts efficiently with tetrazine under intracellular conditions.

Similarly, DAF displayed higher reactivity toward thiol reagents compared with ETF. Cells expressing MBP-DAF showed 61% labeled cells with a median fluorescence intensity of 923, whereas MBP-ETF cells showed 27.8% labeled cells with a median fluorescence intensity of 699. The remaining control samples exhibited comparable low-intensity distributions, with 0–20% labeled cells and median fluorescence intensities around 652. MBP-WT cells showed ~20% labeled cells with a median intensity of 590, suggesting nonspecific background labeling, while cells alone showed only 7.5% labeling. Together, these results indicate that DAF reacts effectively with thiol-based probes.

DAF also demonstrated stronger reactivity toward AzidePlus relative to ETF. Cells expressing MBP-DAF and displaying a single fluorescence population showed 64% labeled cells with a median fluorescence intensity of 5,996. In contrast, MBP-ETF cells exhibited two distinct fluorescence populations with an average labeling efficiency of 91% and a median fluorescence intensity of 1,883. To further interpret the origin of these two populations, additional control samples were analyzed.

Control cells containing only the RS/tRNA system together with ETF exhibited fluorescence profiles similar to one of the two populations observed in MBP-ETF cells, with labeling efficiencies of 99% (median 1,404). This observation suggests that one fluorescence population probably originates from residue ETF or ETF-associated RS/tRNA components reacting with AzidePlus, rather than from specifically labeled MBP-ETF protein. Consistent with this interpretation, the additional fluorescence population was also observed in ETF-treated controls-treated samples. Therefore, the dual-peak distribution observed for MBP-ETF indicates nonspecific or heterogeneous labeling, reducing its reliability for intracellular labeling applications.

Overall, DAF displays a strong reactivity towards H-tetrazines, AzidePlus, and thiols. Moreover, DAF consistently produces stronger and more reliable intracellular labeling than ETF, supporting its suitability for bacterial cell-labeling applications.

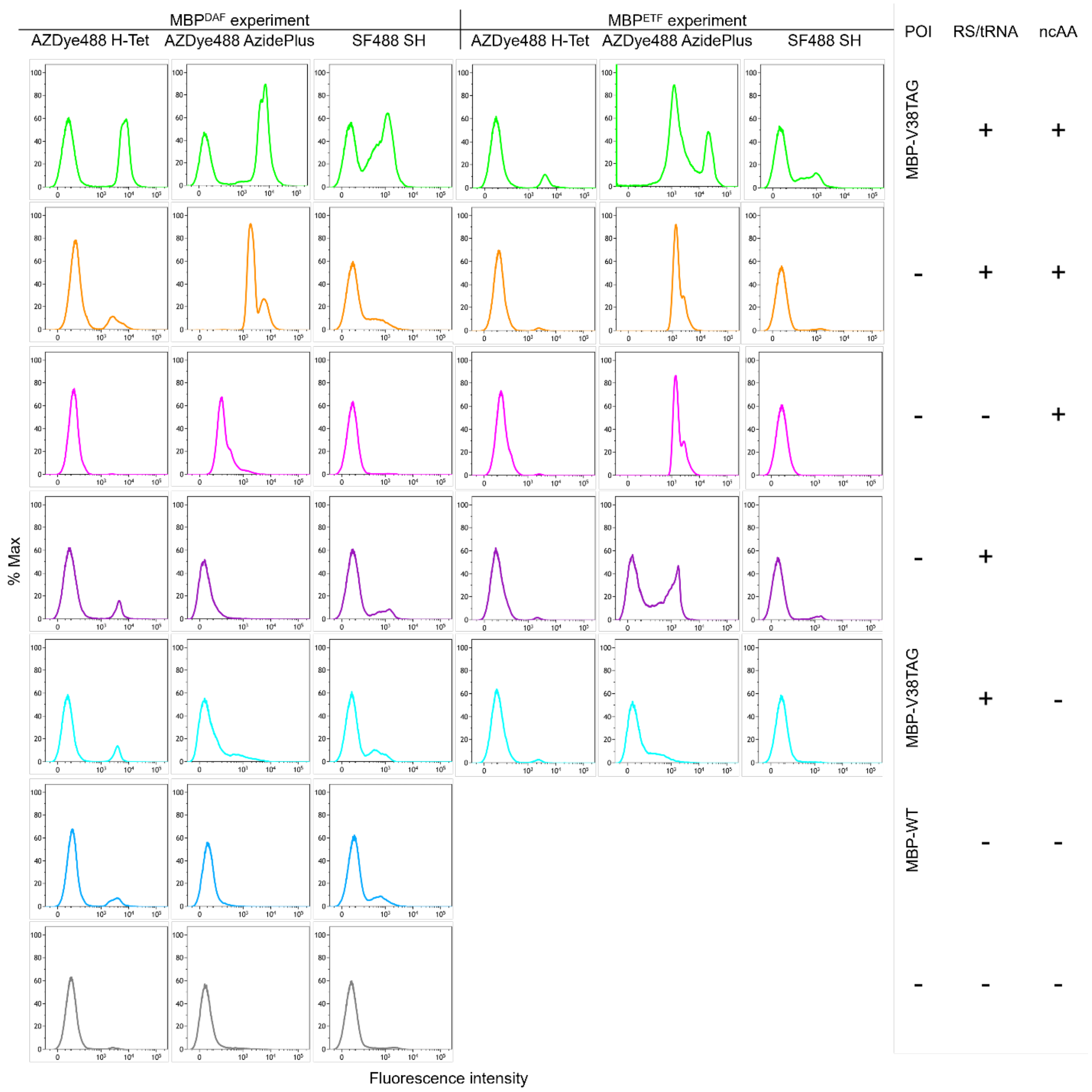

**Fig. S11: Representative flow cytometry histograms showing labeling profiles of different cell samples stained with various 488 fluorescent dyes.** Fluorescence intensity is displayed on the x-axis, while the y-axis represents normalized event frequency (% Max). Cells expressing MBP-V38DAF and the corresponding control cells were stained with 250 µM AZDye488 H-Tetrazine, 100 µM AZDye488 AzidePlus, and 50 µM SF488 SH dyes, as shown in the left panel. Cells expressing MBP-V38ETF and the corresponding control cells were stained with the same set of dyes, as shown in the right panel. POI = protein-of-interest. RS/tRNA = orthogonal translation system. NcAA = non-canonical amino acid

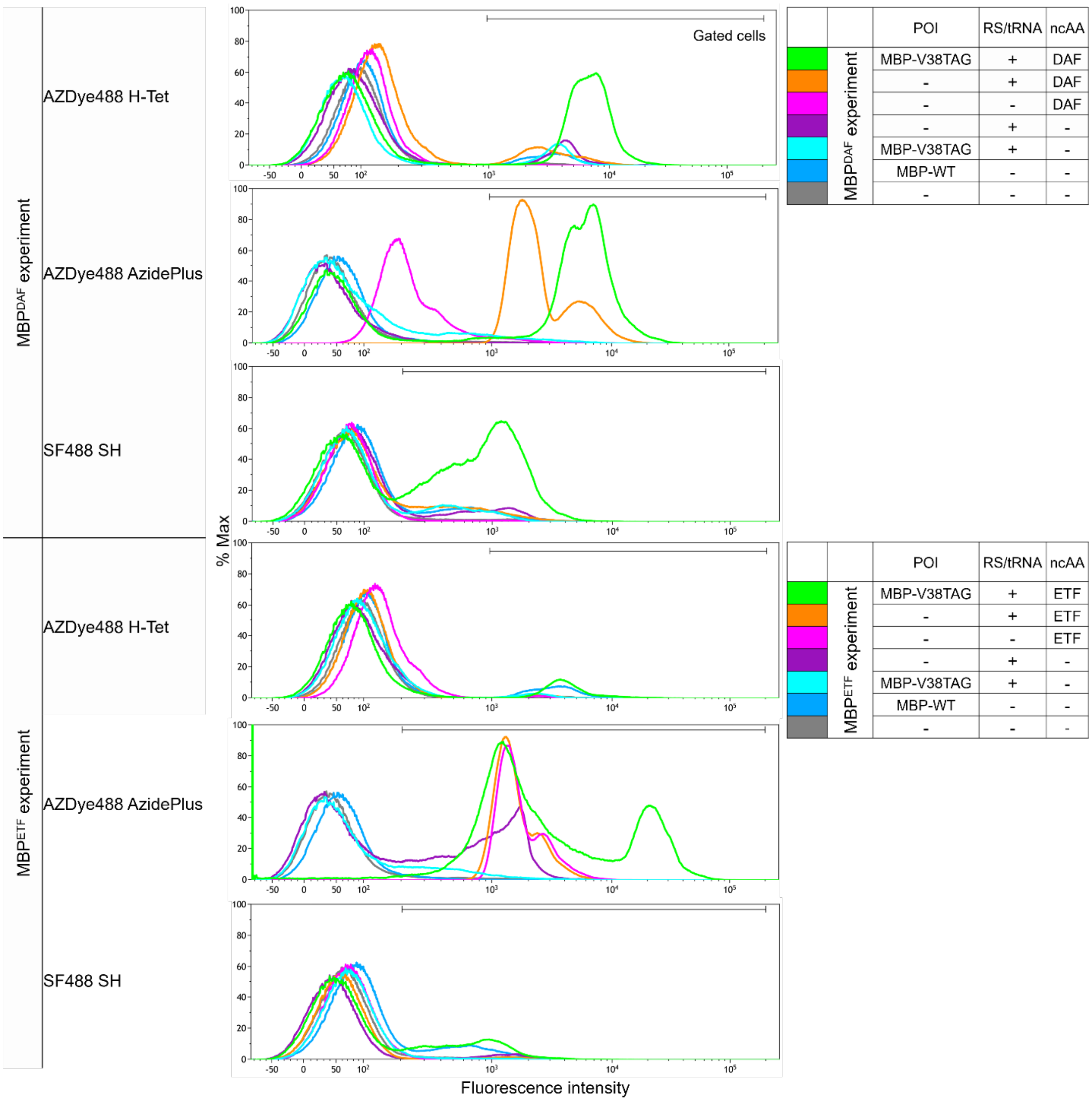

**Fig. S12: Overlay flow cytometry histograms demonstrating differences in staining intensity and fluorescence distribution among 488-labeled samples**. In the MBP$^{DAF}$ experiment (upper panel), cells expressing MBP-V38DAF and the corresponding control cells were stained with 250 µM AZDye488 H-tetrazine, 100 µM AZDye488 AzidePlus, and 50 µM SF488 SH dyes. In the MBP$^{ETF}$ experiment (lower panel), cells expressing MBPV38ETF and the corresponding control cells were stained using same set of dyes. POI = protein-of-interest. RS/tRNA = orthogonal translation system. NcAA = non-canonical amino acid.

**Table S7. The percentage of the gated cell population and the median fluorescence intensity (X-median) of all labeled samples (based on gating shown in Fig. S12).**

| | | | *Dye→* | **AZDye488 H-tetrazine** | | **AZDye488 AzidePlus** | | **SF488 SH** | |
|---|---|---|---|---|---|---|---|---|---|
| | *POI* | *RS/tRNA* | *ncAA* | *%Gated* | *X-Med* | *%Gated* | *X-Med* | *%Gated* | *X-Med* |
| $MBP^{DAF}$ | MBP-V38TAG | + | DAF | 45.33 | 6657.16 | 64.53 | 5996.95 | 60.81 | 923.23 |
| | - | + | DAF | 15.42 | 2916.36 | 99.44 | 2079.74 | 23.37 | 532.69 |
| | - | - | DAF | 0.8 | 2366.24 | 4.76 | 1518.62 | 3.6 | 585.2 |
| | - | + | - | 13.55 | 4122.03 | 1.47 | 2278.92 | 18.41 | 812.01 |
| | MBP-V38TAG | + | - | 13.84 | 3578.62 | 6.24 | 1888.6 | 20.18 | 507.33 |
| | MBP-WT | - | - | 12.09 | 3259.68 | 0.32 | 1633.06 | 19.79 | 590.95 |
| | - | - | - | 2.84 | 2614.33 | 2.04 | 1944.27 | 7.52 | 782.55 |
| $MBP^{ETF}$ | MBP-V38TAG | + | ETF | 13.08 | 3821.74 | 91.56 | 1883.56 | 27.48 | 699.09 |
| | - | + | ETF | 2.74 | 2405.62 | 99.98 | 1404.87 | 4.58 | 959.68 |
| | - | - | ETF | 1.46 | 2397.41 | 99.96 | 1495.7 | 2.5 | 268.83 |
| | - | + | - | 2.29 | 2067.24 | 46.65 | 1010.98 | 7.61 | 1032.23 |
| | MBP-V38TAG | + | - | 3.52 | 2243.29 | 18.59 | 455.27 | 4.83 | 385.55 |
| | MBP-WT | - | - | 12.05 | 3266.05 | 4.01 | 304.79 | 19.79 | 590.95 |
| | - | - | - | 2.8 | 2625.27 | 6.22 | 570.37 | 7.52 | 782.55 |

#### Note S11. Additional imaging experiments in *E. coli* cells.

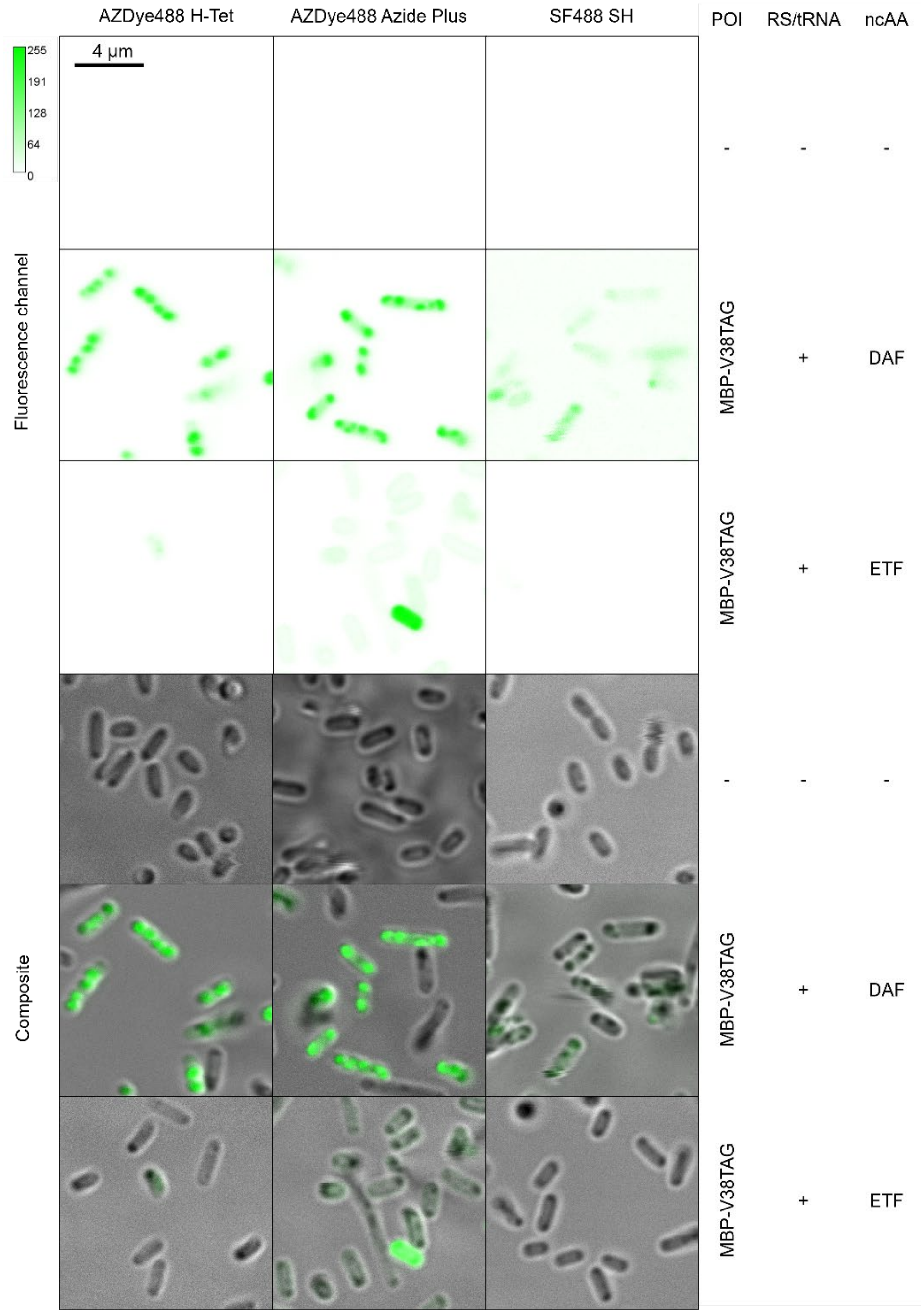

**Fig. S13. Fluorescence microscopy images of 488 dye-labeled bacterial cells.** The upper panel shows the fluorescence channel, illustrating the distribution of fluorescent dye labeling in bacterial cells. The lower panel shows the composite image generated by overlaying fluorescence and bright-field channels. In each panel, the first row shows control cells incubated with dye only, the second row shows cells expressing MBP-V38DAF incubated with dye, and the third row shows cells expressing MBP-V38ETF incubated with dye. Cells were stained with 250 µM AZDye 488 H-tetrazine, 100 µM AZDye488 AzidePlus, and 50 µM SF488 SH dyes. POI = protein-of-interest. RS/tRNA = orthogonal translation system. NcAA = non-canonical amino acid. The scale bars (length and fluorescence intensity) are shown on the top left panel.

#### Note S12. Overall reactivity and suitability of DAF.

**Table S8. Summary of the reactivity of DAF and the related ncAA ETF in different settings** (as free amino acid, genetically encoded in solution, genetically encoded in cells).

| REACTION & CHEMICALS | | DAF | | | ETF | | |
|---|---|---|---|---|---|---|---|
| *Reaction* | *Chemical handle* | Free ncAA[1] | In solution[2] | *In-cell*[3] | Free ncAA[1] | In solution[2] | *In-cell*[3] |
| **Tetrazine ligation** | H-tet | ++ | + | + | + | + | - |
| | M-tet | + | - | NP | - | - | NP |
| **Alkyne-azide cyclo-addition** | Azide (- $Cu^{+}$) | - | - | NP | - | - | NP |
| | Azide (+ $Cu^{+}$) | + | + | NP | + | - | NP |
| | AzidePlus (- $Cu^{+}$) | - | - | NP | - | - | NP |
| | AzidePlus (+ $Cu^{+}$) | NP | ++ | + | NP | + | -* |
| **Thiol-yne** | Thiol | ++ | ++ | + | + | + | - |

[1] 22 hours, pH= N.A. (organic solvent), conc. = 15 mM DAF/ETF + 20 mM second reactant

[2] 22 hours, Tris NaCl pH=8.0, conc. = 30 µM protein + 300 µM dye

[3] 30 min, PBS pH=7.4, [250 µM for tetrazine dye, 100 µM for AzidePlus dye and 30 µM for thiol dye, the intracellular protein concentration is unknown]

NP: Not performed

* heterogeneous labeling

**Table S9. Summary of the properties of DAF and related ncAA.** The properties include genetic encodability (yield and fidelity), (bio)chemical stability, reaction speed, physiological reaction efficiency, multi-reactivity, Raman signal strength, Raman solvatochromism, and overall suitability.

| NcAA → | DAF | ETF | CNF | TCO*K |
|---|---|---|---|---|
| Properties ↓ | 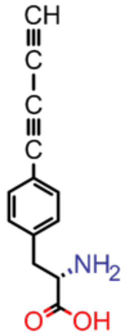 | 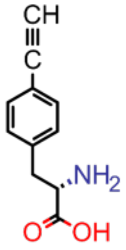 | 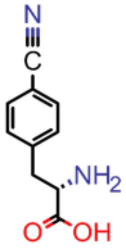 | 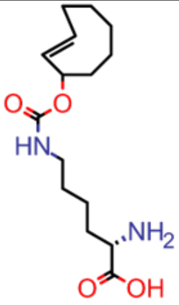 |
| *Encodability* | High | High | High | Medium |
| *Stability* | Medium | High | High | Medium |
| *Reaction speed* | Medium | Medium | N.A. | High |
| *Reaction efficiency (physiological)* | High | Low | N.A. | High |
| *Multi-reactivity* | High | Low | N.A. | Low |
| *Vibrational signal* | High | Medium | Medium | N.A. |
| *Vibrational solvatochromism* | High | High | High | N.A. |
| *Overall suitability* | High | Medium | Medium | Medium |

| High | Medium | Low | N.A. |
|---|---|---|---|

### Note S13. Mass spectrometry analysis of cross-linked EL222 (XL-EL222).

Mass spectrometry of the crosslinked EL222 was carried out with a bottom-up approach. The proteins were digested by trypsin and further analyzed by LC-MS/MS – timsTof Pro (Bruker Daltonics). Data were processed using DataAnalysis 6.1 (Bruker Daltonics) software. The data was searched against the single protein database EL222. An ion at m/z 941.22 (4+) m/z corresponding to the crosslinked peptide VEVQPPAQWVLD$IEASPIASVVSDPR - TSADLCR was identified. Our ncAA DAF is represented here as $. The cross-linked peptide was confirmed by CID fragmentation of ion at m/z 941.2265. Several 'b' and 'y' peptide fragments from both peptides were identified.

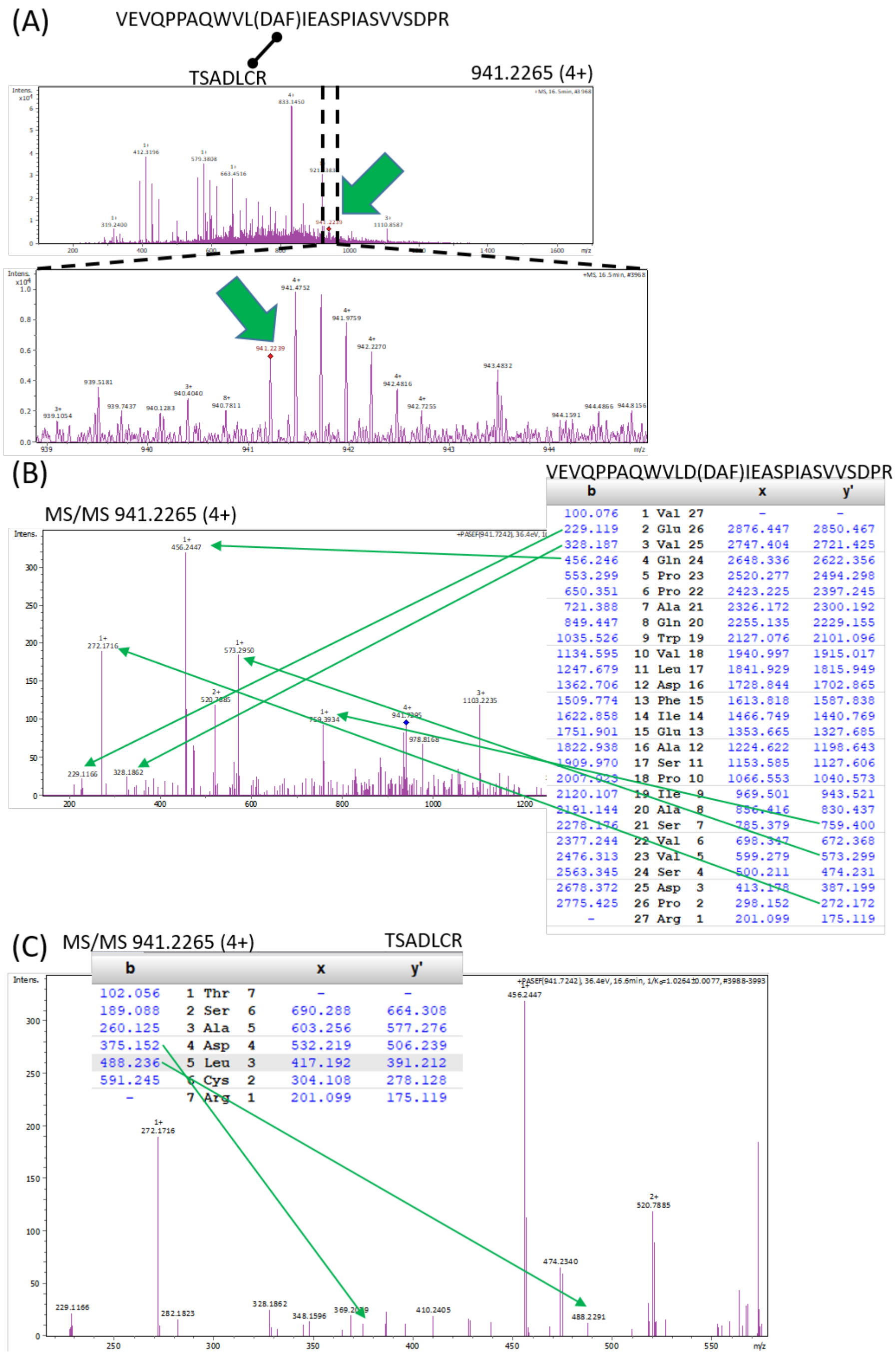

VEVQPPAQWVLD(DAF)IEASPIASVVSDPR

| b | | | | x | y' |
|---|---|---|---|---|---|
| 100.076 | 1 | Val | 27 | - | - |
| 229.119 | 2 | Glu | 26 | 2876.447 | 2850.467 |
| 328.187 | 3 | Val | 25 | 2747.404 | 2721.425 |
| 456.246 | 4 | Gln | 24 | 2648.336 | 2622.356 |
| 553.299 | 5 | Pro | 23 | 2520.277 | 2494.298 |
| 650.351 | 6 | Pro | 22 | 2423.225 | 2397.245 |
| 721.388 | 7 | Ala | 21 | 2326.172 | 2300.192 |
| 849.447 | 8 | Gln | 20 | 2255.135 | 2229.155 |
| 1035.526 | 9 | Trp | 19 | 2127.076 | 2101.096 |
| 1134.595 | 10 | Val | 18 | 1940.997 | 1915.017 |
| 1247.679 | 11 | Leu | 17 | 1841.929 | 1815.949 |
| 1362.706 | 12 | Asp | 16 | 1728.844 | 1702.865 |
| 1509.774 | 13 | Phe | 15 | 1613.818 | 1587.838 |
| 1622.858 | 14 | Ile | 14 | 1466.749 | 1440.769 |
| 1751.901 | 15 | Glu | 13 | 1353.665 | 1327.685 |
| 1822.938 | 16 | Ala | 12 | 1224.622 | 1198.643 |
| 1909.970 | 17 | Ser | 11 | 1153.585 | 1127.606 |
| 2007.023 | 18 | Pro | 10 | 1066.553 | 1040.573 |
| 2120.107 | 19 | Ile | 9 | 969.501 | 943.521 |
| 2191.144 | 20 | Ala | 8 | 856.416 | 830.437 |
| 2278.176 | 21 | Ser | 7 | 785.379 | 759.400 |
| 2377.244 | 22 | Val | 6 | 698.347 | 672.368 |
| 2476.313 | 23 | Val | 5 | 599.279 | 573.299 |
| 2563.345 | 24 | Ser | 4 | 500.211 | 474.231 |
| 2678.372 | 25 | Asp | 3 | 413.178 | 387.199 |
| 2775.425 | 26 | Pro | 2 | 298.152 | 272.172 |
| - | 27 | Arg | 1 | 201.099 | 175.119 |

TSADLCR

| b | | | | x | y' |
|---|---|---|---|---|---|
| 102.056 | 1 | Thr | 7 | - | - |
| 189.088 | 2 | Ser | 6 | 690.288 | 664.308 |
| 260.125 | 3 | Ala | 5 | 603.256 | 577.276 |
| 375.152 | 4 | Asp | 4 | 532.219 | 506.239 |
| 488.236 | 5 | Leu | 3 | 417.192 | 391.212 |
| 591.245 | 6 | Cys | 2 | 304.108 | 278.128 |
| - | 7 | Arg | 1 | 201.099 | 175.119 |

**Fig S14. Mass spectrometry analysis of EL222-L35DAF-V217CYS.** (A) Mass spectra confirming the presence of a peptide encompassing a cross-link between residues 35 and 217 (green arrow): 941.2265 (4+). The bottom panel shows an enlarged view of the spectral region delimited by the dashed lines. (B) Tandem mass spectra (MS/MS) of the cross-linked peptide focusing on the N-terminal fragment. (C) Same MS/MS as (B) but highlighting the C-terminal fragment. Green arrows indicate the peptide fragments confirmed experimentally (expected masses are listed in the tables). Note that these results confirm the existence of a cross-link in the sample but cannot be used to determine its overall abundance (cross-linking efficiency).

#### Note S14. Additional fiSPR experiments.

The cross-linked mutant EL222-L35DAF/V217C (XL-EL222) is compared to the wild-type EL222 (WT-EL222) under virtually identical experimental conditions. Both interactions are monitored in the dark and under *in situ* illumination at a wavelength of 450 nm, a light intensity of 3.3 mW/cm$^2$, and an illumination time of 260 s. Both WT- and XL-EL222 exhibit low binding to dsDNA in the dark, with sensor responses of 0.11 and 0.01 nm, respectively.

**Fig. S15A** shows the kinetic curves describing the interaction between XL-EL222 dimers and dsDNA immobilized on the SPR chip *via* biotin-streptavidin interactions. **Table S8** presents the kinetic association and dissociation rate constants ($k_a$ and $k_d$, respectively), and the equilibrium dissociation constant ($K_D$) of XL-EL222 dimers–dsDNA interaction, along with the $k_a$, $k_d$, and $K_D$ of WT-EL222 dimer–dsDNA interaction. The concentrations of both XL-EL222 and WT-EL222 dimers at equilibrium were determined from the initial concentrations of the corresponding monomers, assuming the same $K_D$ for the dimerization of both proteins i.e., $K_{D,D1}$ ~ 8 µM, $K_{D,D2}$ ~ 0.6 µM, as determined in our previous work (Ref. 50). The $k_a$ and $k_d$ of XL- and WT-EL222–dsDNA interaction differ by factors of ~2 and 3, respectively, resulting in a 5-fold lower affinity of XL-EL222 to dsDNA compared to WT-EL222.

**Fig. S15B** shows the dependence of the sensor response (readout at 300 s from EL222 injection) on the concentration of EL222 monomer (where EL222 indicates both XL-EL222, data from Fig. S15A, and WT-EL222 (the latter data was previously published in Ref. 50). The results are fitted using the Hill model (equation in **Fig. S15B**), which yields an exponent (n) equal or higher than 2 for both proteins interacting with dsDNA, suggesting that more than one monomer of XL- or WT-EL222 is captured by one dsDNA molecule on the sensor surface. The Hill-model fitting also indicates a ~5-fold lower affinity of XL-EL222 to dsDNA compared to WT-EL222, in agreement with the results in **Table S8**.

**Table. S10:** Rate and equilibrium constants of the interaction between XL-EL222 dimers and dsDNA, along with those of WT-EL222 dimer–dsDNA interaction (as previously determined in Ref. 50. The results are averaged from three measurements, and the associated error corresponds to the standard deviation of the measurements.

| | $k_a$ ($M^{-1}s^{-1}$) | $k_d$ ($s^{-1}$) | $K_D$ (µM) |
|---|---|---|---|
| **XL-EL222 / dsDNA** | $(3.3 \pm 0.9) \cdot 10^3$ | $(1.2 \pm 0.1) \cdot 10^{-2}$ | $3.7 \pm 0.9$ |
| **WT-EL222 / dsDNA** | $(6.5 \pm 1.3) \cdot 10^3$ | $(4.4 \pm 0.6) \cdot 10^{-3}$ | $0.7 \pm 0.2$ |

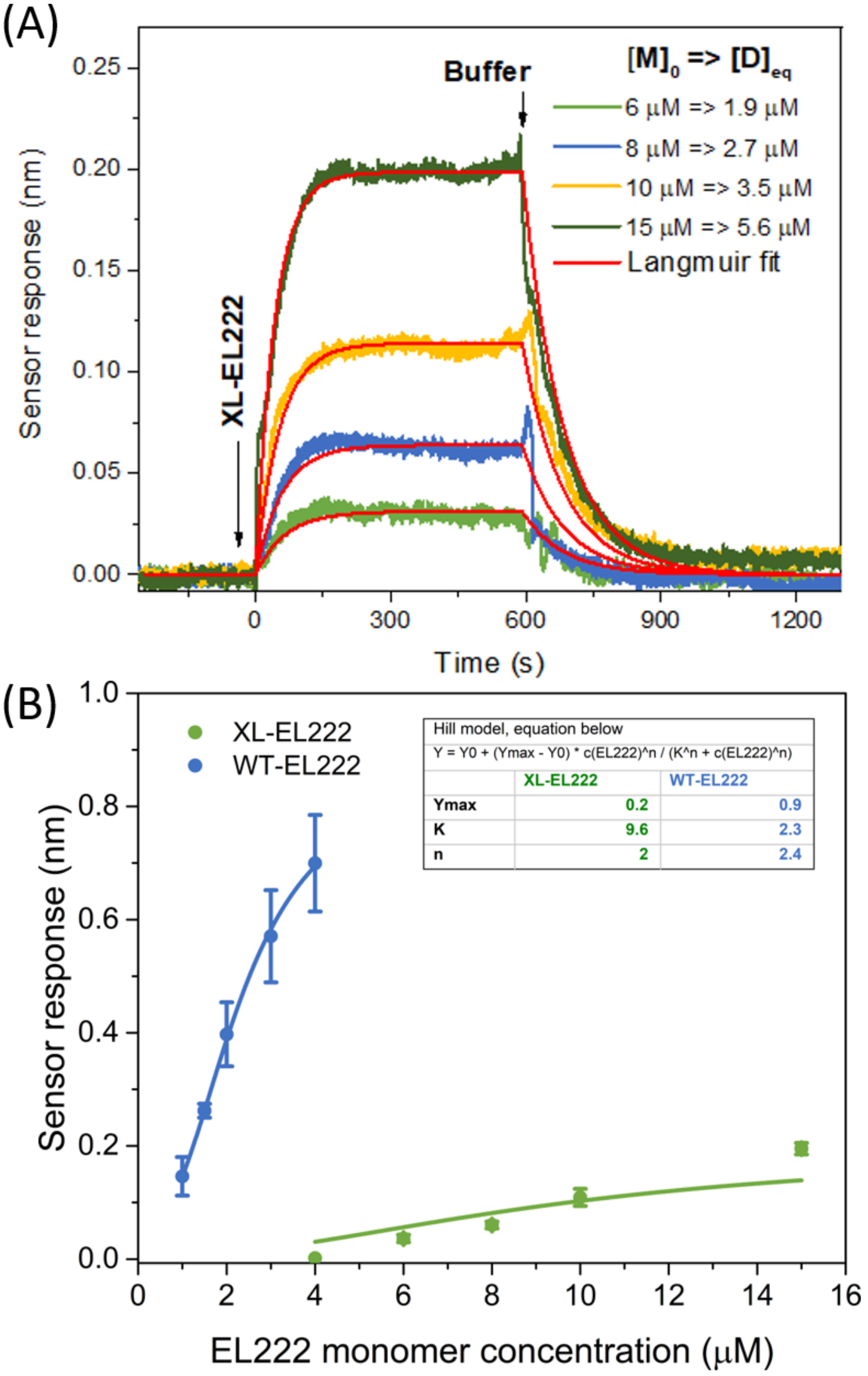

**Fig. S15. SPR characterization of the interaction between EL222 and DNA.** (A) Reference-compensated sensor response to different concentrations of XL-EL222 binding to dsDNA under *in situ* illumination. The red curves correspond to the least-squares fits of the sensor responses using a 1:1 Langmuir model, with $[D]_{eq}$ as the concentration of EL222 dimers at equilibrium. These measurements were repeated three times. (B) Reference-compensated sensor response to EL222 binding to dsDNA as a function of EL222 concentration (for both XL-EL222 and WT-EL222). The solid lines indicate the data fits using the Hill model (equation and parameters reported in the table inset).

#### Note S15. FRET analysis of EL222-DAF.

The fluorescence decay curves of donor-only (D, EL222-WT) and donor/acceptor (DA, EL222-I225DAF-tetrazine-AZDye594, which contains also EL222-I225DAF as the degree of labeling was estimated as 75%) were jointly analyzed (**Fig. S16A**). We used the maximum entropy method to convert the data from the time domain to the lifetime domain by the inverse Laplace transform as described in **Note S17**. For simplicity, we performed a tail fit starting at the top the decay i.e. the beginning of the curve, including the initial rise that is mostly influenced by the instrument response function (IRF) was not taken into account for the analysis. The lifetime distributions in the form of average dynamical content (*D*) as a function of lifetime ($\tau$) are shown in **Fig. S16B**. The distributions featured two main peaks. The most abundant peak centered at ~ 4 ns was assigned to the donor-only (labeled as P($\tau_D$) in **Fig. S16B**) sub-population, and the least abundant at ~ 1 ns was assigned to the donor/acceptor (labeled as P($\tau_{DA}$) in **Fig. S16B**) sub-population, based on the analysis of donor-only curves (which yielded only the component at ~4 ns). **Table S7** lists the amplitude-weighted mean lifetimes, $\langle\tau_x\rangle$ (where the subindex *i* refers to the D or DA distributions), which is the correct form of averaging[12,13].

The lifetime-based FRET efficiencies were subsequently calculated as:

$$E_{FRET} = 1 - \frac{\langle\tau_{DA}\rangle}{\langle\tau_D\rangle}$$

Finally, the donor-to-acceptor distance ($r_{DA}$) was calculated assuming a delta distribution (which is a reasonable assumption for folded proteins) and the Förster equation:

$$r_{DA} = R_O * \sqrt[6]{\frac{1}{E_{FRET}} - 1}$$

The Förster distance $R_0$, estimated as 4.5 nm, was calculated from:

$$R_O = 0.128 * (\kappa^2 \phi_D n^{-4} J)^{1/6}$$

The orientational factor $\kappa^2$ was assumed 2/3 (freely rotating dipoles), the donor quantum yield $\phi_D$ was set as 0.2 based on related LOV domains,[14,15] the refractive index $n$ was 1.33 and the spectral overlap integral $J$ was calculated from the emission of donor-only (EL222-WT) and the excitation of acceptor-only (EL222-I225DAF-tetrazine-AZDye594 lacking FMN).

**Table S11. Parameters derived from maximum entropy analysis of time-resolved fluorescence curves of EL222.** Average donor-only ($\langle\tau_D\rangle$) and donor/acceptor lifetimes ($\langle\tau_{DA}\rangle$), fractional amplitude of the donor/acceptor component ($f_\tau^{DA}$), FRET efficiencies ($E_{FRET}$) and donor-to-acceptor distances ($r_{DA}$) arising from maximum entropy analysis of time-resolved fluorescence datasets of EL222. Donor= FMN. Acceptor= AZDye594. The results are expressed as mean ± standard deviation of two independent experiments (except for the distance since the errors originate mainly from the incorrect estimation of $R_O$).

| **PARAMETER**<br><br>**CONDITION** | $\langle\boldsymbol{\tau_D}\rangle$<br>**(ns)** | $\langle\boldsymbol{\tau_{DA}}\rangle$<br>**(ns)** | $\boldsymbol{f_\tau^{DA}}$ (*) | $\boldsymbol{E_{FRET}}$ | $\boldsymbol{r_{DA}}$<br>**(nm)** |
|---|---|---|---|---|---|
| *DARK* | 3.48±0.11 | 0.83±0.03 | 0.342±0.005 | 0.757±0.002 | 3.7 |
| *LIT 250 s* | 3.47±0.15 | 0.92±0.03 | 0.30±0.02 | 0.732±0.003 | 3.8 |
| *LIT 40 s* | 3.57±0.01 | 0.97±0.18 | 0.20±0.04 | 0.73±0.05 | 3.8 |
| *LIT 20 s* | 3.66±0.05 | 1.05±0.15 | 0.17±0.05 | 0.71±0.04 | 3.9 |
| *LIT 10 s* | 3.92±0.17 | 0.99±0.18 | 0.15±0.01 | 0.75±0.03 | 3.8 |

* $f_\tau^{DA} + f_\tau^{D} = 1$

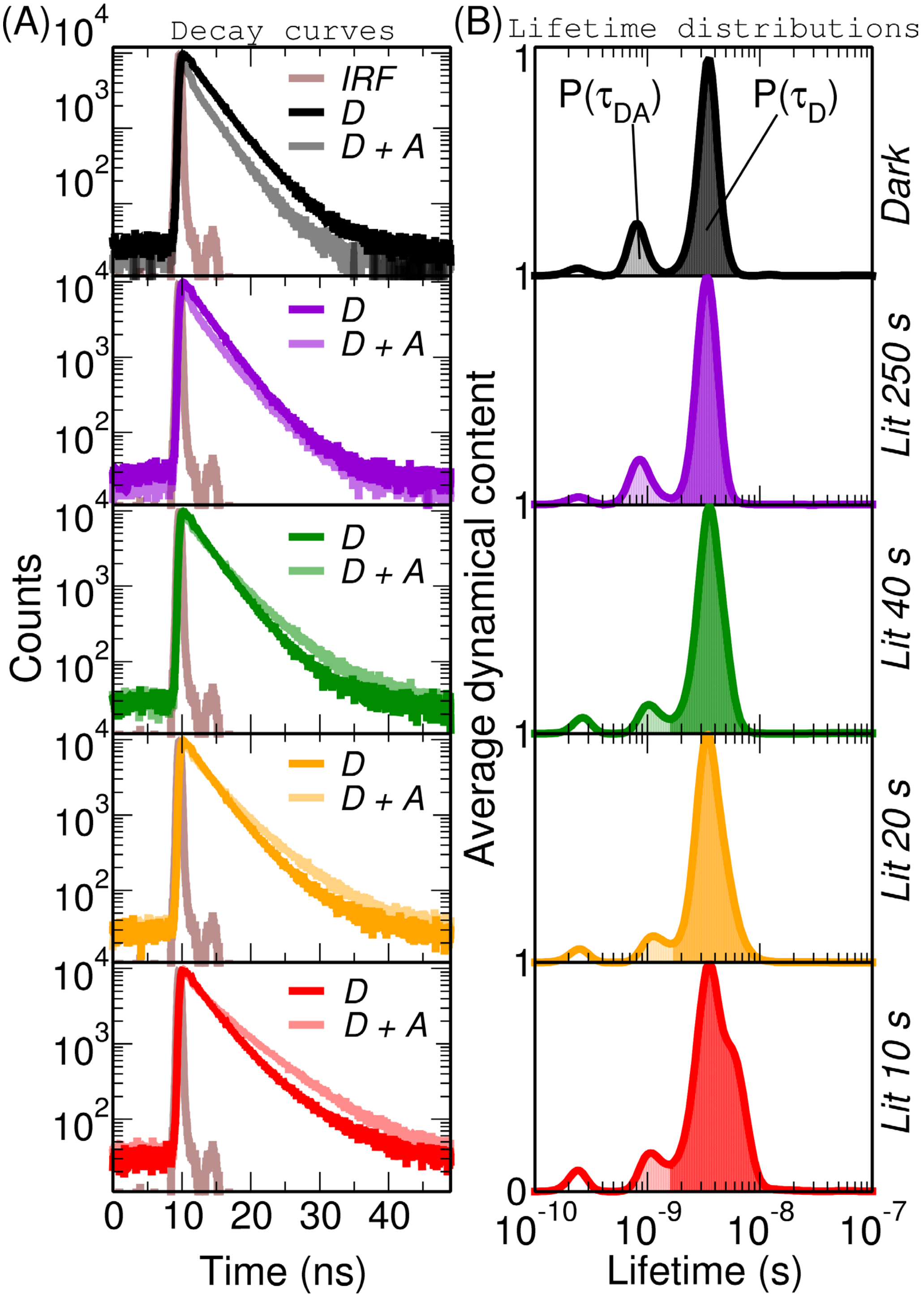

**Fig. S16. Lifetime distribution analysis of fluorescence decay curves of EL222 via the maximum entropy method.** (A) Fluorescence decay curves of donor-only (D, EL222-WT) and donor in the presence of the acceptor (D+A, EL222-I225DAF-tetrazined-AZDye594) measured with $\lambda_{excitation}$ = 450 nm, $\lambda_{emission}$ = 525 nm). IRF is the instrument response function. (B) Corresponding normalized average dynamical content as a function of lifetime. P($\tau_D$) and P($\tau_{DA}$) denote the lifetime distributions assigned to donor-only and donor-acceptor, respectively. The five experimental conditions (one dark and four lit) are shown on the right side.

#### Note S16. Steady-state infrared spectroscopy of EL222-CNF.

1 mM of three different EL222 CNF mutants were measured by infrared spectroscopy both in dark and light states. The difference spectra between two states revealed residues 35 and 151 respond to changes in the local environment. CNF35 senses the close (LOV-HTH interactions present)-open (LOV-HTH interactions absent) equilibrium of EL222, while CNF151 senses the folding-unfolding equilibrium of the linker between the LOV and HTH domains (Ref. 9). However, residue 225, present in DNA binding domain, did not experience a significant change in its microenvironment along the photocycle.

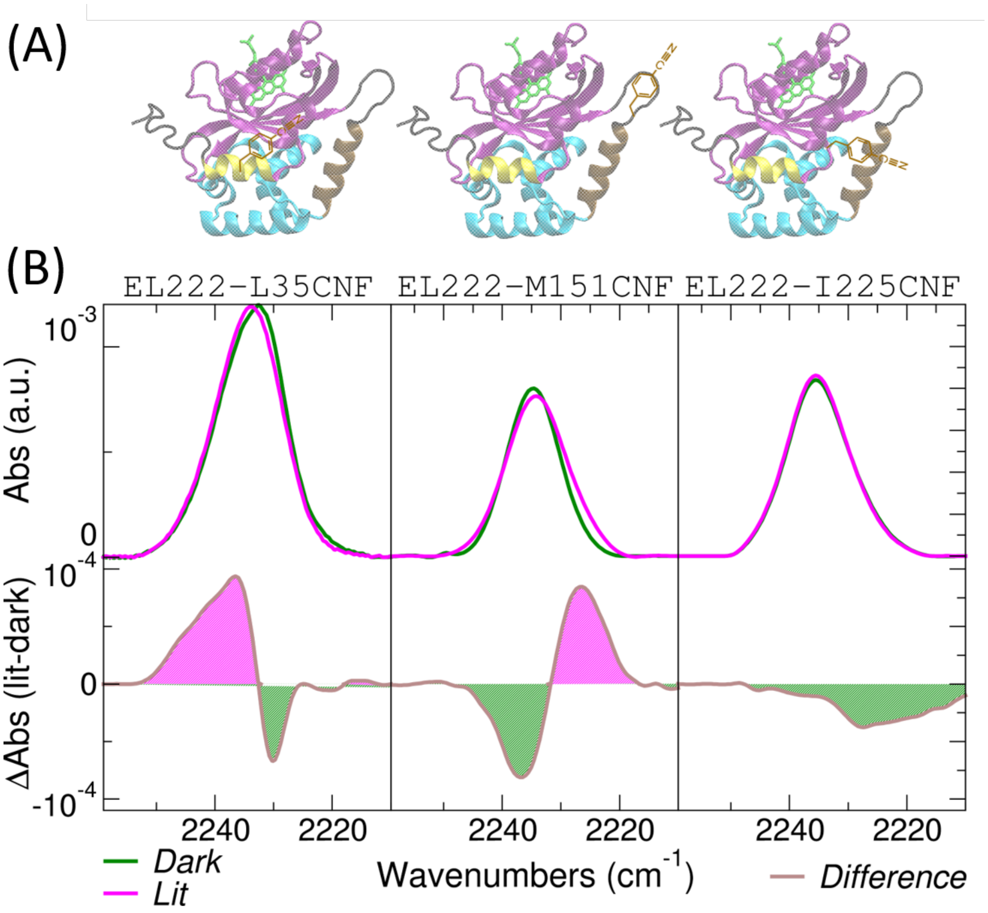

**Fig S17. Stationary Infrared spectroscopy of CNF-containing EL222 variants.** (A) Molecular models of EL222 containing CNF at positions 35 (left), 151 (middle), and 225 (right). (B) Steady-state infrared spectra of the corresponding $EL222^{CNF}$ mutants in the dark and under continuous blue light illumination ($\lambda$ = 455 nm, 25 mW). Absolute spectra are shown in the top panels and difference (light-minus-dark) spectra are shown in the bottom panels. The spectral data have been replotted from Ref. 9 ([Zenodo Data set] https://doi.org/10.5281/zenodo.7086623)

#### Note S17. Additional FSRS of DAF and EL222-DAF.

Raman spectra of 10 mM DAF dissolved in 100 % DMSO showed two intense peaks at 2195 $cm^{-1}$ and 2128 $cm^{-1}$. We focused on the first peak, which can be unambiguously assigned to the vibration of the dyine moiety. The band at 2195 $cm^{-1}$ showed linear dependency on the energy of the Raman pulse ($\lambda_{Raman\ pump}$ = 800 nm) up to 10 µJ (**Fig. S18A**).

To test the sensitivity of DAF to the protein microenvironment, 0.9 mM EL222-L35DAF was mixed with 4 M of urea, a denaturing agent, to globally unfold the protein. A ~ 1 $cm^{-1}$ clear Raman shift was observed between non-denaturing and denaturing conditions (**Fig. 18B**). This result is significantly different from the partial (local) EL222 unfolding induced by illumination, which causes no clear shift in the position of the Raman position (only a change in the area).

To compare the response of DAF and ETF, we measured the stimulated Raman spectra of 1.5 mM of EL222-M151ETF under distinct illumination conditions. The observed light-minus-dark difference spectrum of EL222-M151ETF (**Fig. S18C** top) shows a similar pattern to that of EL222-M151DAF (**Fig.6B**) although with reduced intensity. The lower signal-to-noise provided by ETF is clearly manifested in the time-resolved spectra (**Fig. S18C** bottom).

Not all residue positions act as reporters of photoinduced conformational changes in EL222. For instance, of the three chosen positions, EL222-I225DAF features modest changes in the steady-state Raman difference spectrum (**Fig. S18D** top) and virtually no changes in the time-resolved Raman spectra (**Fig. S18D** bottom).

We also compared the Raman signals in the fingerprint region of EL222, particularly in the 1600 $cm^{-1}$-to-1700 $cm^{-1}$ region, which is mainly reflecting the C=O vibrations arising from the protein backbone (Amide I band). The absolute and difference Raman spectra (**Fig. S18E**) are similar for all three mutants (EL222-L35DAF, EL222-M151DAF, and EL222-I225DAF) showing a positive-negative-positive feature at ~1680 $cm^{-1}$- ~1640 $cm^{-1}$-~1600 $cm^{-1}$, respectively. These results contrast with the response of DAF in the alkyne stretching region (~2200 $cm^{-1}$), which is clearly residue-specific.

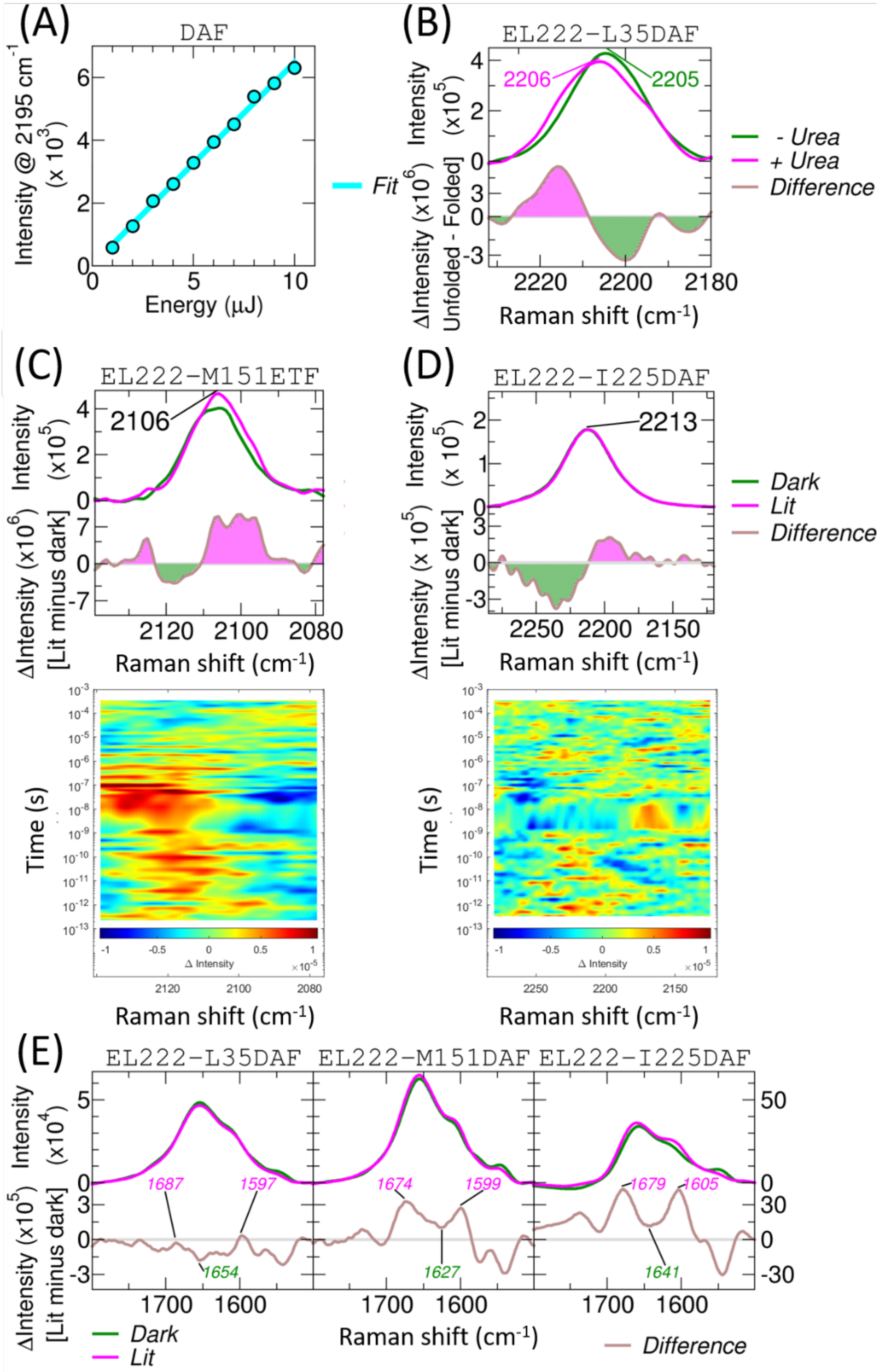

**Figure S18. FSRS of DAF and EL222-DAF.** (A) Energy dependence of stimulated Raman signal of DAF in DMSO. (B) Absolute and difference (unfolded-minus-folded) stimulated Raman spectra of EL222-L35DAF in the presence and absence of 4 M urea. (C) Absolute and difference (light-minus-dark) stimulated Raman spectra of EL222-M151EFT (*top*) and the corresponding time-resolved FSRS upon 480 nm excitation (*bottom*). (D) Absolute and difference (light-minus-dark) stimulated Raman spectra of EL222-I225DAF (*top*) and the corresponding time-resolved FSRS upon 480 nm excitation (*bottom*). (E) Absolute and difference (light-minus-dark) stimulated Raman spectra of EL222-L35DAF (left), EL222-M151DAF (middle), EL222-I225DAF (right) in the 1500-1750 cm-1 spectral "fingerprint" region. $\lambda_{\text{Raman pump}}$ = 800 nm.

#### Note S18. Analysis of time-resolved Raman datasets.

Transient femtosecond-stimulated Raman spectra from femtoseconds to ~0.5 ms in the spectral region from 2100 $cm^{-1}$ to 2300 $cm^{-1}$ (vibration of the diyne moiety of DAF) were subjected to lifetime distribution analysis (LDA) by the maximum entropy method as previously described.[16-18] For LDA, the lifetimes were fixed and logarithmically distributed from $10^{-14}$ s to $10^{-3}$ s with 40 points per decade. The wavelength- and time-dependent amplitudes were the only fitting parameters. The L-curve criterion was used to select the optimal regularization parameter for the inverse Laplace transform (conversion of time domain data to lifetime domain). From the 2D lifetime density maps, the lifetime-dependent average dynamical content *D* (**Fig. S15A**) was computed as the square-root of the summation over the squared amplitudes.[19,20] The peak centers of the *D* lifetime distribution were retrieved (**Fig. S15B**). The decay associated difference spectra (DADS, **Fig. S15B**) were extracted from the 2D lifetime distribution upon integration of the transient spectra in the corresponding time range.

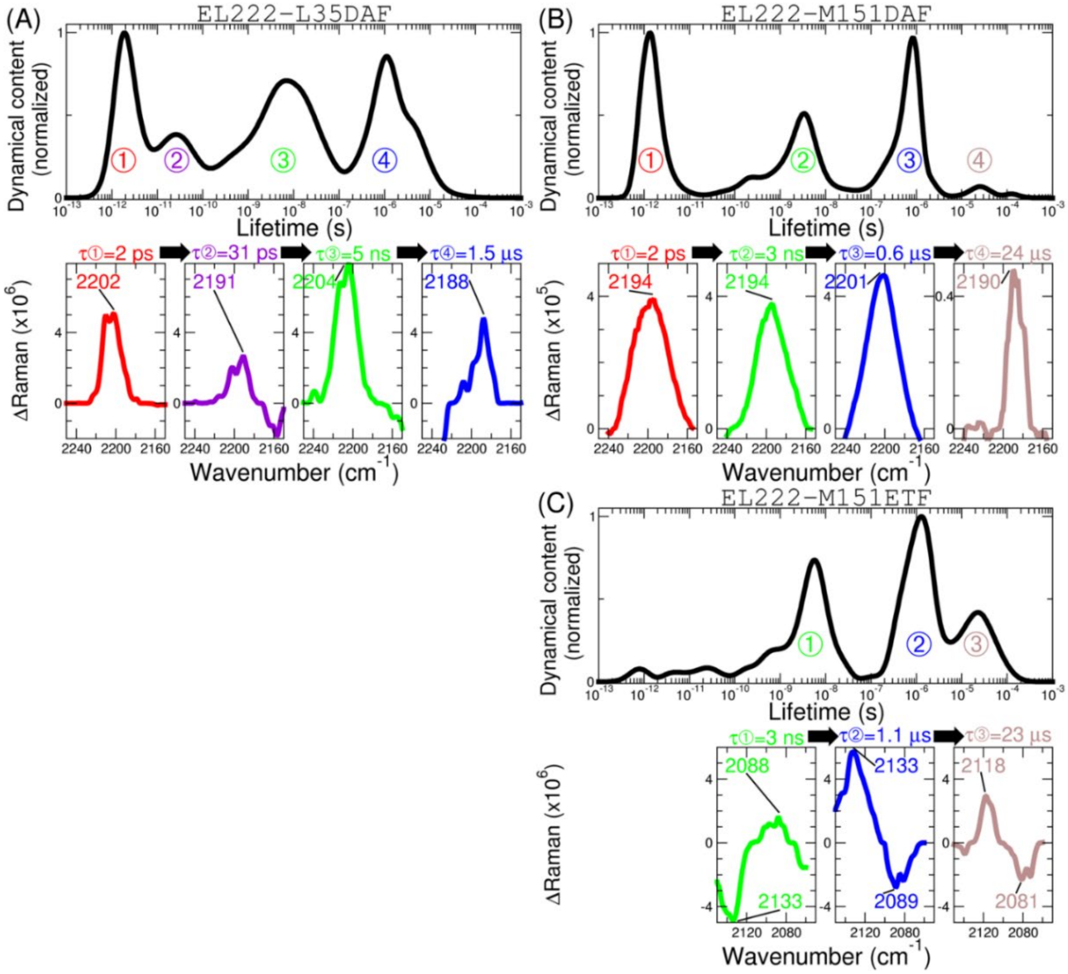

**Fig. S19. Lifetime distribution analysis of time-resolved FSRS datasets via the maximum entropy method.** (A) Average dynamical content as a function of lifetime. (B) Integrated spectra equivalent to decay-associated difference spectra (DADS). (*Left*) EL222-L35DAF, (*right*) EL222-M151DAF. The Raman spectra were analyzed between 2100 and 2300 $cm^{-1}$ (C≡C vibration of DAF).

#### Note S19. Time-resolved absorption of EL222-DAF.

Photoactivation of two EL222 DAF mutants were also studied by transient absorption spectroscopy (**Fig. S20A**) in a custom-built set-up.

The central laser system consisted of two independent 1 kHz laser amplifiers, both seeded by a single Ti:sapphire oscillator. The output from one amplifier (Femtopower, Spectra Physics) was directed into an argon-filled hollow-core fiber (Savanna, Ultrafast Innovations), generating a white-light supercontinuum spanning a spectral range of 300 to 1100 nm and covering 4 picoseconds. This light served directly as the probe beam. The pump beam, centered at 477 nm and delivering ~100 nJ per pulse, was produced via a Noncollinear Optical Parametric Amplifier (NOPA; TOPAS, Light Conversion), driven by the second amplifier (Solstice, Spectra Physics). The timing between the pump and probe beams was scanned up to 100 μs with femtosecond precision, achieved through a combination of an optomechanical delay line and precise electronic synchronization between both laser amplifiers. The polarization of the beams was configured at magic angle. Both beams were modulated using a pair of optomechanical choppers, enabling rapid detection of the transient absorption signal, while correcting for pump scattering and detector dark noise on a shot-to-shot basis. The probe transmitted through the sample was spectrally dispersed using a custom dual-channel prism spectrometer and captured by a CCD camera (Entwicklungsbüro Stresing). The second channel of the spectrometer provided reference measurements for correcting probe fluctuations, following the methodology outlined in [21]. The sample was contained within a 1-mm thick optical cell and was automatically repositioned by 2D linear stage (SmarAct) in the transverse plane to ensure fresh sample material remained in the focal area.

We analyzed the data by lifetime distribution analysis (see details in **Note S17**) and obtained the average dynamical content (**Fig. S20B**) and the decay-associated difference spectra (**Fig. S20C**). The detailed interpretation is explained with **Fig.6** in the main text.

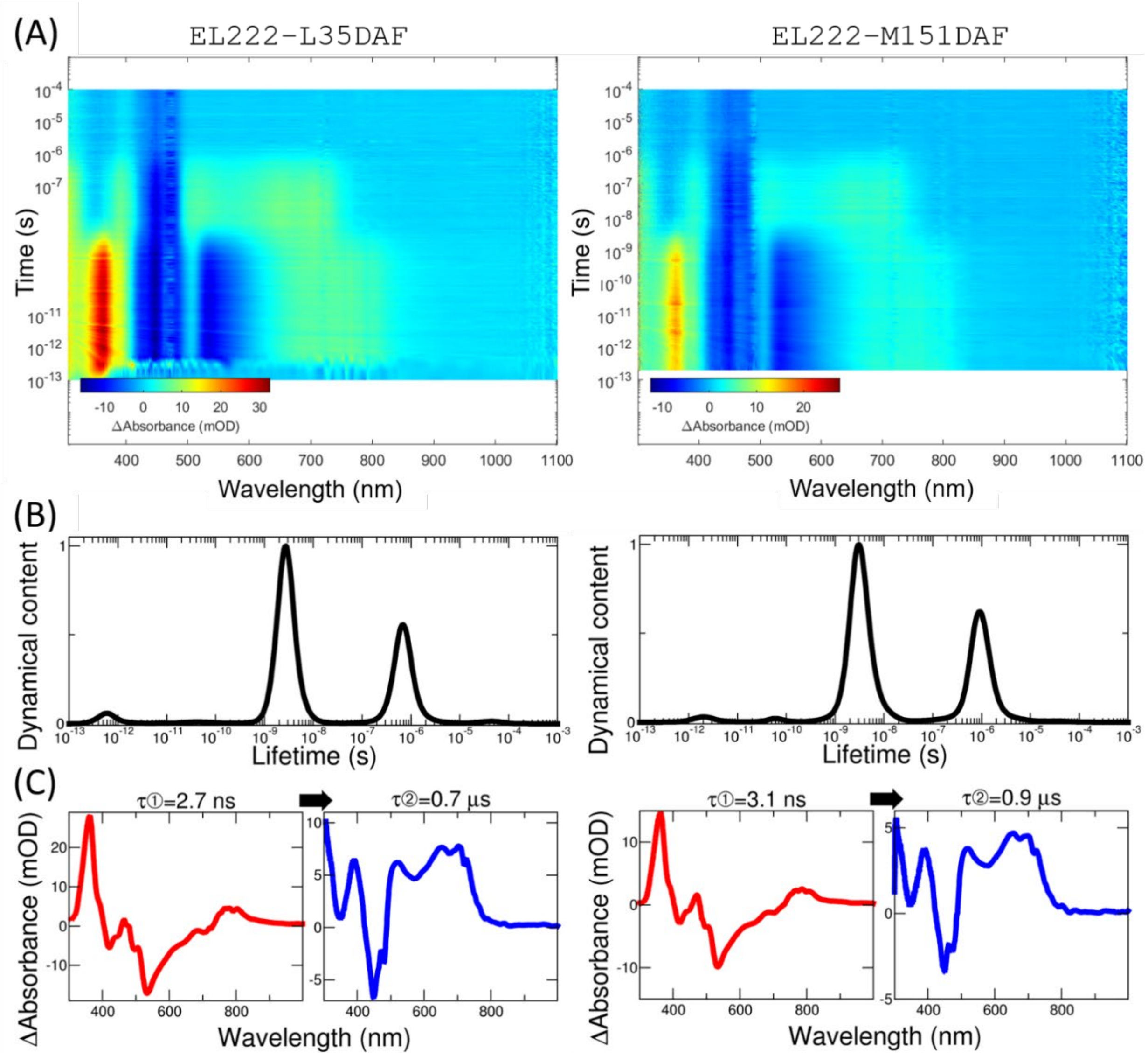

**Fig. S20. Lifetime distribution analysis of transient absorption datasets via the maximum entropy method.** (A) 2D contour plots of differential absorption as a function of pump-probe time delays and wavelength of EL222-L35DAF (*left*) and EL222-M151DAF (*right*) mutants (B) Average dynamical content as a function of lifetime. (C) Integrated spectra equivalent to decay-associated difference spectra (DADS). EL222-L35DAF are the same samples as those shown in **Fig. 8** and **Fig. S14**. The UV/visible spectra were analyzed between 300 and 1000 nm, the region corresponding to the absorption of the FMN chromophore.

#### Note S20. Raman microspectroscopy of DAF.

Solutions of DAF at different concentrations were prepared by mixing a 200 mM stock in methanol with distilled water. Suspensions of *E. coli* cells encoding DAF (in proteins EL222 and MBP) and their negative controls were measured. The first batch was measured directly in the LB medium. The second batch was centrifuged, LB medium discarded, and resuspended in PBS. The suspension was put into a chamber made of microscope slide, spacer, and cover slip (**Fig. S21A**). In all cases, the laser was focused a few microns below the cover slip (thus a few microns into the sample volume).

Measurements of the DAF non-canonical amino acid as a dried sample on a $CaF_2$ slide resulted only in fluorescence background overshadowing any Raman signal. However, when DAF is diluted in water and measured in a sealed chamber, the expected peak at approx. 2210 $cm^{-1}$ was clearly visible after spectra processing (**Fig. S21B**). The higher concentration of DAF could be noticed also by the proportionally increasing fluorescence. The limit-of-detection of our set-up is estimated as ~0.5 mM of DAF.

Raman spectra from *E. coli* cells required stronger data processing than when diluted DAF was measured. This is due to the higher background signal from the cells, medium and buffer. The samples measured in LB medium resulted in severe fluorescence and even with the data processing, no signal from DAF was acquired. Spectra obtained from cells resuspended in PBS were as follows: both EL222 and MBP encoding DAF showed stronger fluorescence, so the resulting spectra are noisier as opposed to the negative control. EL222 samples did not show any clear peak in the “transparent window” region (**Fig. S21C**). However, only the MBP sample led to the detection of the DAF peak at approx. 2205 $cm^{-1}$ (**Fig. S21D**). Taken into account the limit-of-detection, these results suggest that the intracellular concentration of EL222 is below 0.5 mM.

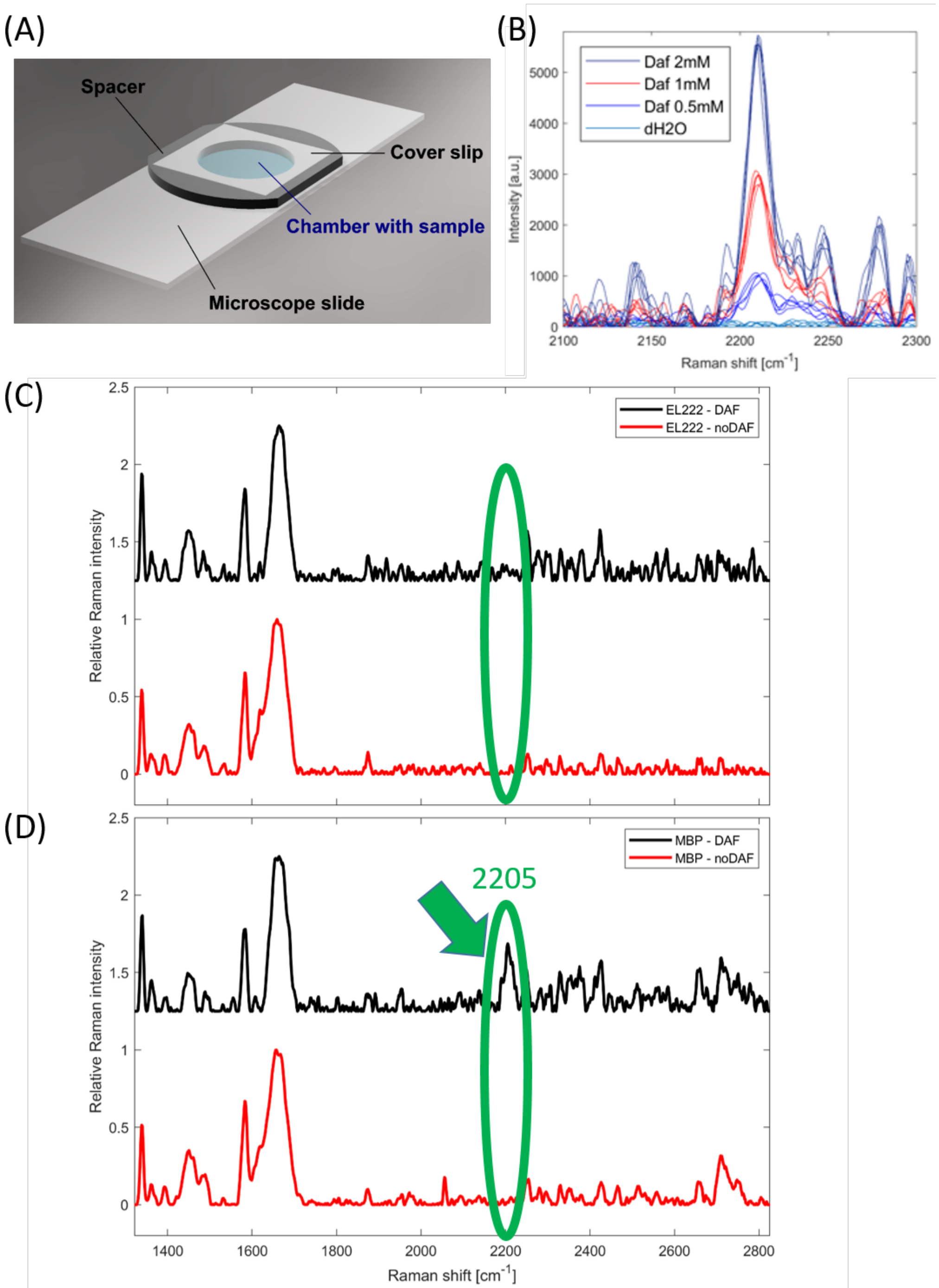

**Fig. S21. Raman microspectroscopy of DAF and DAF-containing proteins.** (A) Raman setup. Solutions or cell suspensions were put into a chamber made of microscope slide, spacer, and cover slip. (B) Raman spectra of DAF solutions at different concentrations in methanol:water mixtures. (C) Raman spectra of suspensions of *E. coli* cells in PBS buffer expressing EL222-M151DAF and the negative control without added DAF. (D) Raman spectra of suspensions of *E. coli* cells in PBS buffer expressing MBP-V38DAF and the negative control without added DAF.

#### SI references.

1 Ungeheuer, F. & Furstner, A. Concise Total Synthesis of Ivorenolide B. *Chemistry* **21**, 11387-11392, doi:10.1002/chem.201501765 (2015).

2 Hamissa, M. F. *et al.* Neutral and charged forms of inubosin B in aqueous solutions at different pH and on the surface of Ag nanoparticles. *Journal of Molecular Structure* **1250**, doi:10.1016/j.molstruc.2021.131828 (2022).

3 Shen, T. & Liu, X. Unveiling the photophysical mechanistic mysteries of tetrazine-functionalized fluorogenic labels. *Chem Sci* **16**, 4595-4613, doi:10.1039/d4sc07018f (2025).

4 Devaraj, N. K., Hilderbrand, S., Upadhyay, R., Mazitschek, R. & Weissleder, R. Bioorthogonal Turn-On Probes for Imaging Small Molecules inside Living Cells. *Angewandte Chemie International Edition* **49**, 2869-2872, doi:10.1002/anie.200906120 (2010).

5 Lang, K. *et al.* Genetic Encoding of bicyclononynes and trans-cyclooctenes for site-specific protein labeling in vitro and in live mammalian cells via rapid fluorogenic Diels-Alder reactions. *J Am Chem Soc* **134**, 10317-10320, doi:10.1021/ja302832g (2012).

6 Blasiak, B., Ritchie, A. W., Webb, L. J. & Cho, M. Vibrational solvatochromism of nitrile infrared probes: beyond the vibrational Stark dipole approach. *Phys Chem Chem Phys* **18**, 18094-18111, doi:10.1039/c6cp01578f (2016).

7 Weaver, J. B., Kozuch, J., Kirsh, J. M. & Boxer, S. G. Nitrile Infrared Intensities Characterize Electric Fields and Hydrogen Bonding in Protic, Aprotic, and Protein Environments. *Journal of the American Chemical Society* **144**, 7562-7567, doi:10.1021/jacs.2c00675 (2022).

8 Romei, M. G. *et al.* Frequency Changes in Terminal Alkynes Provide Strong, Sensitive, and Solvatochromic Raman Probes of Biochemical Environments. *The Journal of Physical Chemistry B* **127**, 85-94, doi:10.1021/acs.jpcb.2c06176 (2022).

9 Beránek, V., Willis, J. C. W. & Chin, J. W. An Evolved Methanomethylophilus alvus Pyrrolysyl-tRNA Synthetase/tRNA Pair Is Highly Active and Orthogonal in Mammalian Cells. *Biochemistry* **58**, 387-390, doi:10.1021/acs.biochem.8b00808 (2018).

10 Avila-Crump, S. *et al.* Generating Efficient Methanomethylophilus alvus Pyrrolysyl-tRNA Synthetases for Structurally Diverse Non-Canonical Amino Acids. *ACS Chemical Biology* **17**, 3458-3469, doi:10.1021/acschembio.2c00639 (2022).

11 Alexander, N. D. *et al.* Selecting aminoacyl-tRNA synthetase/tRNA pairs for efficient genetic encoding of noncanonical amino acids into proteins. *Nat Protoc* **21**, 1374-1428, doi:10.1038/s41596-025-01241-w (2026).

12 Lakowicz, J. R. *Principles of Fluorescence Spectroscopy*. (Springer, 2006).

13 Becker, W. *The bh TCSPC handbook*. (Becker & Hickl GmbH, 2026).

14 Penzkofer, A., Endres, L., Schiereis, T. & Hegemann, P. Yield of photo-adduct formation of LOV domains from Chlamydomonas reinhardtii by picosecond laser excitation. *Chemical Physics* **316**, 185-194, doi:https://doi.org/10.1016/j.chemphys.2005.05.016 (2005).

15 Schüttrigkeit, T. A., Kompa, C. K., Salomon, M., Rüdiger, W. & Michel-Beyerle, M. E. Primary photophysics of the FMN binding LOV2 domain of the plant blue light receptor phototropin of Avena sativa. *Chemical Physics* **294**, 501-508, doi:https://doi.org/10.1016/S0301-0104(03)00390-2 (2003).

16 Lorenz-Fonfria, V. A. & Kandori, H. Bayesian maximum entropy (two-dimensional) lifetime distribution reconstruction from time-resolved spectroscopic data. *Appl Spectrosc* **61**, 428-443, doi:10.1366/000370207780466172 (2007).

17 Lorenz-Fonfria, V. A. & Kandori, H. Transformation of time-resolved spectra to lifetime-resolved spectra by maximum entropy inversion of the laplace transform. *Appl Spectrosc* **60**, 407-417, doi:10.1366/000370206776593654 (2006).

18 Lorenz-Fonfria, V. A. & Kandori, H. Practical aspects of the maximum entropy inversion of the laplace transform for the quantitative analysis of multi-exponential data. *Appl Spectrosc* **61**, 74-84, doi:10.1366/000370207779701460 (2007).

19 Stock, G. & Hamm, P. A non-equilibrium approach to allosteric communication. *Philos Trans R Soc Lond B Biol Sci* **373**, doi:10.1098/rstb.2017.0187 (2018).

20 Bozovic, O. *et al.* Real-time observation of ligand-induced allosteric transitions in a PDZ domain. *Proc Natl Acad Sci U S A* **117**, 26031-26039, doi:10.1073/pnas.2012999117 (2020).

21 Feng, Y., Vinogradov, I. & Ge, N. H. General noise suppression scheme with reference detection in heterodyne nonlinear spectroscopy. *Opt Express* **25**, 26262-26279, doi:10.1364/OE.25.026262 (2017).
